# Neural Correlates of Cognition in Primary Visual versus Neighboring Posterior Cortices during Visual Evidence-Accumulation-based Navigation

**DOI:** 10.1101/568766

**Authors:** Sue Ann Koay, Stephan Y. Thiberge, Carlos D. Brody, David W. Tank

## Abstract

Studies of perceptual decision-making have often assumed that the main role of sensory cortices is to provide sensory input to downstream processes that accumulate and drive behavioral decisions. We performed a systematic comparison of neural activity in primary visual (V1) to secondary visual and retrosplenial cortices, as mice performed a task where they should accumulate pulsatile visual cues through time to inform a navigational decision. Even in V1, only a small fraction of neurons had sensory-like responses to cues. Instead, in all areas neurons were sequentially active, and contained information ranging from sensory to cognitive, including cue timings, evidence, place/time, decision and reward outcome. Per-cue sensory responses were amplitude-modulated by various cognitive quantities, notably accumulated evidence. This inspired a multiplicative feedback-loop circuit hypothesis that proposes a more intricate role of sensory areas in the accumulation process, and furthermore explains a surprising observation that perceptual discrimination deviates from Weber-Fechner Law.

**Highlights / eTOC Blurb:**

- Mice made navigational decisions based on accumulating pulsatile visual cues
- The bulk of neural activity in visual cortices was sequential and beyond-sensory
- Accumulated pulse-counts modulated sensory (cue) responses, suggesting feedback
- A feedback-loop neural circuit explains behavioral deviations from Weber’s Law

In a task where navigation was informed by accumulated pulsatile visual evidence, neural activity in visual cortices predominantly coded for cognitive variables across multiple timescales, including outside of a visual processing context. Even sensory responses to visual pulses were amplitude-modulated by accumulated pulse counts and other variables, inspiring a multiplicative feedback-loop circuit hypothesis that in turn explained behavioral deviations from Weber-Fechner Law.

## Introduction

As sensory information about the world is often noisy and/or ambiguous, an evidence accumulation process is thought to be fundamental to perceptual decision-making, and requires working memory for maintenance and updating of the accumulated information. Neural circuits that perform this are incompletely known, but canonically hypothesized to involve multiple stages—sensory detection, accumulation, categorization, action selection—chained together in a predominantly feedforward manner (Gold and Shadlen 2007; Brody and Hanks 2016; Caballero, Humphries, and Gurney 2018). A substantial body of work seeks possible mappings of these conceptual stages to brain regions. In the neocortex, sensory areas are obvious candidates for the earliest (detection) stage, but have received relatively little attention in regards to other possible contributions.

The BRAIN COGS collaboration (“BRAIN Circuits Of coGnitive Systems,” n.d.) aims to understand the neural underpinnings of such decision-making behaviors from a whole-brain perspective, using the highly tractable mouse model system for which we have developed a navigation-based visual evidence accumulation task (“Accumulating-Towers” task, see (Lucas Pinto et al. 2018)). In the present work, we performed a thorough characterization of neural population activity in layers 2/3 and 5 of six early cortical areas, including the primary visual cortex (V1), secondary visual areas, and retrosplenial cortex. Unexpectedly, only a small part of neural activity in the visual areas was correlated with the momentary visual stimulus. Instead we observed prevalent coding of long timescale and cognitive/internal information, with differences only in degree compared to the retrosplenial cortex, a region thought to be involved in episodic memory, learning, and navigation (Mitchell et al. 2018; Vann, Aggleton, and Maguire 2009). Our findings for the visual cortices also had similarities to related studies of the posterior parietal cortex (PPC), which is thought to be involved in sensorimotor transformations, navigation and decision-making (Lyamzin and Benucci 2018).

We discovered the predominant form of neural dynamics in all areas and layers to be the sequential activation of cells. This resembles place cells in the hippocampus (Moser, Kropff, and Moser 2008), and place preferences have been reported by others in V1 (Saleem et al. 2018). However, most cells were preferentially active on either right- or left-choice trials, a phenomenon previously reported in the PPC (Harvey, Coen, and Tank 2012; Morcos and Harvey 2016). In fact, in all areas the activity levels of sequentially active cells contained further information about the accumulated evidence, behavioral choice, and reward outcome, including all these quantities from the past trial. These phenomena continued throughout the inter-trial-interval, despite the absence of a visual context. Even in V1, only a small fraction (5-10%) of all active cells exhibited sensory-like responses that were time-locked to individual visual cues. Remarkably, in all areas the per-cue amplitudes of these responses were also modulated by evidence, place/time, choice, and reward outcome.

Our findings of decision-related, non-sensory responses in sensory areas, as well as non-sensory modulations of sensory responses, invite the revisiting of two broad strokes of the canonical picture of decision-making based on accumulated evidence: the feedforward nature of the first stages, and the functional modularity of the involved brain regions (Siegel, Buschman, and Miller 2015; Michelson, Pillow, and Seidemann 2017). Top-down feedback is a candidate explanation for choice-related effects in sensory responses (Britten et al. 1996; Romo et al. 2003; Nienborg and Cumming 2009; Yang et al. 2016; Bondy, Haefner, and Cumming 2018; Wimmer et al. 2015; Haefner, Berkes, and Fiser 2016). Inspired by our neural observations that the amplitudes of sensory responses depended on accumulated counts, we proposed a multiplicative feedback-loop circuit model where accumulator feedback acts as a dynamic gain on the sensory input. Interestingly, this model could explain an unexpected psychophysical effect in pulse-based evidence accumulation tasks (Scott et al. 2015; Lucas Pinto et al. 2018), where the perceptual accuracy of animals deviated from the Weber-Fechner Law (Fechner 1966; Gallistel and Gelman 2000). For a fixed ratio of right-to-left counts, the Weber-Fechner Law states that discrimination of the difference should not depend on total counts. However, we found that the psychophysical data only followed this law at high total counts, with a transition between accuracy regimes that was best predicted by the feedback-loop model compared to models without feedback. Our results thus suggest that a behaviorally important feature of the accumulation process may be a gradual amplification of the per-pulse sensory responses, as reflected in neural activity in as early as V1.

## Results

We used cellular-resolution two-photon imaging to record from six posterior cortical regions of 11 mice trained in the Accumulating-Towers task (Fig. 1A-C). These mice were from transgenic lines that express the calcium-sensitive fluorescent indicator GCaMP6f in cortical excitatory neurons (Methods), and prior to behavioral training underwent surgical implantation of an optical cranial window centered over either the right or left parietal cortex. The mice then participated in previously detailed behavioral shaping (Lucas Pinto et al. 2018) and neural imaging procedures as summarized below.

**Figure 1.**
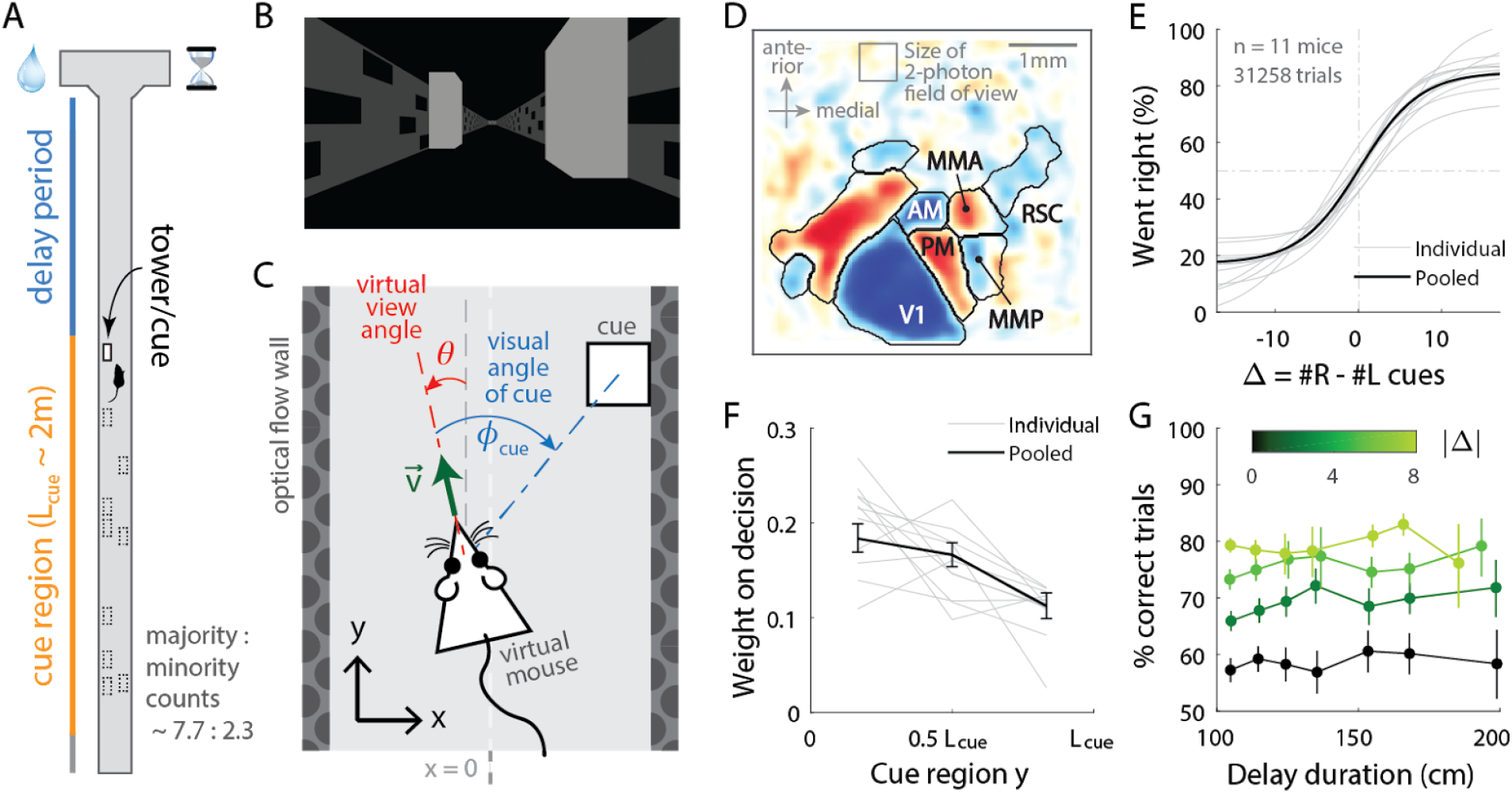
Two-photon calcium imaging of multiple cortical areas during a navigation-based evidence accumulation task. **(A)** Layout of the virtual T-maze in an example left-rewarded trial. **(B)** Example snapshot of the cue region corridor from a mouse’s point of view when facing straight down the maze. Two cues on the right and left sides can be seen, closer and further from the mouse in that order. **(C)** Illustration of the virtual viewing angle θ. The visual angle **φ**_*cue*_ of a given cue is measured relative to θ and to the center of the cue. The y spatial coordinate points straight down the stem of the maze, and the x coordinate is transverse. 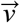 is the velocity of the mouse in the virtual world. **(D)** Average visual field sign map (*n* = 5 mice) and visual area boundaries, with all recorded areas labeled. The visual field sign is −1 (dark blue) where the cortical layout is a mirror image and +1 (dark red) where it follows a non-inverted layout of the physical world. **(E)** Sigmoid curve fits to psychometric data for how frequently mice turned right for a given difference in right vs. left cue counts, Δ #*R*−#*L*. **(F)** Logistic regression weights for predicting the mice’s choice given the evidence Δ in three spatial bins of the cue region. Error bars: 95% C.I. across bootstrap experiments. **(G)** Performance vs. effective duration of the delay period, which is the distance from the last cue to the end of the T-maze stem. Data were pooled over all #R + #L for statistical power, but analyses that account for this yield the same result (Lucas Pinto et al. 2018). Error bars: 95% C.I.

Mice were trained in a head-fixed virtual reality system (Dombeck et al. 2010) to navigate in a T-maze. As they ran down the stem of the maze, a series of transient (200ms), randomly located tower-shaped cues (Fig. 1B) appeared along the right and left walls of the cue region corridor (length *L_cue_* ≈ 200 cm; see Methods), followed by a delay region where no cues appeared. The locations of cues were drawn randomly per trial, with Poisson-distributed mean counts of 7.7 on the majority and 2.3 on the minority side, and mice were rewarded for turning down the arm corresponding to the side with more cues. As mice control the virtual viewing angle θ, cues could appear at a variety of angles **φ**_*cue*_ (Fig. 1C). We accounted for this in all relevant data analyses, as well as conducted control experiments in which θ was restricted to be exactly zero from the beginning of the trial up to midway in the delay period (referred to as θ-controlled experiments; see Methods). In agreement with previous work (Lucas Pinto et al. 2018), all mice in this study exhibited characteristic psychometric curves and utilized multiple pieces of evidence to make decisions, with a moderate primacy effect (Fig. 1E-F). For fixed total numbers of cues on the right (#R) and left (#L) sides, there was no degradation of performance with increasing effective length of the delay period (Fig. 1G). This is compatible with a negligible effect of memory leakage with time, as observed also in rats performing evidence-accumulation tasks (Brunton, Botvinick, and Brody 2013; Scott et al. 2015).

For each mouse, we first identified the locations of the visual areas (Fig. 1D; Methods) using one-photon widefield imaging and a retinotopic visual stimulation protocol (Zhuang et al. 2017). Then, while the mice performed the task, we used two-photon imaging to record from 500μ*m* × 500μ*m* fields of view in either layers 2/3 or 5 from one of six areas (Table S1, Table S2): the primary visual cortex (V1), secondary visual areas (AM, PM, MMA, MMP; as in (Zhuang et al. 2017)), and retrosplenial cortex (RSC). From referencing the retinotopic map to skull landmarks (Methods), areas AM and MMA in our functionally defined imaging sites may coincide substantially with the stereotactically defined coordinates of posterior parietal cortex (PPC) in related studies (Harvey, Coen, and Tank 2012; Morcos and Harvey 2016; Runyan et al. 2017; Goard et al. 2016; Hwang et al. 2017), and may also be functionally similar to PPC as noted in (Minderer, Brown, and Harvey 2019). After correction for rigid brain motion, regions of interest representing putative single neurons were extracted using a semi-customized (Methods) demixing and deconvolution procedure (Pnevmatikakis et al. 2016). The fluorescence-to-baseline ratio Δ*F*/*F* was used as an estimator of neural activity, and only cells with ≥0.1 transients per trial were selected for analysis. In total, we analyzed 10,481 cells from 145 imaging sessions.

### Neural populations in all recorded areas contain a variety of present and past task-related information from sensory to internal/cognitive

In evidence-accumulation tasks, the visual areas are often assumed to perform basic sensory processing that provides necessary input to—but are not otherwise involved in—accumulation and decision formation (Nienborg and Cumming 2009; Wimmer et al. 2015; Haefner, Berkes, and Fiser 2016). We investigated whether such a division of function is actually reflected in the neural activity of these areas, by asking if they contained information about various task-related quantities.

The bulk of neural activity in all areas and layers followed choice-specific sequences of activation previously reported in PPC (Harvey, Coen, and Tank 2012; Morcos and Harvey 2016). Individual neurons were active for short time intervals (Fig. 2C) that within a trial were staggered in time across the population, with ~ 75% of cells preferentially active in either right- or left-choice trials (Fig. 2A; Fig. S2A-C for θ-control experiments). This structure of activity was highly consistent across trials as well as similar for primary vs. secondary visual areas: ~ 90% of cells were active in specific place/time locations (Fig. S1A), and these cells were active in ~ 60% of preferred-choice trials. RSC differed in having a more uniform spacing in between peak activity locations across cells (Fig. 2B; *p* ≤ 10^−4^, K-S test), and cells being less reliably active in preferred-choice trials (~ 45%; Fig. S1B). In all cases, cells with ipsilateral-vs. contralateral-choice preferences were intermixed in the same brain hemisphere (Fig. S1E). Signals could also be identified in all areas/layers that seemed explicitly sensory, in that 5-10% (~2% in RSC) of cells responded in a time-locked manner to individual cues (“cue-locked” cells, discussed in the next sections). As most of these cells responded to a preferred laterality of cues, they also had choice specificity due to the correlation between choice and stimulus (red-highlighted rows in Fig. 2A). Apart from the above quantitative details that differed mostly between the visual areas and RSC, the overall form of neural activity was similar across areas and layers, and resembled previous findings for PPC.

**Figure 2.**
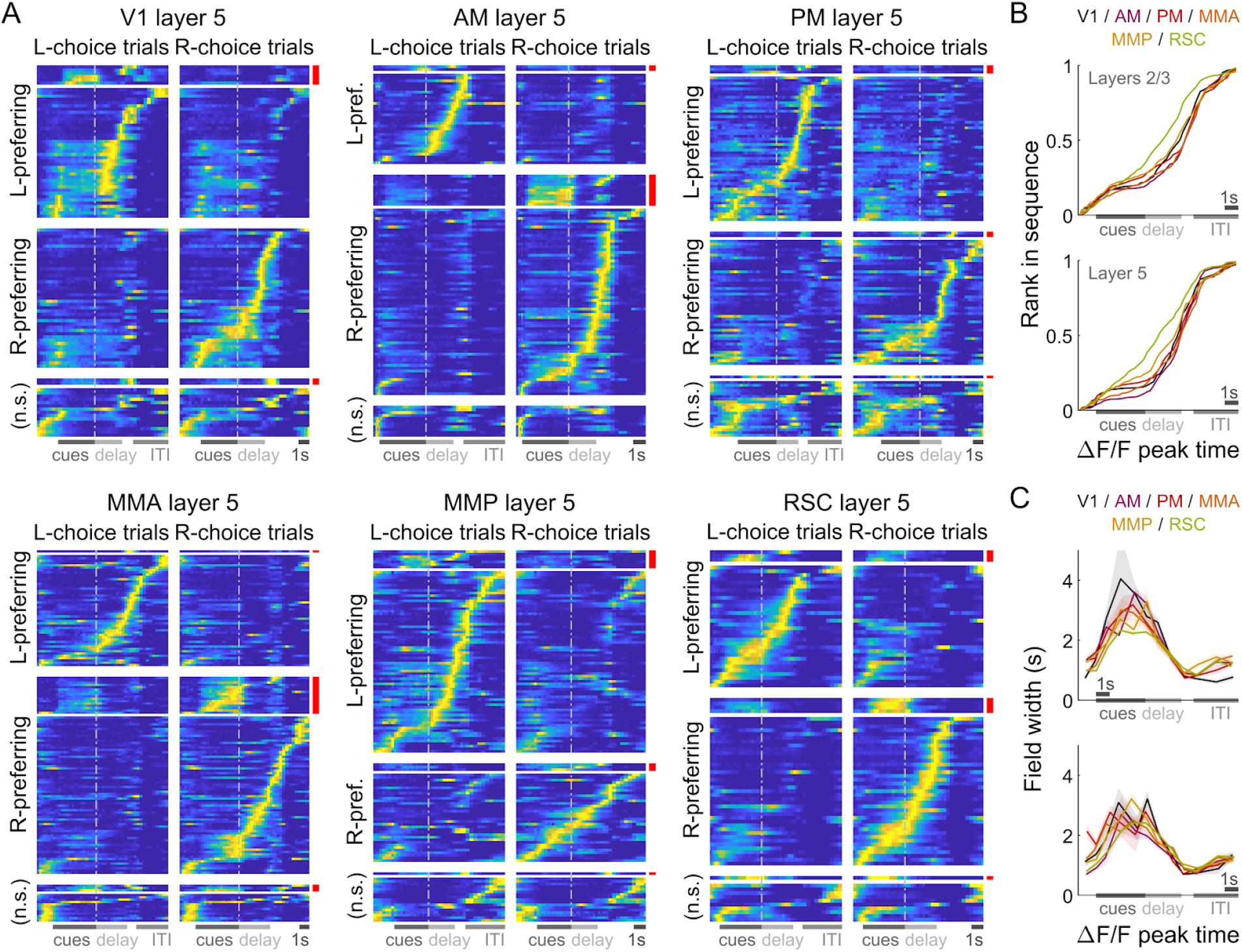
Neural activity in all areas/layers were qualitatively similar, and followed choice-specific sequences of activation throughout the trial. **(A)** Normalized and trial-averaged activity of cells (rows), for single example imaging sessions in the six recorded areas (*n* = 106,122,123,148,132,89 cells respectively). Cells were divided into left-/right-choice preferring populations, and sorted by the location of peak activity in correct preferred-choice trials. Cue-locked cells were separately sorted (red bars). Error trials were excluded in this analysis. **(B)** Rank of sorted cells vs. the location of peak activity, excluding cue-locked cells. Data were pooled across sessions for a given area (colors) and layer (top vs. bottom plots). **(C)** Duration of firing fields vs. location of peak activities. The firing field is defined as the span of time-points with activity at least half the height of the peak (above baseline) in trial-averaged data. Data were pooled across sessions for a given area/layer. Line: Mean across cells. Bands: S.E.M.

Although the majority of cells had striking temporal preferences and could be described as responding to cues or a particular place/time in the trial, the magnitudes of these responses differed depending on other behavioral contexts, only one of which is choice. Even if the subset of neurons that are active in different time periods may be very different, these context-dependent changes in activity could be coordinated across the neuronal population to encode information stably through time. To quantify such population-level information, we trained a separate support vector machine (SVM) per time-point in the trial to linearly decode a given behavioral quantity *X* from the Δ*F*/*F* vector of simultaneously recorded neurons (neural population state; see Methods). As discussed below, *X* is either the evidence defined as accumulated counts of contralateral (ipsilateral) cues, choice, or reward outcome, and the following show that all areas/layers contain information beyond that which is purely sensory.

For evidence decoding, we excluded explicitly sensory a.k.a. cue-locked cells from the neural population state, and dissociated count information from choice by only using trials of the same choice laterality (e.g. contralateral-choice trials for decoding contralateral counts). Fig. 3A-B shows the correlation between the actual counts and the value predicted from neural activity in various areas/layers, with timepoints colored when this is significantly better than chance (cross-validated and corrected for multiple comparisons, see Methods; Fig. S2D for θ-control experiments). In all areas and layers, this correlation was high throughout the cue period and gradually decayed in the delay period, but intriguingly rose again in the inter-trial-interval (ITI). In fact, during the cue period information about evidence in the *previous* trial remained strongly present, at levels comparable to evidence in the present trial (right vs. left plots of Fig. 3A-B).

**Figure 3.**
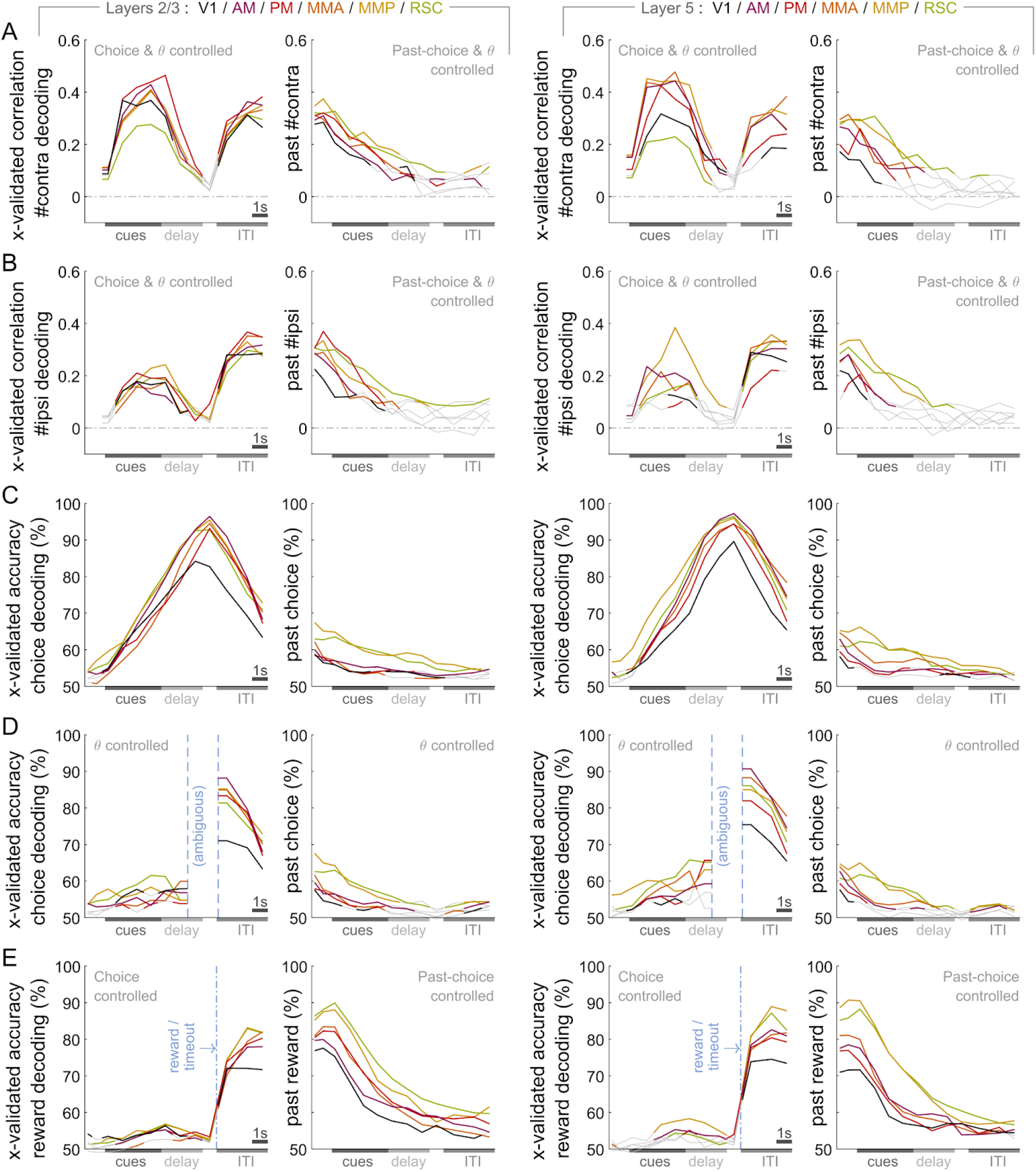
Neural populations in all areas/layers contained information about evidence, choice and trial reward outcome, including from the past trial. **(A)** Correlation between actual and predicted number of contralateral cues from neural population states vs. time, in various areas (lines) and layers (columns), averaged across cross-validation test sets and across mice. Time-points are colored if the decoding correlation is significantly different from chance (permutation test; *p* < 1.1 × 10 ^−4^ corrected for multiple comparisons). **(B)** Same as **(A)** but for ipsilateral cues (*p* < 7.1 × 10^−5^) **(C)** Same as **(A)**, but for the accuracy of classifying choice (*p* < 1.5 × 10^−4^). **(D)** Same as **(C)**, but controlling for view angle by equalizing θ distributions across choice (Methods; *p* < 8.1 × 10^−5^). Ambiguous time-bins are where θ is near-perfectly correlated with choice. **(E)** As in **(C)**, but for decoding whether the mouse received a reward. (*p* < 1.7 × 10^−4^).

Analogously, we decoded choice independently of cue/evidence information by excluding cells that were cue-locked or had activity that depended on Δ = #*R* − #*L* for a fixed choice (Methods). Fig. 3C shows that in all areas, choice classification accuracy rose throughout the cue period. This could reflect the behavioral finding that mice accumulated evidence over an extended period (Fig. 1F), i.e. during which the eventual choice should not be fully predictable. Visual scene differences in right-vs. left-choice trials did not account for all of the choice decoding accuracy (see Discussion). This is explicitly so in control experiments where we eliminated scene differences by enforcing the view angle θ to be zero up to midway in the delay period (Fig. S2E,I-J), as well as in analyses where we weighted the data to have the same θ distribution for both choice categories (Fig. 3D, see Methods). Choice information also persisted throughout the ITI when there was no visual input.

Lastly, we used the same method to decode reward outcomes while controlling for choice. Outcome decoding accuracies were high in all areas/layers starting from the end of the trial, and persisting well into the next trial (Fig. 3E; Fig. S2F for θ-control experiments). In sum, information about evidence, choice, and reward outcome were significantly present in all areas and layers, albeit more weakly in V1. This includes past-trial records of all these variables particularly during the ITI and cue period, with the strongest signals in the most medial areas, RSC and MMR

### Pulses of evidence evoke transient, time-locked responses in all recorded areas

In searching for sensory responses, we indeed found neurons in all areas/layers that had activities clearly time-locked to the pulsatile cues (examples in Fig. 4A-B). In trials with sparse occurrences of preferred-side cues, the activities of these cells tended to return to baseline following a fairly stereotyped impulse response. Individually they thus seemed to code only information about momentary cues, although as a population they can form a more persistent stimulus memory (Goldman 2009; Scott et al. 2017; Miri et al. 2011). Interestingly, the amplitudes of these cells’ responses seemed to be variable in a structured way, both across time in a trial, as well as across trials where the mouse made right vs. left choices (columns in Fig. 4A-B). We wanted to know if these fluctuations also encoded task-relevant information.

**Figure 4.**
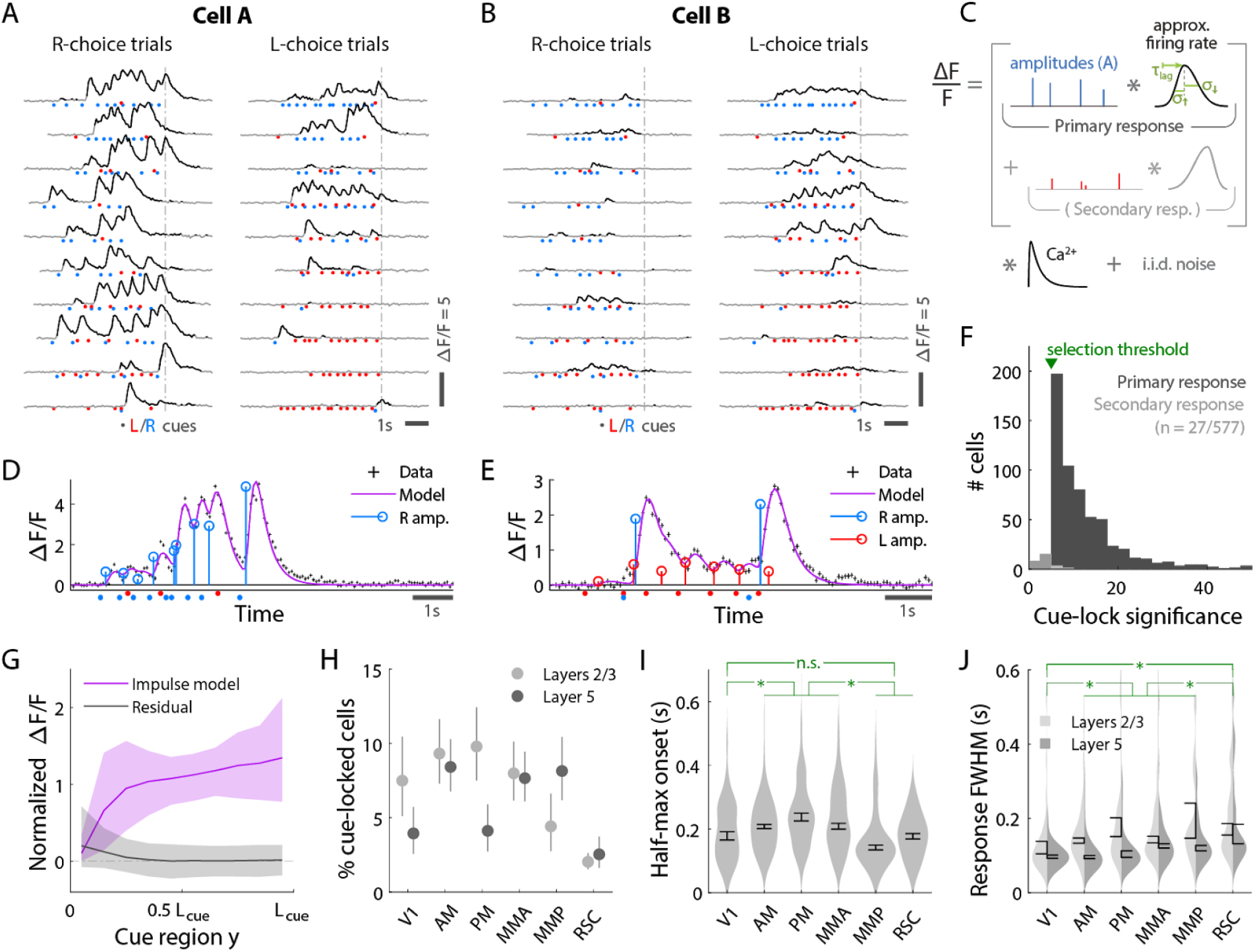
Pulses of evidence evoke transient, time-locked responses that are well described by an impulse response model. **(A)** Trial-by-trial activity (rows) vs. time of an example right-cue-locked cell, aligned in time to the end of the cue period (dashed line). Onset times of left (right) cues in each trial are shown as red (blue) dots. **(B)** Same as **(A)**, but for an *atypical* right-cue-locked cell that has some left-cue-locked responses. **(C)** Depiction of the impulse response model. **(D)** Prediction of the impulse response model for the cell in **(A)** in one example trial. This cell had no significant secondary (left-cue) responses. **(E)** Same as **(D)** but for the cell in **(B)**. The model prediction is the sum of primary (right-cue) and secondary (left-cue) responses. **(F)** Distribution of cue-locking significance for cells with a significant primary response (above 5 standard deviations compared to cues-shuffled fits). **(G)** Trial-averaged impulse response model prediction vs. location in the cue region (purple), compared to the residual (data minus model prediction, black). Δ*F*/*F* was normalized to the mean prediction per cell. The model prediction rises gradually from baseline at the beginning of the cue period due to nonzero lags in response onsets. Line: Mean across cells. Band: 95% C.I. **(H)** Percent of significantly cue-locked cells in various areas/layers. Chance: 3 × 10^−5^%. Error bars: 95% C.I. **(I)** Distribution (kernel density estimate) of the half-maximum onset time of the primary response, for cells in various areas. Data were pooled across layers (inter-layer differences not significant). Error bars: S.E.M. Stars: significant differences in means (Wilcoxon rank-sum test). **(J)** As in **(I)** but for the full-width-at-half-max. Statistical tests use data pooled across layers. Means are significantly different across layers for all areas except V1 and RSC (Wilcoxon rank-sum test).

For a given cell, we estimated the amplitude of its response to each cue *i* by modeling the cell’s activity as a time series of non-negative amplitudes *A_i_* convolved with an impulse response function (Fig. 4C). The latter was defined by lag rise-time and fall-time parameters that were fit to each cell, but were the same for all cues for that cell (deconvolving calcium dynamics; see Methods). For a subset of neurons, this impulse response model resulted in excellent fits when the model included only primary responses to either right- or left-side cues (e.g. Fig. 4D). In much rarer instances, adding a secondary response to the opposite-side cues resulted in a significantly better fit (e.g. Fig. 4F). We defined cells to be cue-locked if the primary-response model yielded a much better fit to the data than a permutation test (discounting for the number of parameters by using AIC_c_ as goodness-of-fit, see Methods and (Hurvich and Tsai 1989)). For these cells, the trial-averaged activity predicted by the impulse response model (Fig. 4G, magenta) was significantly and substantially larger than the residuals of the fits, which had negligible magnitudes and time-structure (Fig. 4G, black). This indicates that there were no large unaccounted-for components, particularly those with long timescales such as a systematic rise in baseline activity.

Significantly cue-locked cells comprised a small fraction of the overall neural activity, but were nevertheless present in all areas/layers and exhibited some progression of response timescales from V1 to other areas. About 5-10% of cells in visual areas were significantly cue-locked, compared to ~ 2% in RSC (Fig. 4H; Fig. S4B for θ-control experiments). Of these, only ~ 5% had secondary responses that were moreover much less significantly time-locked (Fig. 4F); most cells responded to only contralateral cues (Fig. S3A). The onset of the half-maximum response was ~ 200*ms* after each pulse (Fig. 4I), and the response full-width-at-half-max (FWHM) was ~ 100*ms* but increased from V1 to secondary visual areas to RSC (Fig. 4J; θ-control experiments Fig. S4C). The impulse response model thus identified cells that, on a cue-by-cue basis, follow what one might expect of purely visual-sensory responses, up to amplitude changes that we next discuss.

### Cue-locked responses are amplitude-modulated by present and past cognitive quantities in all recorded areas

Studies of perceptual decision-making have shown that the animal’s upcoming choice affects the activity of stimulus-selective neurons in a variety of areas (Britten et al. 1996; Nienborg and Cumming 2009). We analogously looked for such effects (and more) while accounting for the highly dynamical nature of stimuli in our task, focusing on the primary-response amplitudes {*A_i_*} of cue-locked cells. Importantly, the impulse response model deconvolves responses to individual cues, so *A_i_* may be conceptualized as a multiplicative gain factor at the instant of the *i^th^* cue. In observations like Fig. 4A-B, these amplitudes appear to systematically depend on time as well as choice. This may indicate coding of place/time and choice as we have found for non-cue-locked cells, but may also arise indirectly from correlations with other behavioral variables. Most obviously, there may be a receptive field given by the visual angle of the cue (**φ**_*cue*_, Fig. 1C), as well as modulations due to running speed (Niell and Stryker 2010; Saleem et al. 2013) and navigational location (Saleem et al. 2018). A graded dependence on cue counts is another candidate explanation, as cue counts rise during the trial. An effective count dependence may also be due to stimulus-specific adaptation (SSA; (Ulanovsky, Las, and Nelken 2003; Sobotka and Ringo 1994)) or enhancement (Vinken, Vogels, and Op de Beeck 2017; Kaneko, Fu, and Stryker 2017).

Given limited data statistics, we only compared a small set of six conceptually distinct hypotheses involving the above factors. All six models assume the amplitudes to be drawn from a Gamma distribution, with mean given by an angular receptive field shape that is scaled by other effects as follows (summarized in Table S3; see Methods). The baseline hypothesis is the angular-receptive-field-only model. Next, the “*v*” model additionally depends on running speed. The remaining four models all include **φ**_*cue*_ and speed parameters, but multiplied by dependencies on different factors that can explain time-dependent trends. The “SSA” model parameterizes adaptation/enhancement with exponential time-recovery in between cues. The “*y*” model depends on the spatial location of the cue. The “*C*” model has location and speed dependencies indexed by choice. Lastly, the model has cue-count dependence (#*R*, #*L*, or Δ = #*R* − #*L*). This selection of models allows us to ask if cue-locked responses are sufficiently explained by previously known effects, or if after accounting for such there are still effects related to the accumulation process, such as choice or cue-count dependence.

We constructed the amplitude model prediction as the AIC_c_-likelihood-weighted average of the above models, which accounts for when two or more are comparably good (Volinsky et al. 1999). As illustrative examples, Fig. 5A shows how the amplitudes of two simultaneously recorded cue-locked cells in area AM depended on various factors and compared to model predictions. There are clear differences in predictions for right-vs. left-choice trials that can also be seen in the raw amplitude data (2^nd^-4^th^ columns of Fig. 5A; Fig. S4A for θ-controlled experiments). Although both cells responded preferentially to right-side cues, they had oppositely signed choice modulation effects, defined as the difference between amplitude model predictions on contralateral-vs. ipsilateral-choice trials (Methods). Fig. 5B shows another example cell that had no significant dependence on choice, but instead on the accumulated evidence Δ. These three cells are typical of how angular receptive field and running speed effects were often insufficient to explain choice- and count-dependent trends. The dominantly choice-dependent and not count-dependent predictions for cells A and B (Fig. 5A) and conversely for cell C (Fig. 5B) are also exemplary of how large fractions of cells strongly preferred one model over the other (relative likelihoods in Fig. S3B-C).

**Figure 5.**
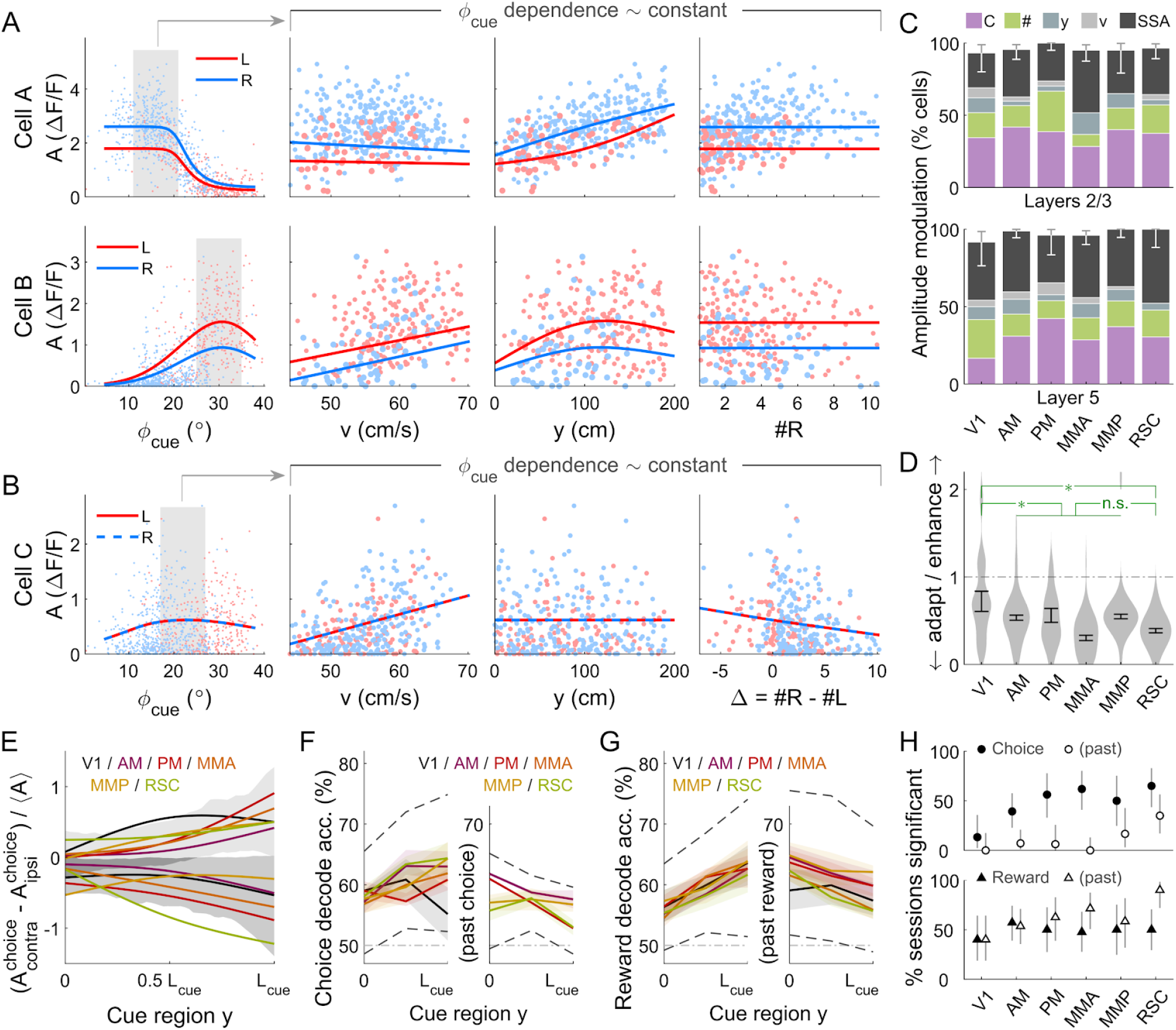
Cue-locked cell amplitudes are modulated by visual angle, speed, location, cue frequency, counts, and choice. **(A)** Response amplitudes of two example right-cue-locked cells (rows) vs. various behavioral variables (columns). Data are marked in blue (red) according to the upcoming right (left) choice, and model predictions are shown separately for right-vs. left-choice trials (lines). Cue counts are slightly jittered for visualization. The data in the rightmost three columns were restricted to a subset where angular receptive field effects are small, corresponding to the shaded area in the leftmost plots. **(B)** Same as **(A)** but for a significantly Δ-modulated neuron. **(C)** Percentages of cells that favor various amplitude modulation models. Error bars: 95% C.I. for sum over the indicated models; the remaining fraction are cells that favor the angular-receptive-field-only model. **(D)** Distribution (kernel density estimate) of adaptation/enhancement factors for cells that favor the SSA model. A factor of 1 corresponds to no adaptation, while for other values the subsequent response is scaled by this amount with exponential recovery towards 1. Error bars: S.E.M. Stars: significant differences in means (Wilcoxon rank-sum test). **(E)** Choice modulation strength vs. place/time, for contralateral-cue-locked cells only (ipsilateral-cue-locked cells in Fig. S3I). Lines: mean across cells in a given area. Bands: std. dev. of data pooled across all areas/layers. Both mean and std. dev. are shown separately for cells with positive vs. negative modulations. **(F)** Cross-validated accuracy of decoding the upcoming (left half) and past-trial (right half) choice from cue-response amplitudes. Solid lines: means across datasets for which this was significant, by area. Bands: S.E.M. by area. Dashed lines: 95% interval for data pooled across areas. **(G)** Like **(F)** but for decoding reward outcome. **(H)** Percents of imaging sessions that had significant choice (top plot) and reward (bottom plot) decoding accuracy. Error bars: 95% C.I. Differences in means (Wilcoxon rank-sum test) for upcoming-choice decoding are significant only for V1 vs. secondary visual areas and vs. RSC, and for past-choice decoding only for V1 vs. RSC. Differences for upcoming-reward decoding are not significant, and for past-reward are significant only for V1 vs. RSC.

To summarize the prevalence and composition of effects, we selected the best model per cell using AIC_c_, defaulting in ambiguous cases to already-known effects (angular-receptive-field-only or speed model, in that order). In all areas and layers, > 90% of cue-locked cells exhibited some form of amplitude modulations beyond angular receptive field effects (Fig. 5C; Fig. S4E for θ-controlled experiments). Overall, 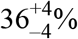 of cells were best explained by SSA while 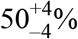 favored either choice or cue-counts models, with non-significant differences across layers (*p* = 0.11 and *p* = 0.09 respectively, z-test). There were also no significant inter-area differences in proportions of cells that preferred SSA, choice, or cue-counts models, except marginally for the choice model in layer 5 of V1 vs. PM (*p* = 0.048, z-test). While there is likely a continuum of cells including those with mixed effects, most cells fall into approximate categories with SSA, choice- or count-related modulations, with remarkably little difference in composition across areas and layers.

Cells in the two largest categories, SSA and choice, had qualitatively different population statistics for how their cue-response amplitudes depended on place/time in the trial. Most cells 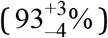 that favored the SSA model corresponded to a phenotype with decreased responses to subsequent cues. Adaptation effects were weakest in V1 and stronger in secondary visual areas and RSC (Fig. 5D, but see Fig. S4F for θ-controlled experiments), although the ~ 0.8s recovery timescale had no significant inter-area differences except for PM vs. RSC (Fig. S3E, *p* = 0.027, Wilcoxon rank-sum test). In contrast, cue-locked cells with both choice laterality preferences were intermixed in all areas and layers (Fig. S3F-G; θ-controlled experiments Fig. S4D). Both subpopulations of positively and negatively modulated cells exhibit gradually increasing effects vs. place/time (Fig. 5E, Fig. S3I).

As the above are reminiscent of the array of effects we have described for non-cue-locked cells, we wondered if long-timescale choice and reward information could similarly be decoded from cue-response amplitudes. We defined the neural population state as the vector of amplitudes in response to contralateral cues only, and otherwise applied the same methods as for Fig. 3. Fig. 5F shows that choice could be decoded from cue-locked amplitudes as expected from Fig. 5E, but that the choice in the previous trial could also be decoded, mostly in the medial areas (Fig. 5H-top). Reward outcomes in both present and past trials could reliably be decoded in all areas (Fig. 5G,H-bottom). During the cue region, there is an apparent ability to predict the upcoming reward (Fig. 5G-left), which likely corresponds to count-related amplitude modulations (“#” model in Fig. 5C). Outcome decoding accuracies could be as high as that for non-cue-locked cells (Fig. 3C-E), in datasets with just 5-15 cue-locked cells (70-80% accuracies, Fig. S3J). Therefore, far from being purely sensory responses, the amplitudes of cue-locked cells instead reflected present- and past-trial information just like the rest of the neuronal population.

### Cue-locked amplitude modulations motivate a multiplicative feedback-loop circuit model

We hypothesize that the cue-locked sensory responses in visual areas provide momentary cue information that drives an accumulation process, which ultimately drives choice. If this is true, modulations of cue-response amplitudes by variables such as evidence and choice (Fig. 5C) may predict specific perceptual biases in decision making. We note that relationships between sensory responses and choice can arise in a purely feedforward circuit structure (Shadlen et al. 1996), because the sensory neural responses have a causal role in producing the behavioral choice. However, as previously argued this should result in similar timecourses of neural and behavioral fluctuations (Nienborg and Cumming 2009); instead, we observed contrasting timecourses: as each trial evolved, there was a slow *increase* in time in choice modulations of cue-locked responses (Fig. 5E-F; Fig. S3I), which was opposite to the behaviorally-assessed *decrease* in time in how sensory evidence fluctuations influenced the mice’s choice (Fig. 1F). Additionally, a feedforward structure predicts that positive fluctuations in right-(left)-preferring cue-locked neurons should produce rightwards (leftwards) fluctuations in choice. Instead, we observed that about half of the cue-locked cells were modulated by choice in a manner opposite to their cue-side preference (Fig. S3F-G). Both of these observations argue against a purely feedforward structure and thus support the existence of feedback influences on sensory responses (Wimmer et al. 2015; Nienborg and Cumming 2009; Haefner, Berkes, and Fiser 2016). We therefore propose that feedback projections from a downstream pulse accumulator induce choice- and pulse-count-modulations of cue-locked sensory responses. We used a simplified dynamical systems model to ask what behavioral consequences would be predicted by such feedback. The model we considered has separate accumulators for the right and left stimulus streams, as psychophysical analyses suggest that there are two near-independent accumulators which are then compared to form the behavioral decision (Scott et al. 2015; Brunton, Botvinick, and Brody 2013). We thus describe below a model for a right-side-specific accumulator given by a single scalar variable *a_r_*(*t*); identical results hold for a left-accumulator *a_l_*(*t*).

Based on neural and behavioral findings, we made a few simplifying assumptions that allowed the model dynamics to be solved for analytically (details in Methods). The sensory units are described by an *n_r_*-dimensional activity vector 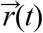, all of which receive the same time-varying input stimulus, *R*(*t*). Because psychophysical performance depended on total accumulated pulse counts but not on the duration of the pulse stream (Fig. 1G; (Scott et al. 2015; Brunton, Botvinick, and Brody 2013; Lucas Pinto et al. 2018)), we assumed that the accumulator *a_r_* performs leak-free integration of its input. At each point in time, we took the input to the accumulator *a_r_* to be a weighted sum of the activities of the *n_r_* sensory units. Thus 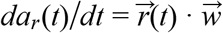, where 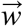 is the vector of input (feedforward) weights. A central feature of our model is that *a_r_*(*t*) feeds back as a dynamic, *multiplicative* gain control on the input to sensory units. By this we mean that the sensory responses 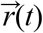 follow the input drive *R*(*t*) plus a multiplicative factor:

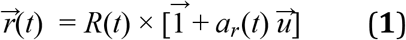

Here 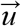 specifies the accumulator feedback weights for each sensory unit, and 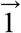 is the *n_r_*-dimensional vector with 1 for each coordinate. Note that, inspired by the transience of cue-locked neural responses, we have assumed in Eq. 1 that the sensory unit activities 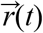 instantaneously follow their input.

As illustrated in Fig. 6A, this feedback-loop model can produce increasing/decreasing sensory responses depending on whether the feedback weight *u_i_* for the *i^th^* sensory unit is positive or negative. Because the accumulator-modulated sensory activity is again fed forward into the accumulator, the circuit compounds the effect of the stimulus and can produce exponentially increasing accumulator values. In sum, the accumulator dynamics are given by

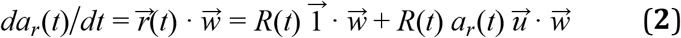

The solution to this is (Methods):

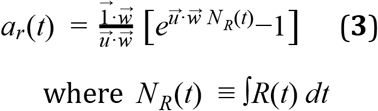

**Figure 6.**
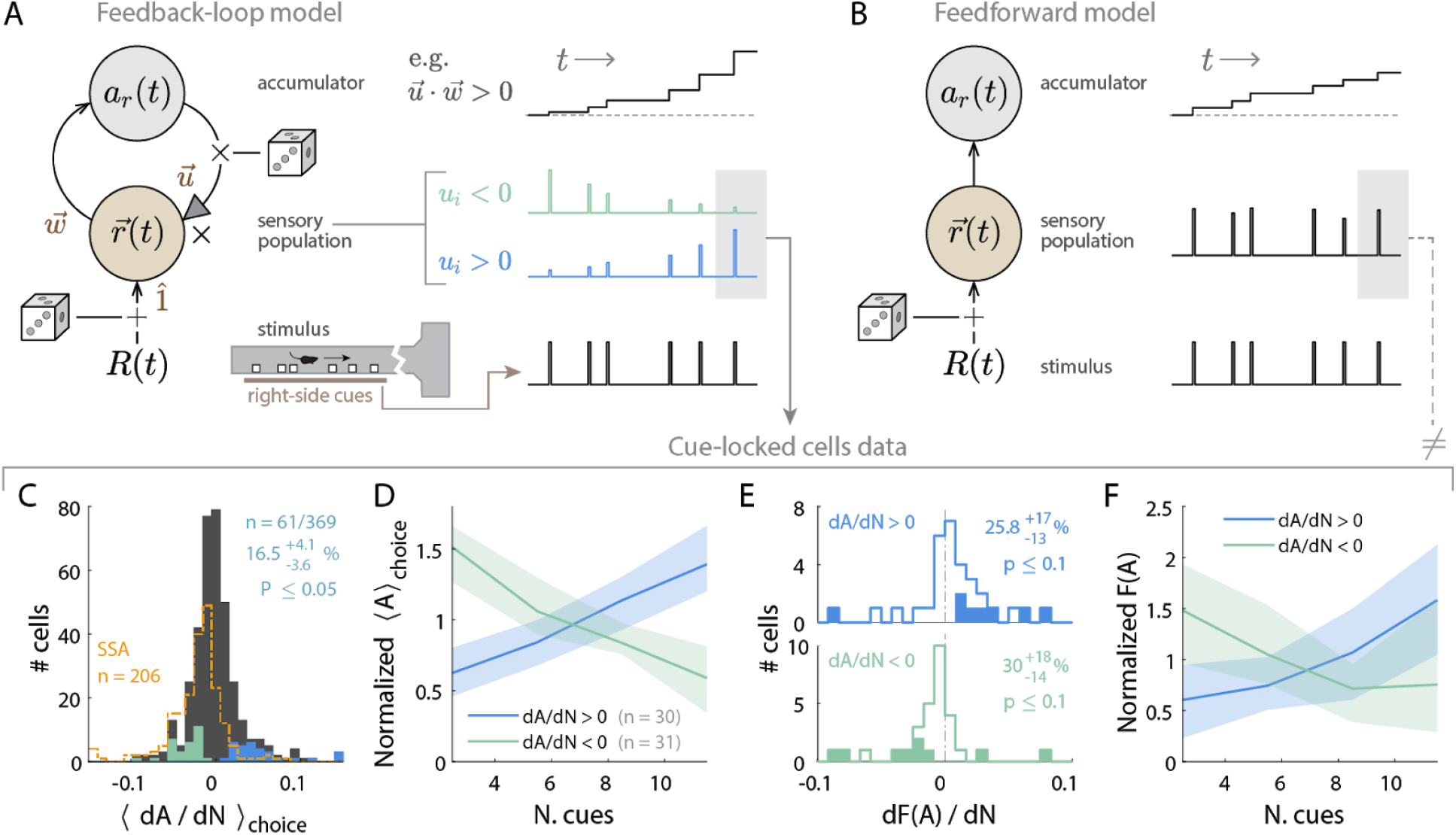
A multiplicative feedback-loop circuit explains cue-locked cell amplitude modulations vs. counts. **(A)** Left: Schematic of the feedback-loop circuit model of a right-stimulus-stream accumulator. Dice indicate locations where additive/multiplicative noise may arise. Right: Single-trial illustration of time traces for dynamics at the sensory vs. accumulator stages of the model, for a given right-stimulus stream (bottom). **(B)** As in **(A)**, but for a purely feedforward model. **(C)** Distribution of choice-averaged slope (*dA*/*dN*) for the linear regression model of amplitude *A* vs. number of preferred-side cues *N*. Cells with significant slopes vs. a permutation test are highlighted in color (excluding cells compatible with SSA, dashed histogram). **(D)** Choice-averaged amplitude at the end of the cue region vs. the total number of preferred-side cues. To account for cell-specific activity rates, this is normalized to the average across counts per cell. Only cells with significant cue-count slopes as highlighted in **(C)** were included. Line: averages across cells with positive vs. negative slopes. Band: std. dev. across cells. Data were pooled across areas and layers. **(E)** Distribution of slopes from linear regression of the Fano factor vs. preferred-side cue counts, separately for positively and negatively modulated cells as in **(C)**. Cells with significant Fano factor slopes vs. a permutation test are highlighted in color. **(F)** Fano factor at the end of the cue region vs. total number of preferred-side cues, normalized to the average across counts per cell. Only cells with significant cue-count as well as Fano-factor slopes as highlighted in **(E)** were included. According to the model, non-monotonic trends are possible for negatively count-modulated cells only (see Methods for derivation). Line: averages across cells with positive vs. negative *dA*/*dN* slopes. Band: std. dev. across cells.

This is in contrast to a purely feedforward version 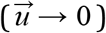, which would result in pure integration:

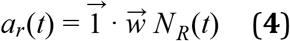

Note that the lack of a leak term in Eqs. 3 and 4, and the multiplicative nature of the feedback in Eq. 3, means that when the input drive *R*(*t*) = 0 then *da_r_*(*t*)/*dt* = 0, i.e. the accumulators are stable. In fact, a feature of Eqs. 3 and 4 is that the accumulator value does not depend explicitly on time, only on the cumulative stimulus input up to that time, *N_R_*(*t*) for arbitrary *R*(*t*). The intuition for pulsatile stimuli is that all changes to the sensory and accumulator states are gated by the stimulus drive, so the entire system only changes state when there is a pulse and is not sensitive to the time interval between pulses. Consequently, psychophysical predictions of these two models, and comparisons between them, depend only on the net stimulus *N_R_* at the end of a trial, and not otherwise on trial duration. In contrast to multiplicative feedback, it is also possible to achieve time-independent integration in an architecture with additive feedback, so long as there is also an appropriately tuned amount of accumulator leak (Seung 1996). However, in such circuits the accumulator continues to drive sensory units (defined as those that directly receive stimulus input) even in the absence of stimuli (Wimmer et al. 2015; Seung 1996). For pulsatile stimuli, this predicts changes in baseline sensory-unit activities in between pulses that do not match our observations of cue-locked cells (Fig. 4A-B,G). A multiplicative architecture, where sensory responses are scaled (Eq. 1) by an accumulator that depends on only the cumulative stimulus (Eq. 3), is thus consistent with both neural and behavioral features that we set out to model.

Our feedback-loop model makes predictions about two properties of sensory unit activities that are compatible with observed trends in the amplitudes of cue-locked cells. First, sensory unit activities in response to pulsatile stimuli have heights linearly related to the accumulator contents (Eq. 1). To focus specifically on possible dependencies on accumulated counts, we first excluded cue-locked cells with responses consistent with stimulus-specific adaptation (SSA; Fig. 5C), and restricted the data to the last-cue response in a trial (restricted to the last third of the cue period, to minimize differences correlated with place/time). We asked whether the last-cue response amplitude *A* also depended on cue count *N*, independently of choice and view angle effects (as shown in Fig. 5). We thus linearly regressed the last-cue response amplitude *A* vs. cue counts *N*, controlling for choice by performing this separately in right-vs. left-choice trials, then averaging the *dA*/*dN* slope across choice (Methods). Within a given choice category, we also controlled for view angle θ by weighting trials so that the θ distribution is the same across cue-counts (Methods). We found two comparably sized subpopulations with significantly positive or negative *dA*/*dN* slopes (Fig. 6C, cyan and green entries; Fig. S4G for θ-controlled experiments). In the model, these two subpopulations would correspond to positively-modulated (*u_i_* >0) and negatively-modulated (*u_i_* < 0) sensory units respectively. For cells with significantly nonzero slopes vs. a permutation test (Methods), Fig. 6D shows that the count modulation effect size is large and fairly consistent across cells, resulting in a factor of ~ 3 change in amplitudes over the behaviorally encountered range of cue counts *N*. The second prediction of the feedback-loop model is that the Fano factor, *F*(*A*) ≡ *var*(*A*)/*mean*(*A*), should increase (decrease) with *N* for positively-(negatively-)modulated sensory units (Methods; illustrated in Fig. S5A-B). Although other mechanisms can also produce non-constant Fano factors, a linear regression model of *F*(*A*) vs. *N* (Methods) indeed showed a compatible prevalence of positive (negative) Fano-factor slopes for the two subpopulations of cells with significantly positive (negative) *dA*/*dN* slopes (Fig. 6E, top vs. bottom, effect size in Fig. 6F; Fig. S4H for θ-controlled experiments). In these ways, the feedback-loop model predicts cue-count dependencies for sensory unit activities that qualitatively match observations for cue-locked cells. This is in contrast to purely feedforward architectures (Fig. 6B; see Eq. 4), where sensory responses 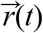 have no systematic count dependence (Methods).

### A multiplicative feedback-loop architecture best explains asymptotic Weber-Fechner scaling in psychophysical performance

Because a multiplicative feedback-loop circuit hypothesis can explain count-modulation of cue responses in the neural data, we wondered if it could also explain psychophysical features of pulsatile accumulation tasks. For otherwise identical models, the presence vs. absence of accumulator feedback on sensory units predicts two different mathematical forms (Eq. 3 vs. Eq. 4) for how left and right accumulator values depend on the input stimulus, and consequently potential differences in the accuracy of comparing the two accumulators to produce a choice. We compared which of the feedforward versus feedback forms best fits two sets of behavioral data: the full Accumulating-Towers dataset (Lucas Pinto et al. 2018), as well as data from rats performing a pulse-based visual accumulation task with no navigational requirements (Scott et al. 2015). For all these data, where the behavior has little dependence on time or trial history (Lucas Pinto et al. 2018; Scott et al. 2015), the fraction of correct trials is only a function of the numbers of cues *N_maj_* on the majority and *N_min_* on the minority side. Two long-standing theories predict qualitatively different trends for this function, yet neither fully matches the data. As described below, our feedback model yields a better fit than either of these famous theories.

First, evidence accumulation is often modeled as an integration process, for which the central limit theorem of statistics states that performance should improve vs. total counts (Fig. S5C; (Scott et al. 2015)). Second, the psychophysical Weber-Fechner Law contrastingly postulates that discriminability depends only on the ratio *N_min_*/*N_maj_*, i.e. performance should be constant vs. total pulse counts *N_tot_* = *N_maj_* + *N_min_* (Fig. S5D; (Fechner 1966)). In both mouse and rat data (Fig. 7A), we intriguingly observed both types of trends. For *N_min_*/*N_maj_* = 0 performance increased with *N_tot_* as qualitatively predicted—but turns out *not* to be fully explained by—the central limit theorem. In contrast, for *N_min_*/*N_maj_* = 0.4 there was little dependence of performance on *N_tot_*, consistent with Weber-Fechner Law.

**Figure 7.**
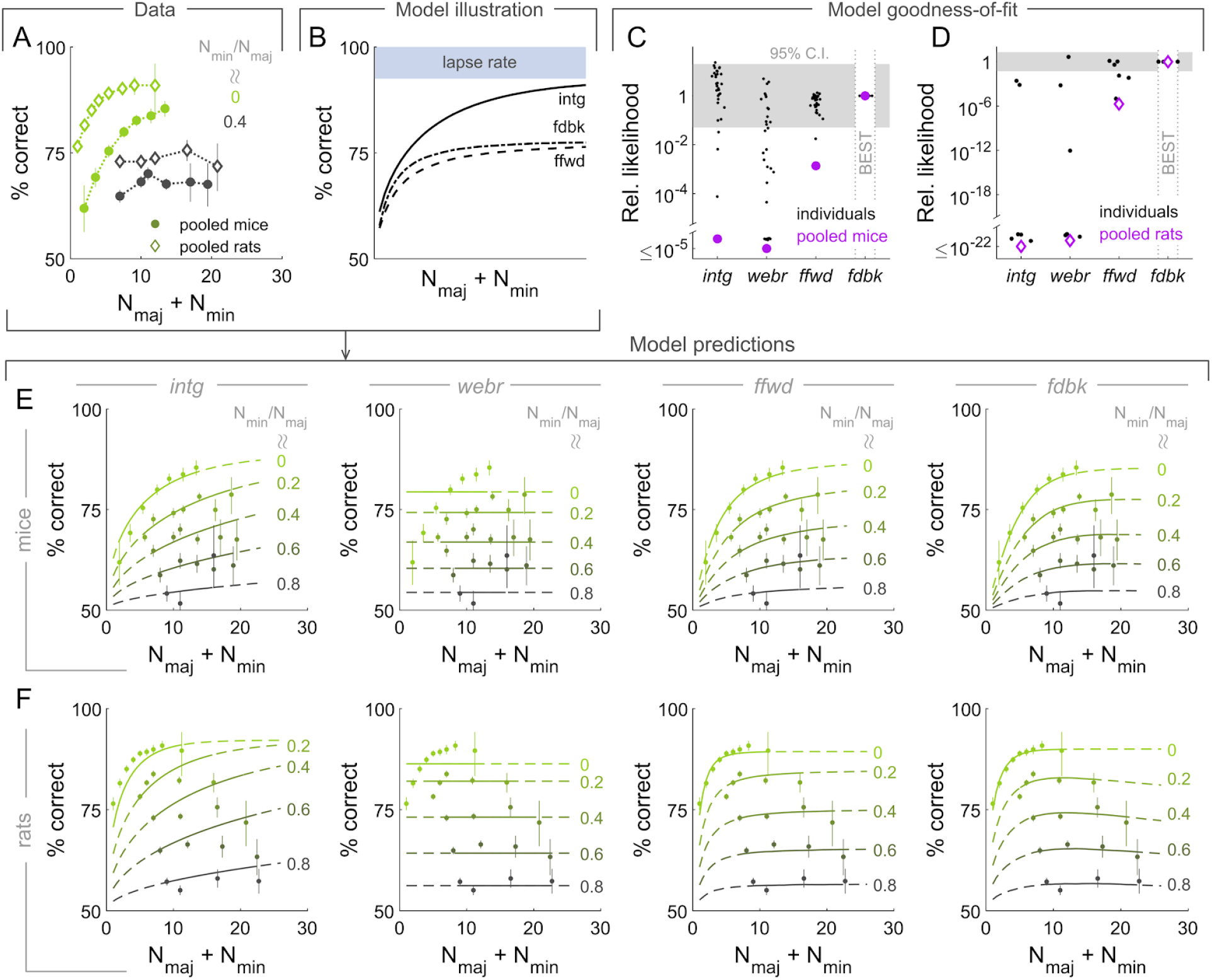
A multiplicative feedback-loop circuit best explains asymptotic Weber-Fechner scaling in perceptual performance, compared to models without feedback. **(A)** Behavioral performance (percent of correct trials) vs. total cue counts, for two fixed ratios of minority over majority cue counts. Data were pooled across mice (rats). Points are joined by lines to guide the eye. Error bars: 95% C.I. **(B)** Illustration of perceptual performance for circuit architectures described in the text, with *N_min_*/*N_maj_* = 0.5 and the same parameters for all noise distributions (ρ_1_ = 1, μ_*u*_ = 0, σ_*u*_ = 0.5, σ_*c*_ = 1, *p_lapse_* = 0.15; where relevant). The *webr* model predicts constant performance. **(C)** AIC_c_ likelihood ratios for models relative to the *fdbk* model, which best fits the pooled mouse data. **(D)** As in **(C)**, but for rat behavioral data. **(E)** Pooled mouse data as in **(D)**, compared to model predictions (lines) for the four models in **(A)**. **(F)** As in **(E)**, but for rat data vs. model predictions.

The above theories each assume that one dominant source of stochasticity drives the psychophysical performance of the accumulator circuit. For the integrator (“*intg*”) model subject to the central limit theorem, this source is a per-pulse sensory noise that is independent across pulses. For pure Weber-Fechner scaling to apply (“*webr*” model), independent per-pulse sensory noise should be negligible compared to a memory-level noise (Gallistel and Gelman 2000). To account for the data deviating from both these theories, we hypothesized that the accumulator circuit instead mixes effects from multiple significant sources of stochasticity. In total, we considered the effects of per-pulse sensory noise, slow modulatory noise that multiplies the sensory response, noise associated with comparing accumulators, and an evidence-independent lapse rate *p_lapse_*. The critical comparison here was then between two models that each had all these sources of stochasticity, but differed in whether there was accumulator-feedback onto the sensory units (“*fdbk*” model, Eq. 3), or if the circuit was purely feedforward (“*ffwd*” model, Eq. 4).

In the following, we summarize the setup of the *fdbk* and *ffwd* models, with details in the Methods. For both models, the effect of sensory noise is to replace the true integrated stimulus in Eqs. 3 and 4 with a Gaussian distribution. Together with a model-specific modulatory noise on sensory unit responses (described below), this leads to a distribution of possible values for the accumulator state in response to a given stimulus. We model noise in the operation of comparing right-side (*a_r_*) to left-side (*a_l_*) accumulators as the decision variable (*c*) being additionally Gaussian-distributed around *a_r_*−*a_l_*. This decision variable predicts that the subject will make a choice to the right with probability *P*(*c* > 0), up to a fraction of lapse trials where the subject instead makes a random choice. The difference in the modulatory noise source between models, is that for the *fdbk* model we hypothesized that the accumulator feedback is noisy or equivalently that the feedback weight is stochastic per trial. This is in contrast to the *ffwd* model, where we hypothesized a non-specific source of slow gain fluctuations, equivalent to scaling the accumulator value with a stochastic variable per trial.

Fig. 7B illustrates how the predicted perceptual performance of the above models depend on total cue counts *N_tot_* = *N_min_* + *N_maj_*, for a fixed ratio *N_min_*/*N_maj_* = 0.5 and identical noise parameters. Both *ffwd* and *fdbk* models predict a transition between a regime at low counts where sensory and accumulator-comparison noise have strong effects, seen in the rising trend of the performance curve, and a Weber-Fechner regime at high counts where performance saturates at less than the lapse rate because modulatory noise dominates (formally shown in the Methods). The distinction between these models is in how the *fdbk* model reduces to simple integration with no modulatory noise at low counts, and thus predicts a steeper rise in performance vs. counts than the *ffwd* model, where modulatory noise is always at play. At high counts, both model predictions converge and exhibit Weber-Fechner scaling.

These four accumulator models have free parameters that specify the distributions of various noise sources, which we estimated by maximizing model likelihoods with respect to the behavioral data (Methods). The *fdbk* model best fitted the pooled data for both mice (Fig. 7C) and rats (Fig. 7D). We were unable to distinguish between models for most individual mice due to small sample sizes, particularly in parameter regions of high *N_tot_*. However, the ~ 10 × larger rat datasets conclusively favored models with Weber-Fechner scaling (*ffwd* or *fdbk*) for all individuals, with all but one rat favoring the *fdbk* model. Overall, the behavior is poorly explained by exclusively integration noise or Weber-Fechner scaling, instead favoring models that involve a mixture of noise effects.

As discussed, the difference between the *ffwd* and *fdbk* models is in how quickly Weber-Fechner scaling starts to dominate with increasing *N_tot_*. Fig. 7E shows the pooled mouse data compared to predictions of the four models, where the *fdbk* model differs from the *ffwd* model in predicting a more constant asymptotic trend. For rats the *fdbk* model matches a slightly non-monotonic performance trend (Fig. 7F), which is mathematically impossible for any of the other models. A circuit architecture with feedback modulations of sensory responses thus best explains the behavioral data, as well as being the only model considered here that explains count dependence of cue-locked amplitudes.

## Discussion

Psychophysics-motivated evidence accumulation models (Ratcliff and McKoon 2008; Stone 1960; Bogacz et al. 2006) have long guided research into how such algorithms may map onto neural activity and areas in the brain. A complementary, bottom-up approach starts from data-driven observations and formulates hypotheses based on the structure of the observations (cf. (Shadlen et al. 1996; Wimmer et al. 2015)). In this direction and as part of a broader project (“BRAIN Circuits Of coGnitive Systems,” n.d.), we exploited the mouse model system to systematically record from and characterize neural activity in layers 2/3 and 5 of six posterior cortical areas during a task involving temporal accumulation of visual evidence. While similar breadths of survey have been made in nonhuman primates performing perceptual decision-making tasks (Siegel, Buschman, and Miller 2015; de Lafuente and Romo 2006), our work differs in the examination of and findings for the earliest cortical areas, starting from V1 (but see (Hernández et al. 2010)).

To first order, the visual cortical hierarchy is thought to process sensory data in order to extract increasingly abstract visual features, although recent work has shown this processing to be modifiable by motor feedback, temporal statistics, learned associations, and attentional control (Roelfsema and de Lange 2016; Gilbert and Sigman 2007; Kimura 2012; Gavornik and Bear 2014; Keller and Mrsic-Flogel 2018; Glickfeld and Olsen 2017; Niell and Stryker 2010; Saleem et al. 2013; Shuler and Bear 2006; Fiser et al. 2016; Haefner, Berkes, and Fiser 2016; T. S. Lee and Mumford 2003; Zhang et al. 2014; Saleem et al. 2018; Makino and Komiyama 2015; Keller, Bonhoeffer, and Hübener 2012; Poort et al. 2015; Li, Piëch, and Gilbert 2004; Stănişor et al. 2013; Petreanu et al. 2012; Romo et al. 2002; Luna et al. 2005; Nienborg Cohen, and Cumming 2012; Yang et al. 2016; Britten et al. 1996). Still, despite such an encroachment on cognitive operations historically ascribed to higher-order areas, few of these studies have concerned perceptual decision-making behaviors, and it is unclear how well they will extrapolate to the latter. One generally prevalent idea is that the information being processed should still be visual in some way, whether via input from the retina, associations that link visual responses to other events, or visual-feature predictions from other parts of the brain. Our findings did not fully follow this prescription, but instead showed novel deviations from a vision-related nature. For example, there was substantial coding for evidence (a cognitive variable not directly tied to the current visual scene) during the cue period, which then declined in the delay period down to chance levels by the end of the trial. Remarkably, evidence information strongly “resurfaced” in all areas during the inter-trial-interval (ITI; Fig. 3A-B), despite a lack of visual correlates or indeed any visual input at all during this period. In fact, there was a large and distinct subset of cells (~25%, see Fig. 2) that coded for multiple abstract quantities like evidence, choice, and reward during the ITI (Fig. 3). Another trial-context-related effect was the factor of ~2 higher accuracy for decoding contralateral (Fig. 3A) vs. ipsilateral (Fig. 3B) evidence in the cue period, whereas in the ITI both were comparable and there was, if anything, higher accuracy for decoding the difference Δ = #*R* − #*L* (Fig. S1J). These phenomena hint at there being possibly two different mechanisms and functions for evidence information during vs. after the trial. They are the most obvious but not the only surprising discovery.

Overall in our data, activity in visual cortex predominantly reflected abstract variables that had no direct correspondences to the current visual stimulus. ≤ 10% of active cells had responses time-locked to the visual cues. The remaining ~90% were sequentially active in place/time, with population activity patterns encoding present as well as past-trial information about evidence, choice, and reward outcome. Although the proportion of cue-locked responses may have been higher had the visual cues been specifically tuned to retinotopic and other (e.g. spatial/temporal frequency) preferences of the recorded areas, our findings nevertheless revealed a rich set of beyond-visual phenomena that pervaded the visual cortices---if anything despite a lack of optimized visual drive. Inter-area differences were also mostly in degree when comparing V1 to secondary visual areas and even the retrosplenial cortex, although the smaller physical extent of secondary visual areas meant that more of the cortical retinotopic space was included in our recordings. Our findings of predominantly beyond-sensory responses in visual cortices are qualitatively different from the strongly stimulus-driven responses in sensory cortices reported in other sensorimotor transformation tasks (Goard et al. 2016; Pho et al. 2018; Runyan et al. 2017).

Although psychophysical models often conceptualize perceptual decision-making as a feedforward process with detection, accumulation, and categorization stages, the neural data here show no clear mapping between neural responses and such a feedforward set of stages. Instead, “downstream” quantities, such as choice and accrued evidence, strongly influenced neural activity in regions as early as V1, including the amplitudes of sensory-like responses (cf. (Nienborg and Cumming 2009)). These neural observations inspired us to propose an alternative multiplicative feedback-loop circuit model. This mathematical model turned out to better predict the psychophysical performance of mice and rats performing pulsatile evidence-accumulation tasks, compared to previous feedforward models. We thus explore below some thoughts on how our findings may fit into more complex and non-feedforward pictures of neural circuits that underlie the Accumulating-Towers behavior.

### The accumulation process may be distributed and include a beyond-sensory role for the visual cortices

One prominent feature in all of our data is the presence of evidence-related signals throughout the behaviorally indicated accumulation period (Fig. 3A-B). These signals have previously been reported in a variety of brain areas downstream of V1 (reviewed in (Brody and Hanks 2016)), yet perturbation studies have tended not to match expectations for an accumulation circuit (but see (Brody and Hanks 2016; Yartsev et al. 2018)). In fact, specifically for the Accumulating-Towers task, a separate optogenetic perturbation study showed that bilateral inactivation of *any* dorsal cortical site had behavioral effects, the details of which differed across areas (L. Pinto et al. 2018). In that study, temporally-specific inactivations induced surprisingly heterogeneous changes in the weighting of evidence vs. place/time, such that even for V1 there was no clear correspondence to its expected function as (only) a source of momentary sensory input. A possible explanation for these neurophysiological and perturbation findings is that the accumulation process is highly distributed, with many brain regions—including sensory cortices—contributing in overlapping but not entirely redundant ways.

One specific proposal for distributed architectures argues that working memory is not a separate and dedicated subsystem of the brain, but rather that all cortical circuits can accumulate information, over timescales that follow a roughly hierarchical progression across brain regions (Hasson, Chen, and Honey 2015; Chaudhuri et al. 2015; Christophel et al. 2017; Sreenivasan, Vytlacil, and D’Esposito 2014). Compatible with these theories, progressing from V1 to secondary visual areas to RSC we observed increasing timescales of cue-locked responses (Fig. 4I-J), increasing strengths of stimulus-specific adaptation (Fig. 5D), and increasing measures of past-trial information (Fig. 3, Fig. 5H). Our results are also compatible with other experimental observations of increasing timescales along a cortical hierarchy (Murray et al. 2014; Runyan et al. 2017; Dotson et al. 2018; Schmolesky et al. 1998), and furthermore show that even regions as early as V1 contain across-trial information about choice, reward, and sensory history, as previously reported in frontoparietal areas (Morcos and Harvey 2016; Scott et al. 2017; Akrami et al. 2018). A gradual increase in timescales across brain areas suggest that each area may contribute incrementally to the persistence of information, and if so, our data support that this process also includes the visual cortices.

### Spatial and decision-related variables as well as visuomotor feedback may all drive choice-specific sequences of neuronal activation

As animals navigate, place cells in the hippocampus are thought to form a cognitive representation of space by virtue of different subsets of cells being active in different locations in the environment (Moser, Kropff, and Moser 2008). This kind of phenomenon in fact describes the bulk of neural activity in all our data. While place preferences have previously been reported for V1 in non-decision-making contexts (Saleem et al. 2018; Fiser et al. 2016), our data more closely resemble observations of PPC in a decision-making context (Harvey, Coen, and Tank 2012; Morcos and Harvey 2016; Krumin et al. 2018). In particular, as mice ran down the T-maze these place-like cells formed sequential activation patterns that differentiated into one of two choice-specific sequences. (Krumin et al. 2018) noted that this could be explained by individual cells having firing fields parameterized predominantly by two spatial factors, the *y* location along the maze and the view angle θ, with small but nonzero gains from including choice as a third factor. Such an ambiguity between θ and choice as descriptions arises because mice execute their navigational choice by controlling θ, so the two variables are necessarily correlated. Our findings are compatible with (Krumin et al. 2018) in that after accounting for θ, choice decoding accuracy from neural activity was small albeit significant (Fig. 3C vs. Fig. 3D). Interestingly however, as discussed below, control experiments where θ was not a factor instead showed substantial choice-related effects (Fig. S2, Fig. S4). In a delayed match-to-sample paradigm, others have also reported sequential neural activity in the posterior cortex that depended on more than visual/spatial factors, in particular a different sequence for each sample stimulus, despite the latter having no visual or spatial correlates during the delay period (Lu and Tank 2018).

Why might neural activity depend on view angle θ ? There are at least three possible interpretations. First, θ determines the visual environment shown to the mouse (Fig. 1B-C). For visual cortical activity, we do expect differences in visual scene to have significant effects, such as angular receptive fields for cue responses (Fig. 5A-B). Second, in navigation θ is also one of the necessary coordinates (*x, y*, θ) for representing the animal’s location in space, which the brain is thought to keep track of internally as well as to correct with sensory input (Moser, Kropff, and Moser 2008). The prevalent dependence of neural activity on (*y*, θ) may thus reflect computations that support spatial cognition, performed by an interactive loop between visual and navigational subsystems of the brain. Third, θ and the eventual choice *C* may together be signatures of a latent decision variable that drives navigational actions. In tasks like ours, mice appear to continuously execute navigational decisions, as inferred from tendencies in how they turn towards the target T-maze arm fairly early on in the trial (Lucas Pinto et al. 2018; Krumin et al. 2018; Morcos and Harvey 2016; Sanders and Kepecs 2012). This suggests that an underlying decision variable may be correlated on a roughly timepoint-by-timepoint basis with the navigational outcome that it drives, i.e. both θ and *C*. One important distinction between these behavioral readouts is that θ is sampled continuously whereas *C* is only known at the end of the trial. If neural activity reflects a decision variable that continuously drives behavior, it can thus be better correlated with the time series θ than the eventual choice *C*, as the latter can only be assumed to be constant (no changes of mind) and binary in value. In this way, a dependence on a latent decision variable can also explain the observed dependencies of neural activity on both θ and to a lesser but nonzero extent *C*.

The multiple interpretations above are all a question of whether θ dependencies originate from the visual sensory input, an internally maintained spatial representation, or a behavior-driving decision variable related to choice. It would in fact be intriguing for all three factors to be present in the activity of the same neural population, as they span distinct types of information all of which are fundamental for navigation: “where does sensory input say I am”, “where does spatial tracking estimate I am”, and “where do I decide I want to go”. However, because internally generated variables can only be *approximately* correlated to behavioral readouts, whereas we have precise information about visual input changes as parameterized by θ, there is an intrinsic and unavoidable difference in the power of any test for these different natures of dependencies.

We directly eliminated these kinds of ambiguities via control experiments where θ was restricted to be zero up to midway in the delay region. Rather than a reduced dependence on choice, we still saw clear choice-specific sequences (Fig. S2A-C), and choice decoding accuracies similar in strength and timecourse to when θ was *not* controlled for in the main experiments (Fig. S2E vs. Fig. 3C). This favors a decision-variable origin for choice-related effects; however, the three factors are not mutually exclusive. We think it likely in the main experiment that all of these cooperate to explain the neural data.

### Information relevant to navigation, sensory evidence, decision, and task history may be widely broadcast including to early sensory areas

In a related evidence-accumulation study, (Morcos and Harvey 2016) described long-timescale information as reflected in the structure of neural state transition probabilities in PPC, resulting in a diversity of neural population activity patterns that only partially decreased towards the end of the trial. Adopting their analysis, we observed qualitatively similar neural state transition structures not only in areas closest to PPC (AM and MMA), but in fact in all recorded areas (Fig. S7). Our main analyses indicate that this diversity of neural states in all areas corresponds to the coding of multiple task-related variables and their history. Our observations of widespread task-relevant information in six pre-selected areas agrees qualitatively with a recent assay of neural coding across the posterior cortex in a vision-based locomotion task (Minderer, Brown, and Harvey 2019). In the latter, mice were rewarded for maintaining a straight trajectory based on optic flow feedback, and neural activity in most of the posterior cortex reflected all of the behaviorally relevant variables (locomotion velocity of the mice, optic flow velocity, and task history). Our work, using a cognitive task that requires accumulation of evidence for decision-making substantially extends the observations of (Minderer, Brown, and Harvey 2019), to now show that the types of information widely represented in posterior cortices can extend far beyond vision and locomotion (functions intimately related to each other), to much more abstract quantities related to cognition such as accumulated evidence, choice, and reward outcome.

Theories of predictive processing as a canonical cortical computation (Keller and Mrsic-Flogel 2018) offer a conceptual framework for why a variety of task-related information should be widespread in the cortex. These models have two general ingredients that pertain to our observations. The first is that inter-area communication occurs in the form of predictions transmitted from one area to another. For example, place-like cells in V1 may reflect predictions for place-related visual features of the T-maze. The second is that top-down inputs modulate locally represented information. This can serve gating functions such as visual attention, as well as implement computations for perceptual inference (Helmholtz, n.d.). For example, evidence/choice/reward-modulations of cue response amplitudes could reflect a combination of raw sensory input with an internal model (“prior”) of the world, forming an experience-guided percept. The prevalence of such phenomena in our data may thus be conceptually related to components proposed by predictive processing models of cortical function.

Although essentially any behaviorally relevant factor may parameterize predictions transmitted to the visual cortices, predictive processing models are specific in that these predictions should still be *about* visual features. However, we also reported phenomena that have no obvious relationships to vision, most obviously, sequential neural activity during the ITI where mice experienced a dark environment (Fig. 2). This could correspond to abstract information like time until the next trial (cf. reward timing in (Shuler and Bear 2006)), but our results go beyond this in showing that ITI activity furthermore encodes evidence, choice, and reward outcome (Fig. 3), with evidence being more of the differenced form Δ = #*R* − #*L* than during the cue period (Fig. 3A-B vs. Fig. S1H). These types of information are suggestively those necessary for task learning and we speculate that their concurrent presence in a local neuronal population may engage plasticity mechanisms beyond the scope of this work. Neural dynamics during the ITI could alternatively (or additionally) serve to maintain this information over long timescales, e.g. for use in the next trial. It is interesting that early sensory cortices may exhibit memory-like functions for cognitive variables, despite these having at most highly abstract relationships to the immediate sensory stimuli. All in all, our observations point to a rich set of cognitive influences that include but are not fully accounted for by predictive visual processing operations.

### Multiplexing of information may utilize orthogonal directions of neural codes

Many previous studies of choice probability (CP) in evidence accumulation have reported positive correlations between CP and the stimulus selectivity of cells (Britten et al. 1996; Celebrini and Newsome 1994; Cohen and Newsome 2009; Dodd et al. 2001; Law and Gold 2009; Nienborg and Cumming 2014; Price and Born 2010; Kumano, Suda, and Uka 2016; Sasaki and Uka 2009; Gu, Angelaki, and DeAngelis 2014). Translated to our task, this means that neurons that responded selectively to right stimuli tended to have increased firing rates when the animal will make a choice to the right. Our data deviates in that highly contralateral-cue-selective neurons can instead be divided into two near-equally sized subpopulations with positive choice modulation (analogous to CP > 0.5) and negative choice modulation (CP < 0.5) respectively (Fig. S3F-G; negative modulations are marginally more prevalent). In a closely related analysis, there are near-equal numbers of cue-locked cells with amplitudes positively vs. negatively modulated by accumulated counts (Fig. 6C). As two simultaneously recorded cells that respond to the *same* visual cue can be *oppositely* modulated (Fig. 5A), these phenomena are not expected from accounts of spatial- or feature/object-based attention in visual processing ((Cohen and Maunsell 2014; Treue 2014); but see (Snyder, Yu, and Smith 2018)). However, our observations are compatible with mixed choice- and sensory-selectivity neural coding reported in other perceptual decision-making experiments (Raposo, Kaufman, and Churchland 2014).

If choice- and count-modulations of cue response amplitudes originate from some form of accumulator feedback, our proposed feedback-loop (*fdbk*) circuit model suggests an interesting possibility. Intuitively, if comparable proportions of sensory units are positively vs. negatively modulated by feedback, the opposite signs of these modulations can cancel out when sensory unit activities are summed as input to the accumulator. In the model, 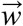 are feedforward weights (“synapses”) of sensory units onto the accumulator, and 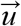 are feedback weights of the accumulator onto sensory units. The condition for cancellation is 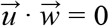, which reduces the accumulator dynamics (Eq. 2) to simple integration: 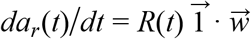. Therefore even with strong feedback connections (|*u_i_*| ≫ 0), the feedback-loop circuit can still be functionally equivalent to pure integration.

The possibility for modulatory effects to cancel out is more general than our feedback-loop model. For a population of neurons, 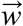 can be thought of as a (linear) readout/coding axis for e.g. sensory information. If the neuronal activities are also modulated along another axis 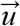, e.g. by choice, then similarly 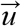 is a coding axis for choice. The coding of these two types of information will not interfere with each other if these axes are orthogonal, 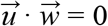, in the sense that choice modulations cancel out when neural activities are projected onto the sensory coding axis. Similar arguments have been made for how motor preparatory activity and feedback do not interfere with motor output (Kaufman et al. 2014; Stavisky et al. 2017), as well as how attentional-state signals can easily be detangled from visual stimulus information (Snyder, Yu, and Smith 2018), and may hint at a general design principle of that allows non-destructive multiplexing of information in the same neuronal population.

### Accumulation may include an amplification process for the sensory signal

Focusing on putatively sensory responses, we found that ~50% of cue-locked cells had amplitudes that depended on choice or accumulated counts in a way that was distinguishable from stimulus-specific adaptation or enhancement (SSA). These responses had two features that were not explainable by a simple model of SSA. First, comparable proportions were best explained by dependence on counts on either the contralateral side, ipsilateral side, or the difference of the two sides (Fig. S3I; cf. (Scott et al. 2017)). For (say) right-cue-locked cells, #*L* or Δ dependencies cannot be explained by SSA. Second, the remaining time-independent #*R* dependencies also cannot be explained by SSA, unless SSA has an infinitely long timescale. However, this would require some additional prescription for “resetting” the adaptation factor e.g. at the start of each trial, because otherwise amplitudes would continue to decrease/increase throughout the ~1 hour long session. Thus while 36% of cue-locked cells gradually adapted to repeated stimuli as might be explained by single-cell mechanisms like fatigue (Grill-Spector, Henson, and Martin 2006), another 50% showed potentially interesting accumulator-related modulations.

The above observation led us to propose the *fdbk* circuit model, where feedback from an accumulator acted as a dynamic gain on sensory responses and thus qualitatively explained count-modulations of cue-response amplitudes. Noise that could arise in multiple components of such a circuit furthermore explained how the psychophysical data deviated from the Weber-Fechner Law at low counts. It has previously been argued that (pure) Weber-Fechner scaling indicates that memory-level noise must dominate over that associated with individual items (Gallistel and Gelman 2000). Instead, we have found in pulsatile evidence accumulation tasks that the behavioral data and *fdbk* model both suggest that multiple significant sources of noise are at play, corresponding to different regimes of high vs. low counts.

The *fdbk* model described the effective structure of the computational circuit, but did not depend on implementation details like anatomical location. Both the sensory and integration stages could in fact be present within a local neuronal network, since our recordings contained cells with sensory-like responses to cues intermingled with others that contain long-timescale information about accumulated evidence. The biological implementation of multiplicative feedback may be an input-output property of individual neurons (Peña and Konishi 2001; Silver 2010), which for pulsatile inputs may be a threshold-linear as opposed to truly multiplicative operation. Alternatively, multiplication can be a network computation, for example involving an intermediate population of inhibitory neurons (Olsen et al. 2012; Atallah et al. 2012; Wilson et al. 2012; Zhang et al. 2014; Fu et al. 2014; Pi et al. 2013; S. Lee et al. 2013; S.-H. Lee et al. 2012). Future experiments will be of interest to establish whether such a circuit structure exists and has a causal role.

To summarize, we have performed a detailed assay of neural phenomena in V1 vs. neighboring posterior cortices, with an aim of better understanding this most basic yet under-explored node of brain areas involved in perceptual decision-making. One long-standing postulated function of the visual cortical hierarchy is to generate invariant visual representations (DiCarlo, Zoccolan, and Rust 2012), e.g. for the visual cues regardless of viewing perspective or placement in the T-maze. On the other hand, predictive processing theories propose that visual processing intricately incorporates beyond-sensory information in a continuous loop of hypothesis formation and checking (Keller and Mrsic-Flogel 2018). Rather than invariant and mostly visual responses, we discovered the bulk of neural activity—even responses to cues—to be strongly influenced by navigational, decision-making and task history variables. Some features of our data are also not fully predicted by predictive processing models. As the visual cues are randomly located per trial, at most their general statistics may be predicted using these cognitive variables, and curiously there was sustained information about the latter even outside of a visual processing context, i.e. the inter-trial-interval. To close the scientific-method loop, our observation of count-modulation of cue-response amplitudes inspired an alternative model of the accumulation process, where input sensory signals are dynamically amplified by the accumulated counts. While this multiplicative feedback-loop architecture results in suboptimal perceptual discrimination compared to simple integration (Fig. 7B), it may reflect other behavioral and engineering pressures that brains have evolved to handle. For example, amplification may be important for selecting a tiny relevant signal out of a massive amount of sensory data regarding the world, and perhaps also out of a massive amount of other neural activity in the brain itself.

## Acknowledgements

We thank B.B. Scott for brainstorming and feedback on the concept of this paper, as well as L. Pinto, C.M. Constantinople, A.G. Bondy, M. Aoi, and B. Deverett for useful and interesting discussions. B. Engelhard and L. Pinto built rigs for the high-throughput training of mice, and S. Stein helped in the training of mice in this study. B. Engelhard and L. Pinto contributed behavioral data from the mouse evidence accumulation task. B.B. Scott and C.M. Constantinople contributed behavioral data from the rat evidence accumulation task. We additionally thank all members of the BRAIN COGS team, Tank and Brody labs. This work was supported by the NIH grants 5U01NS090541 and 1U19NS104648, and the Simons Collaboration on the Global Brain (SCGB).

## Author Contributions

SAK performed the experiments, data analysis and conceptualization/modeling. SYT and DWT designed the experimental setups. SAK wrote the manuscript with input from DWT and CDB. SAK, DWT, and CDB conceived the project.

## STAR Methods

### Experimental Model and Subject Details

All procedures were approved by the Institutional Animal Care and Use Committee at Princeton University and were performed in accordance with the Guide for the Care and Use of Laboratory Animals (National Research Council et al. 2011). We used 11 mice for the main experiments (+4 mice for control experiments), aged 2-16 months of both genders, and from three transgenic strains that express the calcium-sensitive fluorescent indicator GCamp6f (Chen et al. 2013) in excitatory neurons of the neocortex:

- 6 (+2 control) mice (6 male, 2 female): Thy1-GCaMP6f [C57BL/6J-Tg(Thy1-GCaMP6f)GP5.3Dkim/J, Jackson Laboratories, stock # 028280] (Dana et al. 2014). Abbreviated as “Thy1 GP5.3” mice.
- 5 (+1 control) mice (3 male, 3 female): Triple transgenic crosses expressing GCaMP6f under the CaMKII α promoter, from the following two lines: Ai93-D; CaMKII α-tTA [IgS5^tm93.1(tet0−GCaMP6f)Hze^ Tg(Camk2atTA) 1Mmay/J, Jackson Laboratories, stock #024108] (Madisen et al. 2015); Emx1-IRES-Cre [B6.129S2-Emx1^tm1(cre)Kri^/J, Jackson Laboratories, stock #005628] (Gorski et al. 2002). Abbreviated as Ai93-Emx1” mice.
- 1 mouse (control experiments; female) : quadruple transgenic cross expressing GCaMP6f in the cytoplasm and the mCherry protein in the nucleus, both Cre-dependent, from the three lines: Aİ93-D; CaMKII α-tTA, Emx1-IRES-Cre, and Rosa26 LSL H2B mCherry [B6;129S-Gt(ROSA)26Sor^tm1.1Ksvo^/J, Jackson Laboratories, stock #023139].

Mice were randomly assigned such that there were about the same numbers of either gender and various transgenic lines in each group (main vs. control experiments).

### Method Details

#### Optical window implantation surgery

Young adult mice (2-3 months of age) underwent aseptic stereotaxic surgery to implant an optical cranial window and a custom lightweight titanium headplate (~lg CAD design files available at https://github.com/sakoay/AccumTowersTools.git) under isoflurane anesthesia (2.5% for induction, 1-1.5% for maintenance). Mice received one pre-operative dose of meloxicam subcutaneously for analgesia (1 mg/kg) and another one 24 h later, as well as peri-operative intraperitoneal injection of sterile saline (0.5cc, body-temperature) to maintain hydration and dexamethasone (2-5 mg/kg) in order to reduce brain swelling. Body temperature was maintained throughout the procedure using a homeothermic control system (Harvard Apparatus). After asepsis, the skull was exposed and the periosteum removed using sterile cotton swabs. The brain hemisphere for the implant was selected by roughly pre-screening the site for lesser vasculature while the cranium was maintained in a translucent state under a drop of saline. The skull was then dried and leveled using a stereotaxic alignment indicator. The coordinates 2mm caudal, ± 1.7mm lateral to bregma was marked by scoring a ‘V’ with a sterile insulin syringe and then touching a surgical pen to the fringes, allowing the ink to run down the mark. After re-wetting the skull under a drop of saline, a photo of the marked stereotactic location relative to the superficial vasculature was acquired for later comparison to functionally defined visual areas (later section).

A 5mm diameter craniotomy was made using a pneumatic drill (Midwest Carbide Bur No. 1/4, part number 389201) roughly centered around the above marked location, typically spanning the parietal bone on the anterior-posterior sides and partly crossing over the superior sagittal sinus vein. The skull was irrigated with saline throughout this procedure to reduce heating, and the drilled groove was occasionally measured against a reference 5mm diameter steel ring to ensure that the inner edge of the groove was no smaller than the outer circumference of the ring. The cranial window implant consisted of a 5mm diameter round #1 thickness glass coverslip (Warner Instruments) bonded to a steel ring (0.5mm thickness, 5mm diameter, SS316 ring, Ziggy’s Tubes and Wires, Inc.) using a UV-curing optical adhesive (NOA 81, Norlund Products). The implant was lowered glass-side-down into the craniotomy using a custom stereotaxic holder with magnetic prongs (block and rod magnets from K&J Magnetics), leaving about 50-300 μ*m* of the steel ring above the surface of the skull. This served to keep the brain tissue under some pressure, improving stability for imaging, as well as made the imaging plane a little more level. Care was taken to not cut off circulation to the sinus vein, and therefore the inclination of the implant had to mostly follow that of the skull. The steel ring was glued to the skull with cyanoacrylate adhesive (3M Vetbond), which was precisely applied using an insulin syringe with a cut-off tip (i.e. removing the taper). Lastly, the cut flaps of the skin were sealed to the sides of the skull using cyanoacrylate adhesive, exposing an area spanning the parietal bone of both hemispheres plus part of the inter-parietal and frontal cranial bones. Shallow divots were drilled over the surface of the exposed cranium in order to roughen it for better adhesion, and a titanium headplate was attached using dental cement (Metabond, Parkell). The inclination of the skull is about 20° from horizontal, and the inclination of the headplate was kept around 7°-10°. This is a compromise between an increased difficulty for mice to behave in the head-fixed virtual reality (VR) system when the headplate (and by extension their head) is too highly inclined, and the difficulty of imaging from the preparation due to clearance issues when the optical window is too highly inclined relative to the headplate.

#### Behavioral task and training

After at least three and typically five or more days of post-operative recovery mice were started on a water restriction and the Accumulating-Towers training protocol previously described in (Lucas Pinto et al. 2018) and summarized here. Mice received 1-2mL of water per day, and in case of clinical signs of dehydration or body mass falling below 80% of the pre-operative value received supplemental water and edible treats until recovered. All mice were group housed, extensively handled and allowed to socialize in an enclosed enriched environment outside of experimental sessions before being returned to the vivarium at the end of each day. Behavioral training occurred either in dedicated training rigs or similar microscope-equipped setups. Mice were head-fixed so that they could comfortably sit or move on an 8-inch Styrofoam^®^ ball suspended by compressed air, and ball movements were measured with optical flow sensors connected to an Arduino Due running custom code to transform ball rotations into virtual-world velocity. The VR environment was projected onto a custom-built Styrofoam^®^ toroidal screen with visual field spanning ~ 270° horizontal and ~ 80° vertical, using a DLP projector with refresh rates of either 85Hz or 120Hz and RGB color balance of 0, 0.4 and 0.5 respectively. This virtual environment was generated by a computer running the Matlab (Mathworks) based software ViRMEn (Aronov and Tank 2014), plus custom code that controlled the T-maze structure per trial as well as the progression of mice through the shaping procedure.

For historical reasons, 3/11 mice were trained on mazes that were longer (30cm pre-cue region + 250cm cue region + 100-150cm delay region) than the rest of the cohort (30cm pre-cue region + 200cm cue region + 100cm delay region). The structure of the task was otherwise the same, and we included this data because this difference in lengths was not large and none of our results depended on this detail. In VR, as the mouse navigated down the stem of the maze, tall, high-contrast visual cues (Fig. 1B) appeared along either wall of the cue region when the mouse arrived within 10cm of a predetermined cue location. These locations were drawn randomly per trial according to a spatial Poisson process with 12cm refractory period between consecutive cues on the same wall side. Cues were made to disappear after 200ms, although they may fall outside of the field of view sooner depending on the running trajectory of the mouse. Following the notation of (Brunton, Botvinick, and Brody 2013), the task difficulty was set by *γ* ≡ *log*(ρ_*maj*_/ρ_*min*_) =1.2, where ρ_*maj*_ (ρ_*min*_) is the mean density of majority (minority) cues. This corresponded to a mean number of majority:minority cues being 8.5:2.5 for the 250cm cue region maze and 7.7:2.3 for the 200cm cue region maze. Mice were rewarded with 4μ*L* of a sweet liquid reward (10% diluted condensed milk, or 15% sucrose) for turning down the arm on the side with the majority number of cues. The volume of this reward was increased for some mice (e.g. with larger body masses and therefore greater physiological requirements) as well as towards the end of the session in order to encourage sustained high performance. Correct trials were followed by an ITI of 3s in duration, the first second of which the VR display remained on but frozen in place, and the remaining time of which the display was blacked out. Error trials were followed by a loud sound in lieu of a reward and an addition 9s time-out period in the dark. Trial durations were ~11s not counting an additional 9s time-out for error trials, with the cue region corresponding to ~3.5s.

To discourage a tendency of mice to systematically prefer to turn to one side, we used a de-biasing algorithm that adjusts the probabilities of sampling right-vs. left-rewarded trials as described in detail in (Lucas Pinto et al. 2018). If overall performance fell below 55% as calculated over a 40-trial running window, animals were transitioned to an easy 10-trial block of a maze with cues only on one side to increase motivation; this was sometimes also triggered manually by the experimenter. These trials formed a small fraction (~ 8%) of the entire dataset and were included in the analyses in order to maintain, insofar as possible, temporal continuity of the data. Behavioral training and imaging sessions lasted for around 1 to 1.5 hours, during which mice typically completed 150-250 trials. Per session, we computed the percent of correct choices using a sliding window of 100 trials (or the maximum number of trials in that session, in highly atypical cases where this fell below), and included the dataset for analysis if the maximum performance was ≥ 65%.

#### View-angle-restricted control experiments

We trained 4 mice on a control task where the view angle was restricted to be exactly zero in just the cue region (n = 2 mice, 17 sessions), or in the cue region as well as half of the delay region (n = 2 mice, 8 sessions). For the latter variant, the delay region was extended from being 100cm to 170cm long, so that mice had a sufficient amount of time in order to execute turns into the T-maze arm. In the restricted region, only the forward/backward component of the spherical treadmill rotation (corresponding to pitch) is used for advancing the position of the mouse in the virtual world, i.e. *dx*/*dt* ≡ 0 and *d*θ/*dt* ≡ 0. In the unrestricted region, the same rules as in the main experiment (Lucas Pinto et al. 2018) are used to translate treadmill rotations into *x, y* and θ velocities, except that *d*θ/*dt* is controlled to smoothly deviate from zero using an exponential filter. Specifically, the first time the mouse enters the unrestricted region, a view angle velocity gain factor γ_θ_ is initialized to 0 and then updated at every subsequent behavioral iteration using the rule γ_θ_ ← α + (1−α) γ_θ_. The smoothing factor that we found to be adequate for mice to control the virtual world without overly jerky or slow response times is α = δ*T*/(125*ms*), where δ*T* is the treadmill motion sampling interval in milliseconds. Although this paradigm by construction eliminated variations in the visual scene, particularly in where the visual cues appeared, it also removed the visual feedback that encouraged mice to maintain mostly forward movements of the treadmill, analogous to when a human tries to walk down a corridor in the dark. Mice tended to generate larger lateral rotations of the treadmill during the view angle restricted region than in the main, unrestricted experiment (Fig. S2H), which necessitated the above-mentioned exponential smoothing to avoid a sudden panning of the visual scene when control was returned to the mice at the end of the restricted region. This control task was also significantly more difficult for mice to learn and perform, likely because it added a reaction time component to the motor requirements. We attempted to alleviate this by lengthening the delay region and consequently the amount of time the mice could take to turn into the T-maze arms, but that in turn competes with aversive effects of longer trial durations. Nevertheless, we managed to obtain a reasonable dataset using this control task that we consider to be complementary to our data-analysis-based control of visual angle effects for the main experiment. We lowered the required performance level for inclusion in analysis to being ≥ 60% correct trials computed using the same ≥ 100-consecutive trials window.

#### Functional identification of visual areas using widefield imaging

We adapted methods described in (Garrett et al. 2014; Kalatsky and Stryker 2003; Zhuang et al. 2017) to functionally delineate the primary and secondary visual areas using widefield imaging of calcium activity paired with presentation of retinotopic stimuli to awake and passively running mice. We used custom-built, tandem-lens widefield macroscopes to image GCamp6f fluorescence through the optical cranial window implant of the mice that participated in this study. A back-to-back objective system (Ratzlaff and Grinvald 1991) using a combination of either 0.63x, 1x or 1.6x objectives (Leica M-series) was connected through a large filter box holding a dichroic mirror and emission filter. One-photon excitation was provided using a blue (470nm) LED (Luxeon star) through this light path, and the returning green fluorescence passed through the dichroic beamsplitter and was bandpass-filtered at 525 nm (Semrock) before reaching a sCMOS camera (Qimaging, or Hamamatsu). The camera was rotated such that the superior sagittal sinus vein of the mice was parallel to one axis of the square imaging area, which facilitated across-mice registration. The LED was driven by a stabilized power supply and delivered about 2-2.5mW/cm^2^ of power at the focal plane, while the camera was configured for 20-30Hz frame rate and about 5-10μm spatial resolution. To provide software control of the illumination, one rig utilized a mechanical shutter system (Thorlabs) while in the other rig the LED was connected to a MOSFET circuit controlled by custom-written code running on an Arduino Due. The whole system was mounted on custom gantries compatible with our treadmill-based mouse head-fixation system. Visual stimuli were displayed on either a 32” AMVA LED monitor (BenQ BL3200PT), or the same custom Styrofoam^®^ toroidal screen as used for the VR rigs. In both cases, the screen was placed to span most of the visual hemifield on the side contralateral to the mouse’s optical window implant. When using the monitor, the plane of the screen was oriented so as to be roughly centered and perpendicular to the pupil of the mouse’s eye, i.e. at a 30° angle from the anterior-posterior body axis. Due to having to compensate for the inclination of the optical implant, the relative tilt from vertical of the screen was not well-controlled to better than ~ 20°. However, the method for extracting visual area boundaries is highly insensitive to exact placements of the stimulus, as explained below. The space between the headplate and the objective was covered using a custom made cone of opaque material with a soft rubber balloon interface to prevent light from the screen from entering into the imaging system.

The software used to generate the retinotopic stimuli and coordinate the stimulus with the widefield imaging acquisition was a customized version of the ISI package ((Juavinett et al. 2017); from https://sites.google.com/site/iannauhaus/home/matlab-code) and utilized the Psychophysics Toolbox (Brainard 1997). As in (Zhuang et al. 2017), mice were presented with a 20° wide bar with a full-contrast checkerboard texture (25° squares) that inverted in polarity at 12 Hz, and drifted slowly (9° /s) across the extent of the screen in either of four cardinal directions. Each sweep direction was repeated 15 times, totaling four consecutive blocks with a pause in between (induced by the time taken to compute and cache the stimulus pattern). Each block began and ended with 5 seconds each of a delay period where the screen was blanked to gray (50% contrast). The camera was set to continuously acquire a movie of the fluorescence signal throughout the experiment, while the blue LED was controlled by the software to turn on right before the pre-stimulus delay and turn off right after the post-stimulus delay per block. This bright period in the acquired fluorescence movie was later during analysis used to synchronize the timing of the stimulus (recorded at the rate of the screen refresh intervals) to the the frames of the movie. Unlike previous work, the imaging plane was focused on the surface of the brain such that the superficial vasculature remained in sharp contrast. After correction for rigid brain motion (next section; this was negligible in most preparations), the time-averaged fluorescence image was preprocessed using a custom algorithm to identify locations that corresponded to vasculature. This utilized the fact that vasculature both reduced the illumination intensity and absorbed some of the fluorescence of the underlying neural tissue, and thus appeared as pixels that were much darker than the median intensity in a disk of surrounding pixels. The distribution of pixel over median intensity ratio (using the entire image) was used to estimate a threshold for marking a given pixel as vasculature, i.e. if this ratio was two or more standard deviations below the mode of the distribution. For pixels in the middle of large vasculature, this threshold was not very efficient because the median intensity of surrounding pixels was similarly low. The algorithm was thus iteratively applied 5 times, with the median computed using only pixels that had not been marked as vasculature in the previous iteration, which gradually corrects the threshold bias even for vasculature of highly disparate sizes ranging from the sinus vein to ≥ 1 pixel wide tributaries. The size of the disk was also chosen adaptively such that no more than 50% of the pixels within the disk consist of vasculature-marked pixels. The premise behind this algorithm is that the spatial scale over which the cortical retinotopy varies is sufficiently larger than the dimensions of the superficial vasculature (mostly ≤ 10μ*m* in diameter away from the midline). We preferred this as being more well-controlled way of subtracting visible vasculature effects, compared to previous prescriptions (which do not work well for low signal-to-noise samples) of de-focusing the vasculature by placing the imaging plane 0.5-1mm below the surface of the brain.

Retinotopic maps were computed similarly to (Kalatsky and Stryker 2003), with some customization that improved the robustness of the algorithms for preparations with low signal-to-noise ratios (SNR) such as the Thy1 transgenic mice. First, the baseline fluorescence per pixel was estimated using the average intensity in the pre-stimulus and post-stimulus delay periods for the different sweep-direction blocks. To account for possible bleaching of the sample over time, an exponential function was fit to these baseline measurement samples vs. time. An approximate Δ*F*/*F* was then calculated per pixel using the predicted baseline from the exponential fit, and the Fourier first harmonic phase and power of this time series data was computed for each sweep-direction block. When expressed in units of the stimulus bar angle (either azimuthal or altitudinal) relative to the assumed center of gaze of the mouse, this phase of the response corresponds to the retinotopic preference, up to a lag induced by the delayed and negative hemodynamic response. The hemodynamic lag should be the same for both forward and backward sweeps of the stimulus while in contrast the retinotopic response phase reverses together with the reversal of the stimulus, and can in principle be subtracted from the desired retinotopy using this linear relationship. In practice we noticed that mice tended to have asymmetries in the strengths of neural responses to forward vs. backward sweeps, which in low-SNR scenarios lead to a much poorer phase estimate in one of these directions. Reasoning that the hemodynamic lag should vary fairly smoothly across the cortical tissue (excluding explicitly identified vasculature locations), we estimated this lag using only pixels for which the first harmonic power in forward vs. backward sweeps differed by at most 40% from the average power. For the remaining pixels, the median lag in a disk of surrounding pixels was assumed. The retinotopy of a given pixel was then calculated as the average response phase after removing this lag from both forward/backward measurements.

Boundaries between the primary and secondary visual areas were detected using a gradient-inversion-based algorithm described in (Garrett et al. 2014) and for which Matlab-based code was publically available. We again made some changes to improve the stability of this algorithm for samples with a diverse range of SNR values. The azimuthal and altitudinal retinotopy maps obtained above were first corrected for the explicitly identified vasculature locations by replacing the values in those pixels by the 2D-Gaussian-weighted average (σ = 25μ*m*) value of neighboring pixels, excluding other vasculature-marked pixels. This interpolation was repeated until no unresolved pixels remained. These maps were then smoothed by applying the same 2D-Gaussian-weighted average, and their spatial gradients computed using a noise-robust estimation method (http://www.holoborodko.com/pavel/image-processing/edge-detection/) This information was combined into a single visual field sign (VFS) map, which is defined as being +1 if the cortical retinotopy follows the layout of the physical world, or −1 if the cortical layout is a mirror image of the physical world. The VFS of a given pixel was calculated as the sine of the angle between the two gradient vectors, and the rest of the procedure follows closely (Garrett et al. 2014). In brief, the VFS map was thresholded to identify pixels that were above +1.5 or below −1.5 standard deviations of zero, and a series of morphological image processing operations were performed to identify sufficiently large contiguous areas of either sign. Adjoining areas with identical sign were resolved by detecting when the visual field coverage exceeded 1.1 (1 being full coverage), e.g. V1 and area AM both of which had VFS = −1. This defined the visual area boundaries on a per-mouse basis. For illustration purposes only, the average VFS map across mice (Fig. 1D) was computed by first registering together individual maps using a rigid motion correction algorithm. The area boundary detection algorithm was re-run on this average VFS map to produce the borders shown on that figure.

In order to compare the location of the above functionally defined visual areas to previous studies that defined regions by their position relative to skull landmarks, we used the photo of the marked location 2mm caudal, 1.7mm lateral of bregma relative to the surface vasculature of the brain as previously explained (optical window implantation methods). This location was manually annotated relative to the shadow of vasculature visible in the retinotopic maps, and corresponded approximately to the anteromedial corner of the secondary visual area AM.

#### Two-photon cellular-resolution imaging during VR-based behavior

The custom virtual reality plus two-photon scanning microscopy rig used in these experiments follow a design previously described in (Dombeck et al. 2010). The microscope was designed to minimally obscure the ~ 270° horizontal and ~ 80° vertical span of the toroidal VR screen, and also to isolate the collection of fluorescence photons from the brain from the VR visual display by enclosing the light path in a light-proofed aluminum shell except for the access required for the objective lens and laser light entry. Two-photon illumination was provided by a Ti:Sapphire laser (Chameleon Vision II, Coherent) operating at 920nm wavelength (940nm for the one mouse with nuclearly expressed mCherry protein), and fluorescence signals were acquired using a 40x 0.8 NA objective (Nikon) and GaAsP PMTs (Hamamatsu) after passing through a bandpass filter (542/50, Semrock). The amount of laser power at the objective used ranged from ~ 40*mW* for superficial cortical layers to no more than 150mW for the deepest acquired samples (500–550μ*m*). About 1.5 times more power was used for imaging the Thy1 strains of mice than those with Ai93-D lineage. Prior to imaging, the region between the base of the objective lens and the headplate was shielded from external sources of light using a black rubber tube cut from a black latex balloon. This rubber tube was permanently glued to a silicone ring (Krazy Glue) and the ring itself removably attached to the titanium headplate with silicone elastomer (Body Double, Smooth On Inc.) prior to each imaging session. The diameter of the rubber tube matched that of the objective lens close to the silicone ring and later widened so as to fit loosely over the entire objective lens assembly, allowing for flexible vertical and to some extent horizontal movements of the sample relative to the imaging focal plane. Horizontal scans of the laser were performed using a resonant galvanometer (Thorlabs), resulting in a frame acquisition rate of 30Hz and configured for a field of view of approximately 500 × 500μ*m* (512 × 512 pixels) in size. Microscope control and image acquisition were performed using the ScanImage software (2015 versions and beyond; (Pologruto, Sabatini, and Svoboda 2003)).

A single field of view at a fixed cortical depth and location relative to the functional visual area maps was continuously imaged throughout the 1-1.5 hour behavioral session. This was identified by first focusing the two-photon microscope close to the surface of the brain (layer 1), where the shadow of vasculature could be clearly seen and matched to those in the widefield retinotopic map. This vasculature pattern was thus used to locate a two-photon imaging field of view (FOV) within a visual area (or retrosplenial cortex) of interest, and then the imaging plane is lowered to the desired depth below the dura. Due to this two-photon FOV being comparable to or even larger in size than the extent of any one secondary visual area, there can be some ambiguity that we do not claim to resolve when labeling neural data as being approximately from one of the four secondary visual areas. Nevertheless, there is a fairly robust distinction between V1, secondary visual areas, and retrosplenial cortex (RSC). We identified imaging depths as corresponding to layer 2/3 vs. layer 5 based on the typical diameter of neurons at a fixed zoom, which was visibly larger for layer 5 and also with much fewer horizontal processes. This corresponded approximately to a depth of ≥ 350*um* for layer 5, and it is likely that much of our recordings originate from layer 5a due to reduced visibility at high depths. Layer 4 appeared very sparsely labeled for the transgenic mouse lines that we utilized, but there can be some ambiguity that we do not resolve regarding the presence of layer 4 neurons in either category of data labeled as layer 2/3 or layer 5. We altogether analyzed 145 datasets, with ≥ 6 sessions (median 10) and from ≥ 3 mice (median 6) per area and layer (Table S1, Table S2), after a requirement that there be at least 100 consecutive trials per session where the mouse made ≥ 65% correct choices (overall performance per mouse is > 67%, mean 70%; Table S2).

Data related to the VR-based behavior were recorded using custom Matlab-based software embedded in the ViRMEn engine loop, e.g. the treadmill velocity, position in the virtual world, stimulus and trial generation parameters, and so forth. This VR simulation software was run on a separate PC (Windows 7 operating system) than the computer that ran the ScanImage acquisition software. In order to synchronize the behavioral data with the fluorescence imaging frames, we utilized the I2C digital serial bus communication capabilities of ScanImage (version 2015 and beyond) to transmit from the VR computer an identifying timestamp per VR iteration (block, trial, and iteration number). An NI SCB-19 breakout box was used to connect two digital output lines from a NI-DAQ card (62xx or 63xx series) in the VR computer to the FPGA utilized by Scanlmage. The Scanlmage system acts as the 12C slave and the VR computer as the 12C master, and a unidirectional variant of the protocol was implemented in order to be able to emulate this using a NI-DAQ card. Custom C++ code (MEX program) was used to translate the desired synchronization message to the binary 12C format and perform buffered digital I/O (2048-bit buffer) with the NI-DAQ card, using a lightweight execution thread so as not to block Matlab computations. This call was benchmarked to take at most a few tens of microseconds and was negligible relative to the VR iteration duration, which was locked to the refresh rate of the projector (85Hz in the imaging rig although small deviations could occur due to the non-real-time nature of the Windows operating system). There is a stereotyped lag of ~ 2*ms* between transmission and reception of the 12C message, likely due to overhead on the end of the NI-DAQ card driver/hardware. This is also small compared to the 30Hz imaging frame rate. Because the VR iterations were nearly a factor of 3 higher than the imaging frame rate, up to 3 VR time-stamps could be recorded per imaging frame. When analyzing behavioral quantities corresponding to a given imaging frame, only the first associated VR iteration is considered. This, together with potential software and projector refresh lags, makes it unlikely that e.g. the timing of visual stimuli are known to better accuracy than a few tens of milliseconds. This is however of the same order as uncertainties caused by the relatively slow dynamics of the calcium indicator GCamp6f.

#### Data pre-processing and cell finding

All imaging data were first corrected for rigid brain motion by using the Open Source Computer Vision (OpenCV) software library function *cv* :: *matchTemplate*. This correction was performed in 3000-frame stacks, which were used to compute a median fluorescence image that was used as the template to which the individual frames should be registered. Because the median is sensitive to the finite (integer) resolution of the imaging data, which is particularly a problem for low-signal samples, this median was computed on a 10-frame binned average of the data (i.e. the median of 300 temporally averaged frames). The template-matching algorithm located the optimal row and column shifts per frame to maximize the normalized cross-correlation coefficient of that frame with the median-based template. 30 pixels on both x and y extents of the median image were cropped to define the template, thereby allowing up to a 30-pixel shift. Sub-pixel registration was performed using Gaussian interpolation to locate the optimal shifts. The image stack was then corrected by applying these optimized shifts, and the entire procedure was repeated at most 5 times or until the obtained shifts were no greater than 0.3 pixels, whichever occurred first. 2-3 iterations was typical for data in our hands. All 3000-frame stacks that comprised the imaging session were then registered together by applying the same motion correction algorithm to the final template per stack.

For a given imaging dataset, fluorescence timecourses corresponding to individual neurons were extracted using a deconvolution and demixing procedure that utilizes the Constrained Non-negative Matrix Factorization algorithm (CNMF; (Pnevmatikakis et al. 2016)). This algorithm hypothesizes that neural activity is sufficiently well-described as a time-independent spatial component (weights in a small number of pixels) multiplied by a single timecourse (temporal component). The timecourse is furthermore assumed to correspond to a temporally sparse approximate-spiking signal with a calcium impulse response modeled using a low-order autoregressive process (typically 2). The problem is computationally hard and solved using an iterative approach analogous to coordinate-descent methods, where the spatial components are first hypothesized, then the temporal components optimized while keeping the spatial component fixed. This is then used to further refine the spatial components, and the procedure is iterated another time. Because the problem is highly non-convex, the results are sensitive to the initial hypotheses for spatial components. We used a custom, Matlab Image Processing Toolbox (Mathworks) based algorithm that estimated this initialization in a data-driven way i.e. without strong assumptions on the roundness of components or requiring the user to hypothesize the number of cells to be found, albeit with other sample-insensitive parameters related to the spatial scale of the image and SNR thresholds. This alternative initialization procedure is outlined below and may be incorporated in future updates of the CalmAn software suite (Giovannucci et al. 2018).

To improve SNR and reduce processing time, for the initialization stage only the data is downsampled by a factor of 10 in time (to 3Hz) by computing the time-average fluorescence value in disjoint groups of 10 frames. For the *i^th^* pixel we then compute the mode 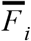 of the distribution of fluorescence values (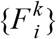 where *k* indexes the imaging frame), and estimate the standard deviation 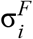 of this distribution using the full-width-at-half-max (FWHM), which is less sensitive to long tails in this distribution as would be present if the pixel corresponded to some part of a neuron that was not always active but had high enough fluorescence values when it was active. The matrix of 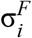 was smoothed to reduce noise and incorporate prior expectations that a neuron should span more than just one pixel, using the guided filtering method (*imguidedfilter* function; see (He, Sun, and Tang 2013)), which better preserves the boundary between neurons and neuropil-only pixels. The rest of the initialization algorithm uses the significance-transformed data, 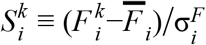, which acts as a whitening filter as well as being expressed in units that allow for thresholds to be specified in a way that scales naturally with the SNR of the sample. In order to parallelize the procedure, we first divided the field of view up into smaller contiguous regions as follows. First we computed the high-tail fraction matrix as the fraction of time-points per pixel where the fluorescence significance exceeded 5 standard deviations, capping this at the 95% quantile to reduce sensitivity to anomalously active cells. Edges were identified in this matrix by finding the zero-crossings of the Laplacian of Gaussian filtered map (*edge* function; (Marr and Hildreth 1980)), with threshold zero so that the edges always formed closed contours. The interior of these contours were filled in using a morphological hole-filling operation (*imfill* function), producing a binary-valued matrix. Contiguous nonzero regions of this matrix mostly corresponded to entire individual cells, or subsets of cells that were adjacent to or overlapped with each other. These regions were processed in parallel to resolve the (potentially overlapping) sets of pixels that corresponded to the spatial footprint of individual cells, as described in the next paragraph.

For data restricted to a given processing-region above, the significance time-series 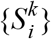 was thresholded above 4 standard deviations after smoothing by taking the geometric average in the 9-pixel neighborhood. The connected components of this 3D binary tensor were then found (*bwconncomp* function), and corresponded to snapshots in time where ideally only one neuron was active (in the case of sparse activity) and occasionally when two or more adjacent neurons were co-active. To resolve the latter, we computed the matrix of pairwise correlations 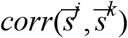 where *j* and *k* indexes pairs of imaging frames, 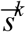 is the vector of 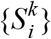 with pixels outside of the connected component set to the negative mean value of pixels inside (to penalize mismatches in extent), and *corr*(…) computes Pearson’s linear correlation coefficient. This matrix typically had an approximate block structure where low values corresponded to frames where one neuron or the other began or ceased to be active, causing the spatial profile to be poorly correlated across frames. When this kind of transition structure was detected, we split up the connected component into subsets of contiguous frames where the spatial correlation remained high, discarding the poorly correlated intermediaries. After identifying all these connected components, a merging/discarding procedure was performed because the same cell could be active at discontiguous points in time, causing it to register as more than one component. The strategy taken here is to handle the most clear-cut cases first and then deal with more ambiguous and edge cases in multiple stages. This part of the procedure uses the spatial shape of each component, defined as the time-averaged spatial profile 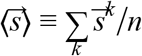 where *n* is the number of frames spanned by the connected component. First, components with highly correlated spatial shapes were merged, by computing the pairwise correlation matrix and thresholding this above 0.8. Cliques in this thresholded matrix were merged into a single component per clique. A subsequent pass first identified ambiguous components as those that were insufficiently large compared to the minimum expected 2D size of cell somata. This was achieved using a heuristic set of morphological operations and includes cases such as small bright spots as well as thin crescent-like shapes, which can occur under a combination of the cell being only slightly active as well as various nonuniformities that can make some parts of the cell appear brighter than others. The same merging procedure was then repeated with two modifications: (1) the pairwise correlation was computed using only pixels within the extent of components marked as ambiguous; and (2) a lower threshold of 0.6 was used to determine which components to merge. Excessively large spatial components were then split up by using morphological erosion operations (*imerode* function) to identify weakly connected “islands”, which occurred when adjacent cells have highly identical activity timecourses. Since this modifies the list of hypothesized components, the procedure of merging components with correlated spatial shapes was repeated with a threshold of 0.7 (still 0.6 for ambiguous components). In the last stage after components have been located in all of the parallel-processed regions of the full dataset, we resolved redundancies caused by e.g. ambiguous boundaries between processing regions by discarding all components that had an insufficient number of pixels that were not overlapping with other components.

In some brain regions, particularly those close to the midline, we have observed that the imaging field of view appears to drift slowly and systematically throughout the ~ 1 hour session. We suspect this to be due to distortion of the brain tissue e.g. from a combination of re-hydration and exercise, which dilates blood vessels. This can cause cells to move out of the field of view, which cannot be correctly modeled by a matrix factorization based source extraction method. As a compromise, we ran the cell finding procedure in approximately 30-minute chunks of the session data (downsampled via averaging to 15Hz to improve SNR and to reduce processing time and memory load), and then identified post hoc components that were the same across these independent reconstructions. This matching was based on the spatial components extracted by the matrix factorization process, which are first centered using image translation to form shape templates 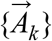 the center-of-masses (COM) of which are all at the same pixel location. We remind the reader that (*A_i_*)_*k*_ for the *k^th^* spatial component is an analog quantity that specifies the intensity of fluorescence in a given pixel *i*. The set of candidate components was initialized using shape templates extracted in the first 30-minute chunk, and then updated using information from temporally subsequent chunks in a serial fashion. For a given new component being examined, all candidate templates the COM of which were within the diameter 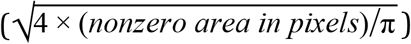 of the new component were considered for potential matches, creating a list {(*c_i_*, *t_i_*)} of potential pairings where *c_i_* indexes a new component and *t_i_* indexes a candidate template. The same components could appear multiple times in this list. The list was then sorted in order of descending 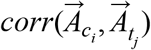 in order to greedily pair the most correlated components first. The first pair in this list was accepted as a match if 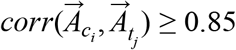, and then all occurrences of *c_i_* and *t_i_* were removed from future consideration. This procedure was repeated until no further matches could be made. The shape templates were updated with the new 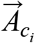 whenever a match was registered, so as to allow for slow systematic changes in the components vs. imaging time. For the purposes of neural data analysis, the time-series data of matched components was concatenated to form a single putative component. This meant that some cells could only be identified in some subset of the imaging session, and the time-points with no information were marked with NaNs. For population-level analyses like the decoding studies, only the subset of cells that were all identified throughout the imaging session were used. For other analyses, care was taken to account for this potential lack of information (i.e. not the same as zero activity levels).

The above procedure for initializing and solving for neuron candidates was by construction very permissive, as the goal was to form hypotheses for as many components as possible including non-somatic sources like transverse and apical dendrites so that these could be correctly demixed by the CNMF algorithm. However the quality of non-somatic components can be poor due to lower SNR, a much larger space of possible spatial configurations, compartment-specific calcium dynamics (incompatible with the matrix factorization hypothesis), higher occupancy leading to pervasive overlap with other components, and high degrees of temporal correlation with e.g. neuropil, and so forth. For the purposes of this work, we are only interested in somatic activity as those can be more definitively localized at a particular cortical depth. We thus performed a semi-automated step of selecting high quality somatic components for analysis, by first classifying components into five morphologically defined classes: (i) doughnuts, which are somatic components that have visible nuclear exclusion in the average fluorescence intensity; (ii) blobs, which are somatic components with no visible nuclear exclusion; (iii) puncta, which are small and typically bright spots that deviate from somata mostly by size; (iv) filaments, which are long, thin, and potentially branching shapes likely corresponding to pieces of dendrites; (v) a catch-all category for shapes that do not fit well into the other categories. We manually classified components for a single exemplary layer 2/3 dataset, and then trained a 30-tree forest of boosted decision trees (Matlab Statistics and Machine Learning Toolbox) to perform this classification based on 6 heuristically defined metrics based on the spatial component 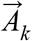 (normalized so that 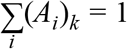) and the corresponding temporal activity signal *T_k_*(*t*):

- The major axis length (largest eigenvalue) of the (*A_i_*)_*k*_ - weighted covariance matrix between {*row_i_*} and {*col_i_*}, the latter being the vectors of row and column indices for the pixels in the component.
- The minor axis length (smallest eigenvalue) of the above covariance matrix.
- The *T_k_*(*t*)-weighted average correlation between the spatial component and the time-series fluorescence data corresponding to nonzero pixels of the spatial component.
- The number of connected components with >1 pixels, after morphologically eroding the nonzero pixels of the spatial component with a disk-shaped structural element of radius 4 pixels. For somatic components this will typically still be one connected component because the erosion of a sufficiently large disk is still a disk. However this can break up dendritic components and completely erase puncta, and so provides discrimination power.
- The fraction of pixels occupied by the morphological skeleton (*bwmorph* function with argument “thin” until unchanged) after dilation with a disk-shaped structural element of radius 2 pixels. This should be close to 1 for dendritic components, but deviates for somata that are not perfect disks.
- The 25% quantile of {(*A_i_*)_*k*_} restricted to pixels within a central region of the component. This region is defined by first computing the convex hull of the component, then eroding that with a disk-shaped structural element of radius 4 pixels. This value is typically high for somatic components, but for components that are curved the center of the convex hull can fall outside of the nonzero region of the component entirely, leading to low values.

This classifier was quite insensitive to the spatial scale of neurons (e.g. layer 2/3 vs. layer 5 data) and performed very well in separating puncta and dendrite-like components from somata, although it did not distinguish well between the doughnut vs. blob categories of somata, and also misclassified some soma-like components (especially when small or oddly shaped, e.g. if half cut off by the field of view) as being in the “unknown” category (v). We manually corrected the classification labels for 10-20% of components, taking care to remove miscellaneous indicators of poor component quality, including those with egregiously drifting baselines, huge overlaps between components, and overly large spatial components that seemed to include neuropil or more than one cell. We paid little attention to trying to separate doughnut- and blob-like components, both of which (and no other category) were accepted for analysis.

All neural data analyses were based on the baseline-subtracted and normalized fluorescence time-series 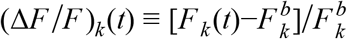 for the *k^th^* putative neuron. *F_k_*(*t*) for a given imaging frame indexed by *t* was defined as the spatially averaged fluorescence value using uniquely assigned pixels i.e. where (*A_i_*)_*k*_ ≠ 0 and (*A_i_*)_*l*_ = 0 for all other components *l* ≠ *k*. The baseline fluorescence 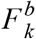 was estimated in 3-minute long windows as the modal value of the distribution of *F_k_*(*t*) in that window. Because of this simple piecewise constant approximation, there can be small discontinuities at the 3-minute boundaries. We did not do use a more complicated (e.g. linearly interpolated or sliding window) correction because the baseline levels of cells in our data are very stable, so this discontinuity is expected to be well within noise levels, and furthermore there is no correlation between the 3-minute boundaries and any particular trial feature, given that the trials have variable durations determined by the mouse’s behavior.

Activity-based cell finding methods like CNMF have a deteriorating efficiency of identifying the more infrequently active cells, which adds ambiguity to our reports of the fractions of cells with various response phenotypes. We therefore restrict all our analyses to a more well-defined subset of cells that have on average ≥0.1 transients per trial (choosing a task-specific measure because of the direction of this work). The noise level 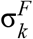 for a given cell *k* is first estimated using the FWHM of the (Δ*F*/*F*)_*k*_(*t*) distribution. A transient is defined as a contiguous block of frames of at least 500ms in duration where 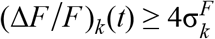, and is assigned to the trial corresponding to the first time-point in this block.

### Quantification and Statistical Analysis

#### Dataset size and mouse strains

All of the two-photon imaging datasets (11 mice, 145 sessions) were included in all neural data analyses. The behavioral accumulator models used data from 17 mice (144366 trials) that performed the Accumulating-Towers task (Lucas Pinto et al. 2018), and data from 6 rats (266984 trials) that performed the flash-accumulation task (Scott et al. 2015).

Unless where explicitly noted, data included all strains of mice listed in the Experimental Model and Subject Details section. As the Ai93-Emx1 strain had higher expression levels of the fluorescent indicator, they produced significantly higher signal-to-noise (SNR) recordings than the Thy1 GP5.3 strain, and contributed more to the layer 5 datasets (see Table S2). As such, some analyses that are sensitive to SNR (e.g. detection of time periods in which cells were active) exhibited small but significant differences, whereas relative quantities such as the proportions of choice-specific and cue-locked cells were less sensitive (Fig. S1B-D, Fig. S3K-M). We believe these small strain differences to be driven mostly by SNR differences as well as the different contributions of layer 2/3 vs. layer 5 recordings as data were pooled across layers for these plots for better statistical power. In short, results from all strains were highly comparable up to numerical differences that were not relevant to our conclusions.

#### General statistics

We summarize the distribution of a given quantity vs. areas and layers using quantile-based statistics, which are less sensitive to non-Gaussian tails. Either the arithmetic mean or the median is used as a central measure, as stated in figure legends. The standard deviation is computed as the difference between the 84% and 16% quantiles of the data points. The standard error (S.E.M.) is computed as the standard deviation divided by 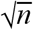 where *n* is the number of data points. For uncertainties on fractions/proportions, we compute a binomial confidence interval using a formulation with the equal-tailed Jeffreys prior interval (DasGupta, Tony Cai, and Brown 2001). The significance of differences in means of distributions were assessed using a two-sided Wilcoxon rank sum test (equivalent to the Mann-Whitney U-test, Matlab function *ranksum*). The p-value threshold for evaluating significance is 0.05 for all tests, unless otherwise stated.

#### Violin plots

The distributions of various quantities are visualized across areas/layers using violin plots, which utilized kernel density estimates of the distributions (Matlab function *ksdensity*) with bandwidth selected using an optimal rule-of-thumb for normal distributions (Scott’s rule, see (Silverman 1986)). This bandwidth was computed separately for the data for a given area/layer, then the average bandwidth was used to construct the kernel density estimate for all areas/layers, for comparability. The width of these plots indicate the estimated density at a given y-axis location. Inset error bars show the S.E.M. as described above.

#### Behavioral metrics

##### Psychometric function

The fraction of trials where a given mouse turned right is first computed in 11 evenly spaced bins centered around Δ = [−15, −12,…, 15], resulting in a list of fractions {*f_i_*} where *i* indexes the bin and the corresponding bin-average {Δ_*i*_}, which can differ from the bin centers depending on the generated distribution of Δ across trials. These data are fit to a 4-parameter sigmoid function *p_R_*(Δ) = *p*_0_ + *B*[1 + *e*^−(Δ−Δ_0_)/*λ*^]^−1^ by minimizing a weighted-least-squares loss function using the Matlab function *fit*:

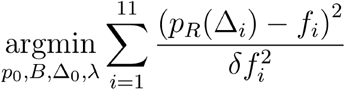

The weights on the data points are given by {δ*f_i_*} being the 68% (one standard deviation) binomial probability content confidence interval on the observed {*f_i_*}.

##### Utilization of evidence

As detailed in (Lucas Pinto et al. 2018), we modeled the dependence of the mice’s choices on cues at various spatial locations along the maze using a logistic regression model where the factors are the amount of evidence (#R − #L cues) in equally-sized thirds of the cue region. The regression was performed using the Matlab function *glmfit* with the data assumed to be binomially distributed with a logistic link function. Statistical uncertainties on the regression weights were determined by repeating this fit using 1000 bootstrapped pseudo-experiments, where the same number of trials were drawn with replacement from the original experiment.

#### Neural coding of task-related information

##### Tosk-epoch time axis

In order to compare data across mice and trials in which the T-maze was traversed at different speeds, we resampled the Δ*F*/*F* time-traces of cells using a task-epoch axis that is piecewise linear in time within fixed segments being the pre-cue period, cue period, delay period, a “turn” period up to the end of the trial, and the ITI. For each trial, we kept track of the imaging frames in which the virtual world y position of the mouse first crossed into the cue region 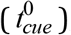, delay region 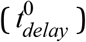, and the arm region (past the stem; 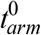), as well as the first frame of the ITI 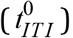, and for error trials only the first frame of the penalty time-out period 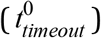. For a given imaging frame *t*, where *t* = 0 (*t* = *t_end_*) corresponds to the start (end) of the trial, the task-epoch time is defined as:

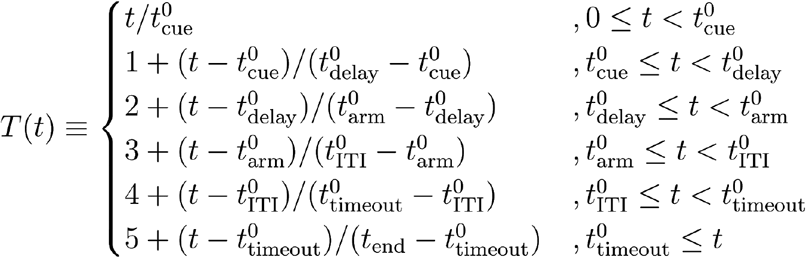

The number of resampling bins per task-epoch segment was selected to result in approximately equal duration bins (~ 200*ms*). For the data in our hands, these were {8,17,12,5,11,28} bins respectively. For display purposes, the duration of a given epoch is shown as the average duration in seconds across all trials in the data.

##### Choice-specific sequences

For comparability, we followed the analysis and presentation in (Harvey, Coen, and Tank 2012), but with adjustments necessary for the faster dynamics of the GCaMP6f indicator as well as to report additional statistics. The following analyses only used data from trials where the mouse made a correct (rewarded) choice. First we located all periods of time in which the trial-average Δ*F*/*F* for a given cell was at least 25% of the maximum for at least 2 epoch bins 400*ms*). The cell was defined as choice-specific if a two-sample t-test detected that the distribution in right-vs. left-choice trials of Δ*F*/*F* in these active periods were significantly different at a 5% significance level. For these choice-specific cells, the side selectivity was defined as the one with the greater distribution average. A ridge-to-background excess was defined using the Δ*F*/*F* averaged over only preferred-choice trials (or all trials if the cell was not choice-specific) as the maximum value in time minus the modal value of the distribution. The mode was computed using a fast algorithm that does not require binned estimates of the probability distribution function (Bickel and Frühwirth 2006), and we generally find it to be a robust and simple-to-calculate estimate of baseline activity for cortical neurons, which are sparsely active. In the extreme theoretical case of a completely inactive cell, the distribution of Δ*F*/*F* values should be approximately Gaussian and the mode of this distribution corresponds to the zero activity i.e. baseline state. If the cell were to be active, given the good contrast of the GCaMP6f indicator and high SNR of our data this contributes some probability mass to a long high tail of Δ*F*/*F* values. Even for highly active cells or trial-averaged data, which is temporally less sparse, the mode remains little shifted (unlike other summary statistics such as the mean) especially since cortical cells have bursty firing patterns that sample a wide dynamic range of Δ*F*/*F* values.

We selected cells to include in the choice-specific sequences analysis by asking if they have sufficiently stereotyped task-related activity. First we generated 1000 null hypotheses per cell by randomly rotating the Δ*F*/*F* time series, which maintained the temporal statistics of the data but broke their correspondence with task-epoch axis. A cell was determined to have significantly task-localized activity (Fig. S1A) if no more than 5% of these randomized pseudo-datasets have a ridge-to-background excess greater than that of the actual data. This test is rather permissive for very sparsely active cells, so long as they were active more than once throughout the entire session and there is some task-related locality to their activity patterns. Complementary to this, we also asked the question of whether cells reliably responded within their putative firing fields (Fig. S1B). The latter is defined using the Δ*F*/*F* averaged over only preferred-choice trials (or all trials if the cell was not choice-specific), by finding all time-points that have average Δ*F*/*F* values being at least 50% of the maximum and selecting the period that is contiguously high and containing the maximum. The reliability index was defined as the fraction of preferred-choice trials in which the activity averaged in this firing field is ≥ 3 times noise, where the noise is estimated as 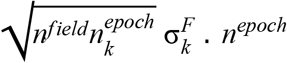 is the median number of imaging frames per epoch bin, 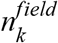 is the number of epoch bins in the firing field of cell *k*, and 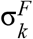 is the estimate of per-imaging-frame noise explained in the previous section. Only significantly task-localized cells with reliability ≥ 20% were included in Fig. 2.

##### Δ-modulation of activity in firing fields

For non-cue-locked cells, we tested for dependence of a given cell’s activity on evidence strength by fitting a linear model that regressed its firing-field-average Δ*F*/*F* against the cumulative value of Δ at the onset of its firing field (example cells in Fig. S1I). This analysis was used only to eliminate significantly Δ-modulated cells from the choice decoding analyses (Fig. 3C-D).

From a purely behavioral standpoint, the strength of evidence is correlated with both the mouse’s eventual choice and the likelihood of it receiving a reward at the end of the trial. It is also correlated with sensory effects that could be caused by differences in view angles at the time that the cell was active. We controlled for all these possible alternative explanations for the neural activity as follows. For choice-specific cells (previous section), this analysis uses only trials that corresponded to the preferred choice for that cell. For non-choice-specific cells, the slope computation described below was first performed separately for trials of each choice, then the average slope across the two choices was taken. Effects arising indirectly from view-angle-specific neural activity was controlled for by weighting the data used in fitting this linear model such that the distribution of θ in that data are equal when conditioned on Δ (Runyan et al. 2017). First we divided the data into five bins defined by Δ ≤ −5, −4 ≤ Δ ≤ −2, −1 ≤ Δ ≤ 1, 2 ≤ Δ ≤ 4, and Δ ≥ 5. We then computed the locations of view angle bin edges [θ^(0)^, θ^(1)^,…, θ^(3)^] such that each bin contained insofar as possible the same proportion of θ values, or equivalently the [0,33.3%, 66.7%, 100%] quantiles. Let 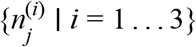 be the counts of data in the *j^th^* Δ bin that fall into these θ bins. We defined the target proportions as the geometric mean across Δ bins, 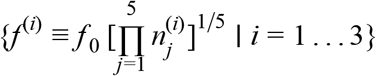, where *f*_0_ is a normalization constant such that 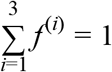. The weight for a given data point was thus 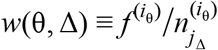, where *j* <Δ is the Δ bin and *i*_θ_ is θ the bin in which that data fell. Importantly, the use of the geometric mean resulted in a weight of zero for bins where one or more of the data categories were unavailable, as this cannot be compensated for via weighting. This caused a loss of statistical power, but this was small because for a fixed choice, there was little dependence of θ on Δ (re Fig. S5E). Lastly, cells that were active in the ITI had no direct view angle dependence since the virtual world display was either frozen in place or later blacked out, but could instead be responding to the delivery or lack thereof of the reward. For these cells we controlled for the reward outcome in lieu of θ using the same distribution-weighting method. To test for modulation by evidence in the previous trial, we analogously controlled for the past reward outcome (only) for all cells. This neglects correlations of the present view angle with the past outcome, but we have in unpublished analyses observed these to be negligible, explainable by that mice predominantly interrupt their motor actions by stopping to lick (or in anticipation thereof) at the end of the trial.

The above-mentioned weighted least-squares linear fit can be computed analytically ((Press et al. 2003) chapter 15.2). Let {*F_k_*} be the average activity in the cell’s firing field in a given trial *k*, {Δ_*k*__} be the corresponding cumulative #R-#L at the onset of the firing field in that trial, and {*w_k_*} be the weights of the data as described above. The slope of the linear fit is 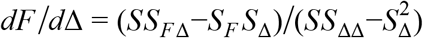, where 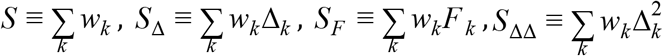, and 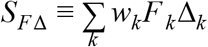. We categorized a cell as being significantly Δ-modulated if *dF*/*d*Δ is larger in magnitude than the same in at least 95% of null hypotheses constructed by permuting the Δ*F*/*F* data across trials and repeating the weighted fits. The increase in significantly Δ-modulated cells throughout the cue period can be due to greater statistical power for regression where a larger range of Δ was sampled, but this does not affect trends from the delay period and beyond.

##### Decoding from neural population activity

For the decoding analyses, we downsampled the Δ*F*/*F* time-series data by averaging in (disjoint) groups of two epoch bins, resulting in ~ 400*ms* duration samples. Behavioral variables to be controlled for, i.e. the view angle in the main experiments and the Arduino sensor velocity in control experiments, were similarly averaged in the epoch bins corresponding to the resulting Δ*F*/*F* time series. For each of this epoch bins, a SVM classifier (Matlab function *fitcsvm*) was trained on 2/3rds of the data (i.e. across trials but restricted to a given epoch bin) and the classification accuracy (proportion of correctly classified test trials) was computed on the remaining 1/3rd of held-out data. This procedure was repeated with the other two disjoint subsets of 1/3rd of held-out data, i.e. using a 3-fold cross-validation paradigm. This 3-fold cross-validation procedure was itself repeated 35 times, each with a different random partitioning of the data into thirds. Thus the average accuracy over a total of 105 cross-validation sets was reported. The SVM classifier used a box constraint penalty factor of 1, a linear kernel function, and the predictors were standardized.

To control for a behavioral variable ψ of interest, we computed weights for the training data input to the SVM classifier so that the distribution of ψ in that data are equal when conditioned on the target quantity to be classified. This weighting procedure is identical to that described above for assessing Δ-modulated cells, except that we used predetermined bin edges [−85°,…,−5°,+ 5°,… + 85°] for when the view angle should be controlled, and the obvious right/left choice categories for when choice should be controlled. The use of fixed-resolution θ bins is important because near the end of the T-maze corridor the view angle distributions for right-vs. left-choice trials necessarily becomes highly non-overlapping as the mouse executes a turn into either arm, as well as spans a large range of possible values, which would have resulted in a large degree of residual differences in view angle distributions had the same quantile-based strategy as above been used. For these same reasons, we were mostly unable to dissociate view angle from choice-related effects past midway in the delay region up to the end of the trial in Fig. 3D, as ≥ 50% of data had to be discarded (received zero weight) due to there not being data in the other choice category with the same view angles. For present and past choice decoding, the view angle was controlled for. For reward decoding the choice that lead to that outcome was controlled for, i.e. the past choice was controlled for when evaluating the past reward decoding accuracy.

The significance of the decoding accuracy was assessed by constructing 100 null hypothesis pseudo-experiments where the Δ*F*/*F* of cells for a given epoch bin were permuted across trials. The entire procedure above was repeated per pseudo-experiment, and the p-value of the decoding accuracy was defined as the fraction of null hypotheses for which the cross-validated decoding accuracy was greater than or equal to that of the actual experiment. The data from each imaging session was analyzed separately. To correct for multiple comparisons when determining whether the decoding p-value for a particular dataset was significant, we used a hybrid Hochberg-Hommel type modification (Gou et al. 2014) to the Bonferroni procedure. For a given type of decoder, say, upcoming choice, we constructed a list of p-values across all datasets. The p-values were sorted in descending order, resulting in a list [*P*_1_, *P*_2_,…] where *P*_1_ is the largest p-value, and the first rank *i*_α_ is found such that *P*_*i*_α__ ≤ α (*i*_α_ + 1)/(2*i*_α_). α = 0.05 being the significance level of this test. All decoding accuracies corresponding to *P_j_* ≤ α/*i*_α_ are considered to be significantly above chance. This correction is likely conservative because given the ~ 1*s* duration of firing fields even just on a per-cell basis (Fig. 2C), there is reason to think that decoders for consecutive time-points (separated by ~ 400*ms*) have positively correlated accuracies, i.e. the number of repeated tests should be smaller than the naive counts.

##### Analysis of view-angle-restricted control experiments

Overall, we observed qualitatively similar neural phenomena in the control experiments where the view angle was restricted to zero throughout the cue region (and where relevant, half of the delay region). For choice decoding, as running speed is known to modulate the response of at least visually tuned cells (Niell and Stryker 2010), we controlled for this by weighting trials such that the distribution of treadmill movement speeds are the same when conditioned on choice (Fig. S2G). This had little effect as mice ran at a fairly stereotyped speeds on each trial (Fig. S2I). On the other hand, the lateral (X only) treadmill velocity did differ between right- and left-choice trials in these control experiments (Fig. S2H-bottom), albeit to a much lesser extent in the main experiments (Fig. S2H-top). This observed difference in motor behavior is likely due to there being no direct visual feedback during the view-angle restricted period, unlike in the main experiment where running at an angle was visually like running “into” the wall. Since the correlation between the sign of the lateral velocity and choice was high for some mice, controlling for it resulted in a substantial reduction of the accuracy for decoding choice from the neural data (Fig. S2J; but not for the RSC dataset, where the mouse had less of a choice vs. lateral-velocity correlation). It is possible that we have discovered motor-action-specific activity in the visual cortices, or a more subtle and motor-action-specific form of sensorimotor-mismatch-driven neural activity than previously reported (i.e. studies where the visual scene was abruptly halted as the mouse was running, see (Keller, Bonhoeffer, and Hübener 2012)). However, X-velocity differences were subtle in the main experiment where we still find above-chance choice decoding accuracies, suggesting a more parsimonious explanation that the neural population reflects the mouse’s choice, and that the latter is itself drives motor actions even without visual feedback during the control task.

#### Impulse response model for cue-locked cells

This analysis excluded some rare trials where the mouse backtracks through the T-maze, because these can have unusual neural effects outside of the scope of the models. Trials were thus included if the *y* displacement between two consecutive behavioral iterations is > −0.2*cm* (including all time-points up to the entry to the T-maze arm), and if the duration of the trial up to and not including the ITI was no more than 50% different from the median trial duration in that session.

We modeled the activity of each cell as a time series of non-negative amplitudes *A_i_* in response to the *i^th^* cue, convolved with a parametric impulse response function *g*(*t*):

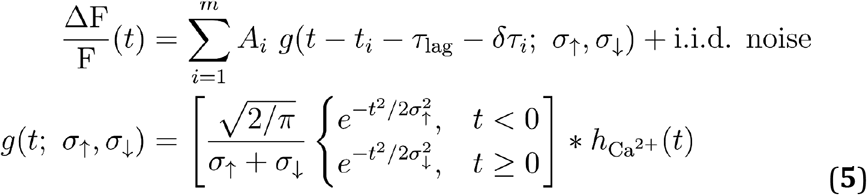

{*t_i_*|*i* = 1,…, *m*} are the appearance times cues throughout the behavioral session. The free parameters of this model are the lag (*τ_lag_*), rise (σ_↑_) and fall (σ_↓_) times of the impulse response function, the amplitudes *A_i_*, and small (L2-regularized) time jitters δτ_*i*_, that decorrelates variability in response timings from amplitude changes, *h*_*Ca*^2+^_(*t*) is a calcium indicator response with parameters from literature (i.e. same for all cells; (Chen et al. 2013)), which deconvolves calcium and indicator dynamics from our reports of timescales. This is parameterized as a difference of exponentials, 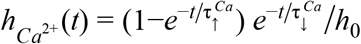, where 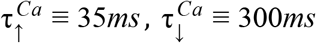 and *h*_0_ is a normalization constant such that the peak of this function is 1. We note that timescales cannot be resolved to better than about the imaging data rate of 1/15*Hz* ≀ 67*ms*.

We maximized the model likelihood to obtain point estimates of all the parameters, using a custom algorithm detailed below. The significance of a given cell’s time-locking to cues is defined as the number of standard deviations that the impulse response model AIC_c_ score (bias-corrected Aikaike Information Criterion, (Hurvich and Tsai 1989)) lies above the median AIC_c_ of null hypothesis models where the timings {*t_i_*} of cues were randomly shuffled within the cue region. In addition, a small fraction of cells responded to both left- and right-side cues. We parsimoniously allowed for different impulse responses to these by first selecting a primary response (preferred-side cues) as that which yields the best single-side model AIC_c_, then adding a secondary response if and only if it would improve the fit (see below for algorithm details). We defined cells to be cue-locked if the primary response significance exceeds 5 standard deviations. Statistics reported in the text include the peak of the response *τ_lag_*, the onset of the half-maximum response, 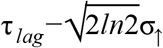, and the full-width-at-half-max (FWHM), 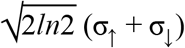.

Because of the nonlinear dependence on a large number of parameters and likely multiple local optima for this problem, we used a custom coordinate-descent-like algorithm (Wright 2015) where subsets of parameters are optimized at a time while keeping others fixed or in simplified forms. The three subsets are the impulse response shape parameters {τ_*lag*_, σ_↑_, σ_↓_}, the amplitude parameters {*A_i_*}, and the time-jitter parameters {δτ_*i*_}. The optimization procedure utilizes the C++ linear algebra package Eigen (version 3; http://eigen.tuxfamily.org/). an Eigen-based implementation of the Broyden–Fletcher–Goldfarb–Shanno (BFGS) algorithm for gradient-based function minimization (https://github.com/PatWie/CppNumericalSolvers), and is presented in Algorithms 1-4, which takes as input the activity data *F*(*t*) for a given cell (simplifying the Δ*F*/*F* notation) and outputs a predicted activity time-trace *m*(*t*). Model selection is performed using AIC_c_. In particular because the model assumes that the data are Gaussian-distributed around the model prediction, this calculation is:

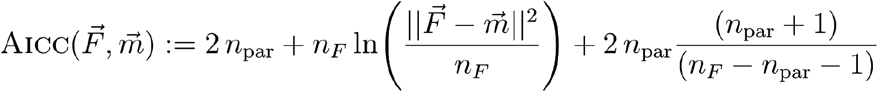

where we have treated the discretely sampled time-series data as vectors, 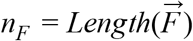, and *n_par_* is the number of free parameters in the model.

If we assume for a moment that the shape and jitter parameters are fixed, finding the optimal amplitudes can be formulated as a linear regression problem 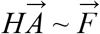, where 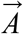 is the *n_A_* × 1 vector of amplitudes, 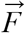 is the *n_F_* × 1 vector of Δ*F*/*F* time-series data, and *H* is an *n_F_* × *n_A_* regressor matrix the columns of which are the same impulse response timecourse but shifted to have peaks located at *t_i_* + τ_*lag*_ + δτ_*i*_ for the *i^th^* column (*t_i_* is the onset time of the *i^th^* cue). We assume there is a function *RegMat* that computes this matrix given the shape parameters and onset times. In order to account for the finite duration of imaging frames, the impulse response shape (Eq. 5) is evaluated by taking the integral within the imaging frame in question (this is particularly important for stability of the fit when the duration parameters are small relative to the frame rate). That is, for the calcium response function in a given imaging frame at [*t, t* + Δ*t*] we compute *h*_*Ca*^2+^_(*t*) ~ *H*_*Ca*^2+^_(*t* + Δ*t*)−*H*_*Ca*^2+^_(*t*) where the cumulative integral function is:

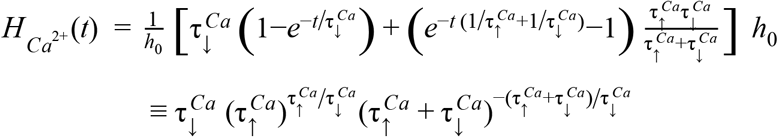

Similarly for the full impulse response function we compute *g*(*t*) ~ *G*(*t* + Δ*t*)−*G*(*t*) where

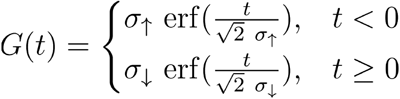

The procedure *OptimShape* (Algorithm 2) assumes zero jitter, and uses gradient descent (BFGS algorithm] to optimize the shape parameters while keeping the amplitude parameters fixed, alternating this with steps where the shape parameters are kept fixed while the amplitudes are obtained using a fast nonnegatively-constrained conjugate gradient solver (Eigen function *Eigen* :: *internal*:: *constrained_cg*). We use L2 regularization to break potential degeneracies in the solution as well as constrain the shape and jitter parameter values to be within a reasonable range given the duration of trials. Parameter bounds are implemented by using a working-set of unbounded parameters that the BFGS optimizer “sees”, which are then transformed using a softplus-like rectification function by *RegMat* before using them to compute the regressor matrix *H*:

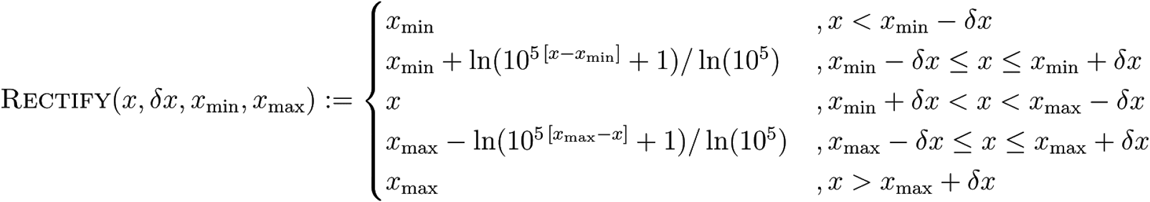

This function transitions quickly but smoothly between the hard bounds [*x_min_,x_max_*] and a linear regime in between. The bounds are very liberally set to [0, 20*s*] for τ_*lag*_, and [0, 10*s*] for σ_↑_ and σ_↓_. The jitter parameters {δτ_*i*_} are bounded to be in the range [−τ_*lag*_, + τ_*lag*_].

The procedure *RefineTimings* is highly similar, except that the shape parameters as obtained from procedure *OptimShape* are held fixed while the jitter and amplitude parameters are iteratively optimized (Algorithm 3). Thus the jitter parameters allow for small variations in timings of either the experimental measurement of the cue presentation times or in the neural responses, but these are restricted degrees of freedom in the sense that they are not allowed to cause deviations in the shape parameters. Lastly, in the case where a cell has two significant response components i.e. to both right- and left-side cues, procedure *RefitA* (Algorithm 4) is called to jointly optimize the two sets of amplitudes (i.e. for right-cue vs. left-cue responses). This more fairly allows the two components to compete in explaining the data, as the secondary component was previously obtained by fitting the model on the residual *F(t)*–*m*(*t*) where *m(t)* is the prediction of the primary response model.

As detailed in Algorithms 2-4, L2 regularization terms are included for the amplitude and jitter parameters. We select the strength of the regularization terms to contribute a fixed small amount to the cost function. For the amplitude parameters, regularization enters the cost function as the second term in *H^T^H* + *λ_A_I*, so to make the regularization effect be comparable to that of the regressor matrix *H*, we should select 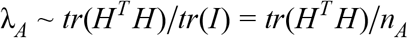. We want the regularization to have a small effect instead, so this motivates choosing *λ_A_* = 10^−3^ *tr*(*H^T^H*)/*n_A_* as in Algorithms 2-4. For the jitter parameters, we ballpark the sum-squared residuals 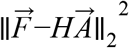 to be of magnitude (σ^*F*^)^2^ per time-point, where σ^*F*^ is the estimated fluorescence data noise for that cell.

**Algorithm 1.**
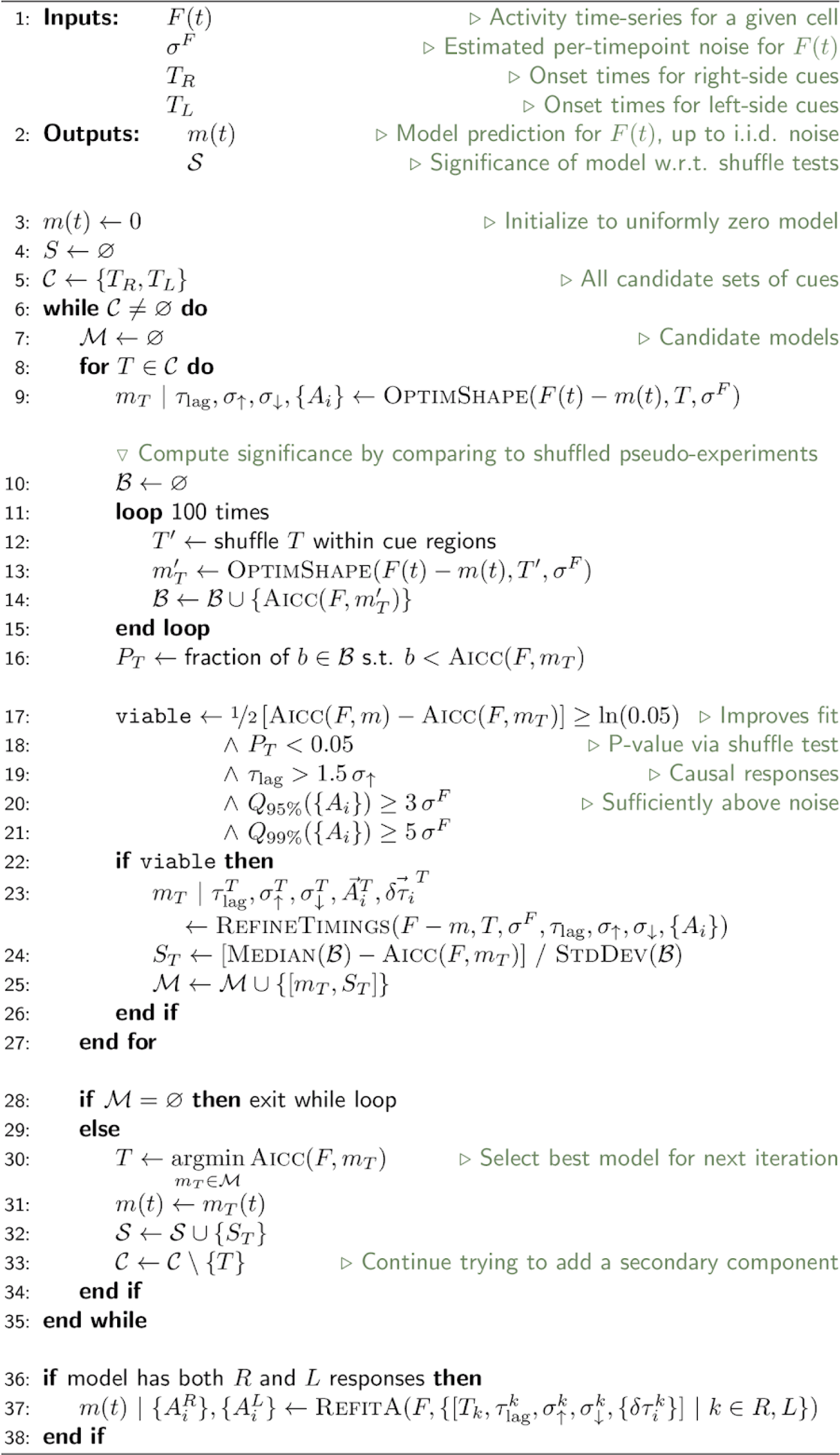
Impulse response model selection.

**Algorithm 2.**
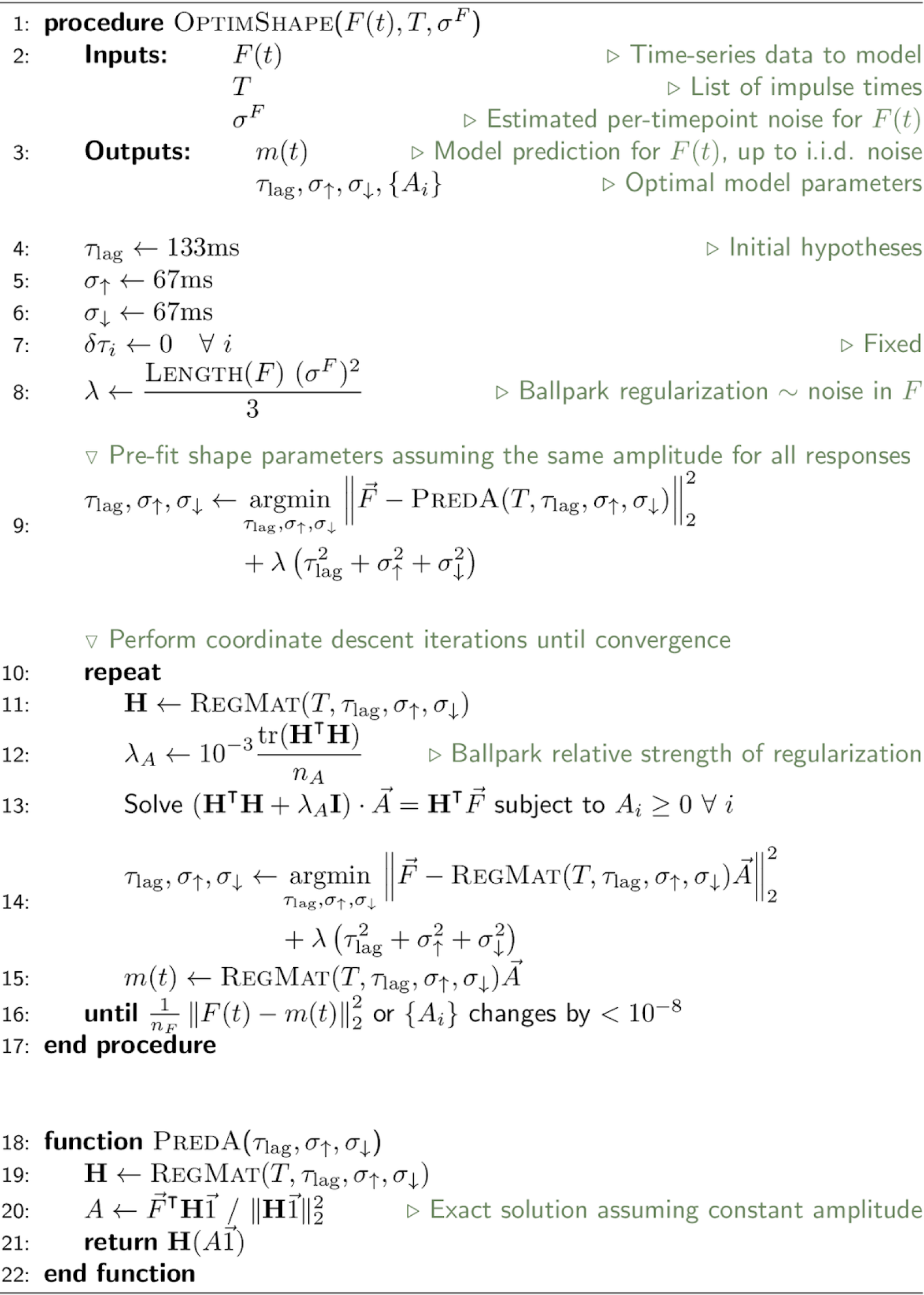
Optimize shape and amplitude parameters.

**Algorithm 3.**
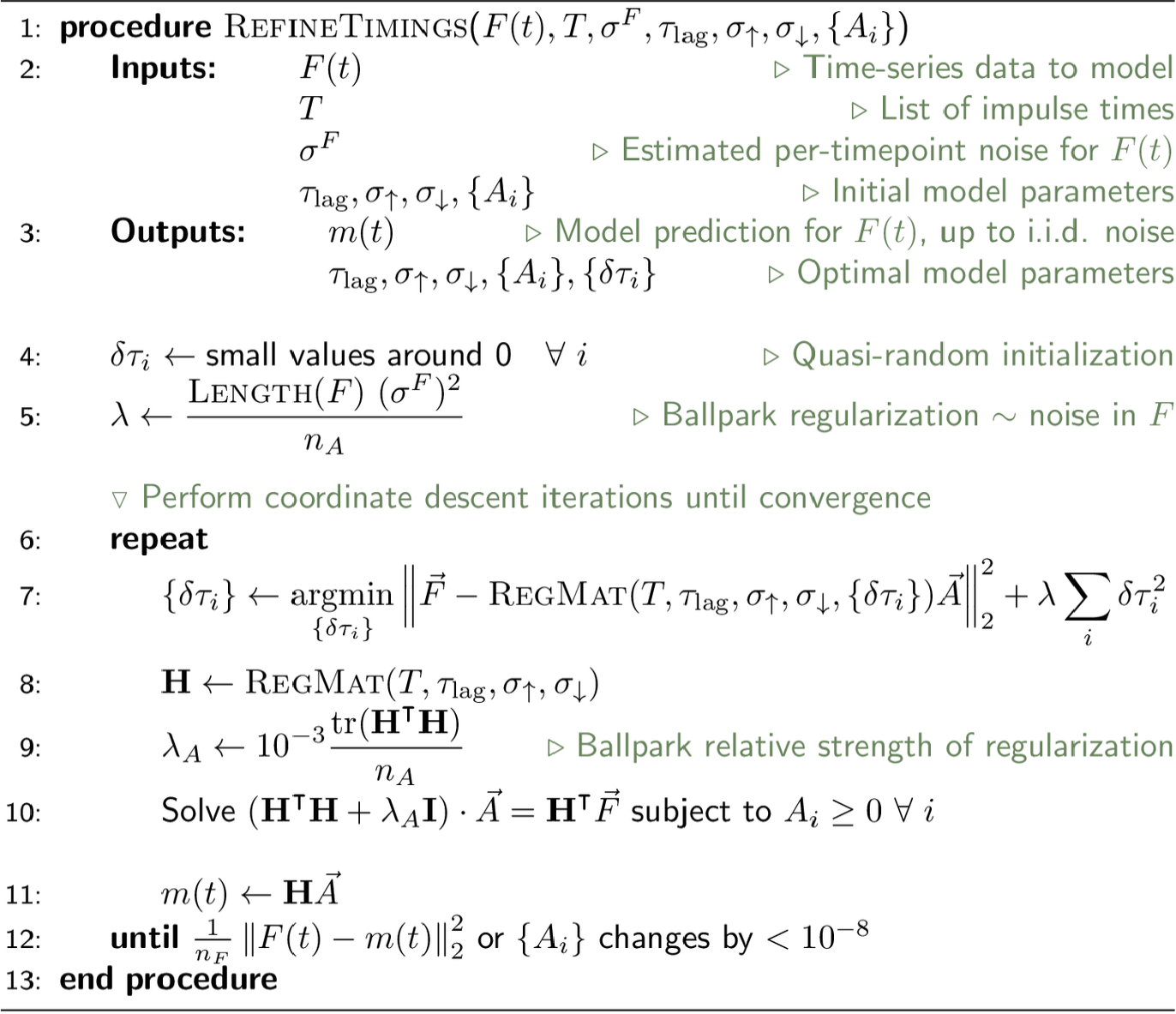
Optimize jitter parameters.

**Algorithm 4.**
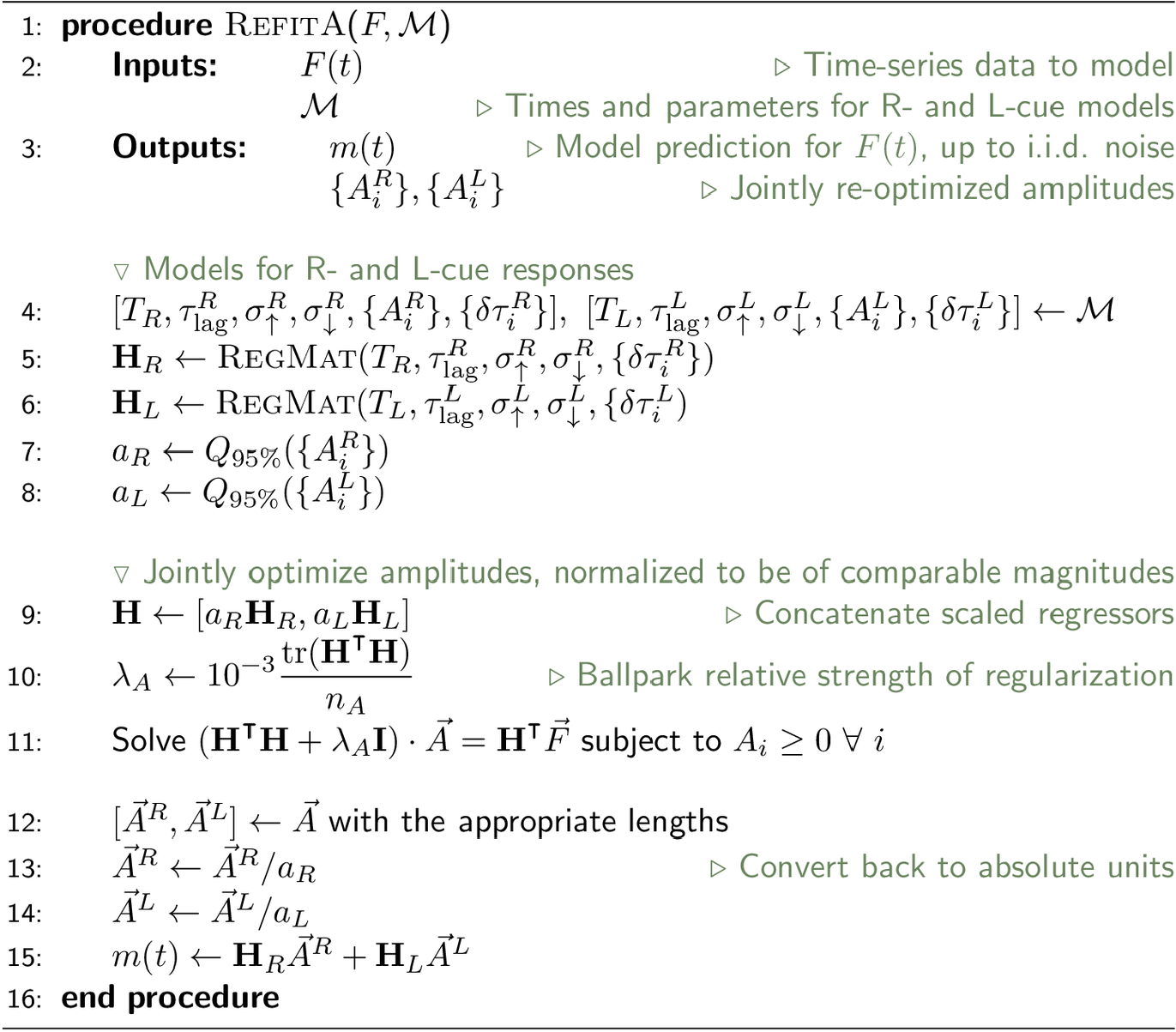
Refit multi-component model amplitudes.

#### Amplitude modulation models

The cue-locked amplitudes are modeled as samples from a Gamma distribution with mean functions as follows:

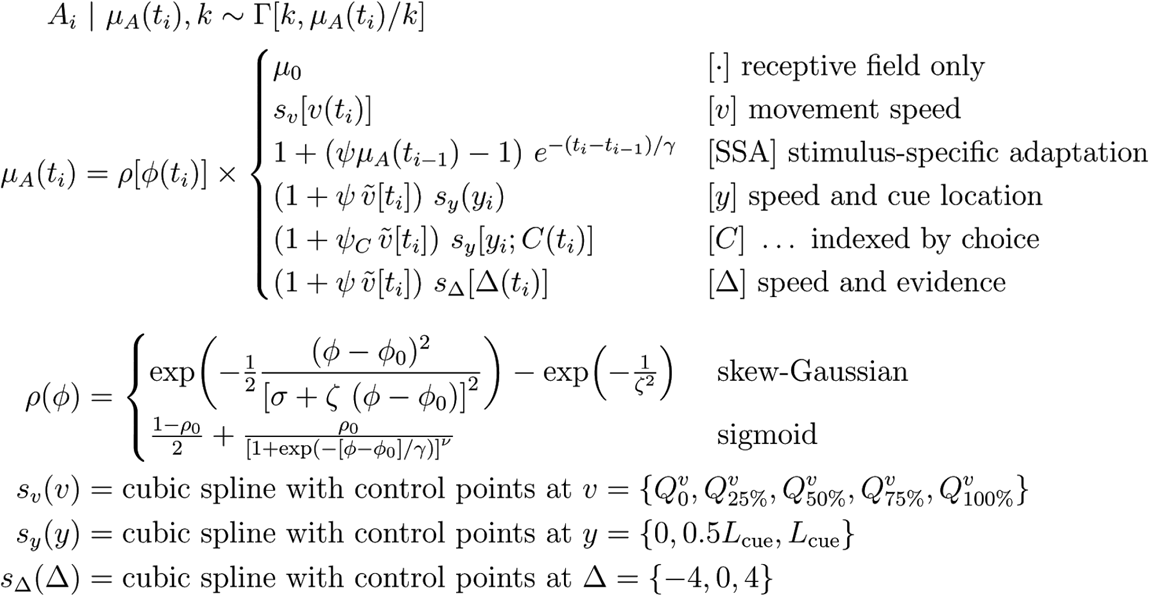

*t_i_* is the onset time of the *i^th^* cue, which is located at distance *y_i_* along the cue region and appears at a visual angle *φ_cue_*(*t_i_*) relative to the the mouse. *C*(*t_i_*) is the upcoming choice of the mouse in that trial, and Δ(*t_i_*) is the cumulative right minus left cue counts up to and including cue *i. v_i_* = *v*(*t_i_*) is the speed of the mouse in the virtual world at the time of the cue, and for the simple linear speed dependencies the standardized version 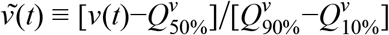 is used, where 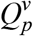 is the *p* probability content quantile of the speed distribution. The shape parameter *k* for the Gamma distribution is furthermore indexed by choice for the C model. The spline-based functions were implemented using an open source C+ + cubic spline fitting library (https://github.com/ttk592/spline/), with modifications (for evaluation convenience) available from https://github.com/sakoay/spline. For a given set of control points {*x*_(*i*)_ = 1,…, *n*), the spline *s*(*x*) is a piecewise continuous and twice continuously differentiable 3^rd^ degree polynomial in between adjacent control points *x_(i)_* and *x_(i+1)_*. Natural boundary conditions were used at the endpoints *x_(1)_* and *x_(n)_*, i.e. the second-order derivatives were defined to be zero at those points so that the solution linearly extrapolates past the range of control points. For the SSA model, the response to the first cue in the session is defined to be μ_*A*_(*t*_1_) =^1^.

Because neural activity can be very different in the rare cases where the mouse halts in the middle of the cue region, only data where the speed v is within 25% of its median value were included in the construction of this model. Point estimates for the model parameters were obtained by minimizing the Gamma-distribution negative log-likelihood (Matlab function *fmincon*):

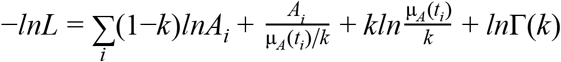

We selected a Gamma distribution hypothesis because the amplitudes were restricted by definition of the impulse response model to take on non-negative values. However because the Gamma distribution is defined only in the positive domain, we had to make an assumption about how to treat data points where *A_i_* = 0. We reasoned that we could substitute these with a noise-like distribution of amplitudes, which were obtained by fitting the impulse response model (Eq. 5) using the same cue timings but simulated noise-only data, which comprised of a *ΔF/F* time-series drawn i.i.d. from a Gaussian distribution with zero mean and standard deviation being *σ^F^*, the estimated fluorescence noise level for that cell. This yielded a set of 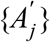 which had some nonzero values from chance fluctuations in the simulated data. All *A_i_* ≤ 10^−3^*σ^F^* were then replaced with 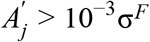 values from this set. This substitution drives the model μ_*A*_ away from zero in the regime of very small response amplitudes, but not beyond noise levels. In order not to run into undefined log-likelihood values during optimization, μ_*A*_ is also bounded below by a small number, μ_*A*_ → *max*(μ_*A*_. 10^−3^σ^*F*^).

Bounds were set on the parameters so that μ_*A*_ ≥ 0 and aspects of the solution remained within a reasonable range. For the angular receptive field hypotheses, the preferred response angle was restricted to *φ*_0_ ∈ [-2π, 2π]. For the skew-Gaussian hypothesis in particular, the width/skew parameters were restricted to σ ∈ [0, 10π] and ζ ∈ [-10π, 10π]. For the sigmoid hypothesis, the restrictions were *ρ_min_* ∈ [0, *max_i_A_i_*], *ρ_max_* ∈ [0, *max_i_A_i_*], γ ∈ [0, 100], and v ∈ [0, 200]. The spline-based functions are defined by their values at the control points, all of which were bounded between [0, *max_i_A_i_*]. The shape parameter of the Gamma distribution was restricted to *k* ∈ [0, ∝). Lastly, the SSA adaptation strength was restricted to γ ∈ [0, 3] and the associated timescale to γ ∈ [0, 7*s*].

The AIC_c_ score for a given model is 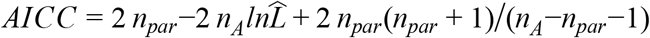 where 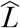 is the optimal model likelihood, *n_A_* the number of response amplitudes i.e. data points, and *n_par_* the number of free parameters for that model. For any one model, the best (lowest AIC_c_) option for the angular receptive field shape *ρ*(φ_*cue*_) is first selected. The relative AIC_c_-based likelihood for two models, then used for model selection as described in the text, is *exp*([*AICC (model 1)-AICC*(*model* 2)]/2).

##### Choice modulation strength

The location-dependent choice modulation strength for cue-locked amplitudes is defined as 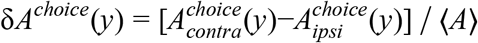 where 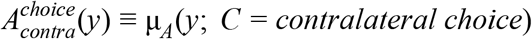 as in Eq. 7, and analogously for ipsilateral choices. This is computed by evaluating the amplitude model prediction vs. location in the cue region, but at fixed φ_*cue*_ corresponding to zero view angle (+ 22° for right-side cues and − 22° for left-side cues) and Δ = 0. The normalization constant is:

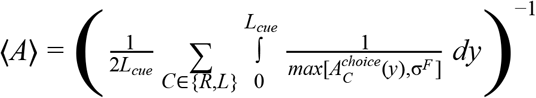

In practice the location-averaging of the model prediction is performed by evaluating the function at 1000 evenly spaced points over the cue region [0, *L_cue_*]. We preferred the harmonic over the arithmetic mean because the activity of cortical cells including cue-locked ones tend to be sparse, leading to a prevalence of small amplitude values that was poorly reflected by the arithmetic mean (which is dominated by high outliers}. To prevent this from being undefined in the rare cases where 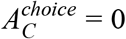, we used the larger of the model prediction vs. the activity noise level σ^*F*^ of that cell.

#### Feedback-loop model

As explained in the main text, the sensory and accumulator states are specified by:

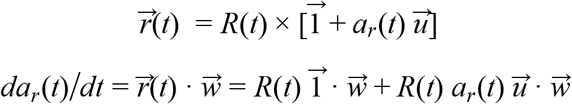

This is an ordinary differential equation that can be solved by the integrating factor method. Writing it in the form 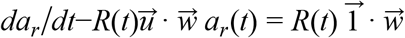, we identify the integrating factor to be 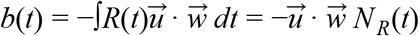, giving a solution:

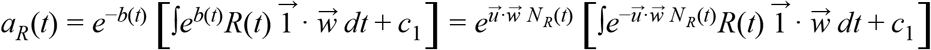

where *c*_1_ is a constant of the integration. We can simplify this:

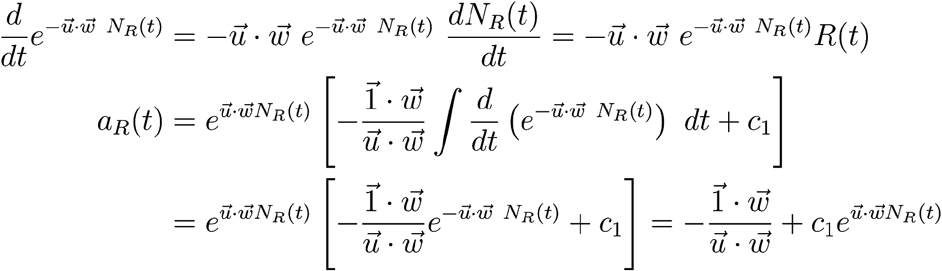

We assume that at the start of the trial, the accumulator has zero content. This requires 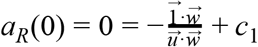, and substituting 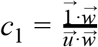 into the above we obtain Eq. 3.

In the case of weak feedback, a Taylor’s series expansion of Eq. 3 gives 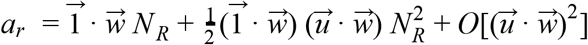. The accumulator thus reduces to perfect integration in the zero feedback limit 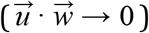, and exhibits growing nonlinearities vs. *N_R_* for stronger feedback. Interestingly the relevant measure of feedback strength is not the “synaptic” weights 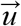, but the projection of the feedback onto the feedforward direction, 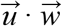. In other words, near-linear integration can be achieved in a neural circuit with strong feedback synapses, so long as there is net cancellation such that 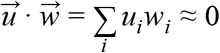. This requires the sensory population state to be at least 2-dimensional.

##### Dependence of cue-response amplitudes on counts

We performed a quasi-independent check that count modulation of cue-locked amplitudes could be detected using a simple linear model in a more controlled experimental situation, as opposed to the multi-factor amplitude modulation models that accounted for angular receptive field effects, running speed, and so forth (Fig. 5). This was achieved by restricting the data to the last third of the cue period, which minimizes behavioral differences like running speed, place dependence, etc.

To control for categorical choice effects, we fit a different linear model 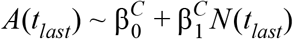 for each subset of trials with choice *C* ∈ {*right, left*}, where *A*(*t_last_*) is the amplitude of the response to the last cue in the last third of the cue period, and *N*(*t_last_*) is the total number of preferred-side cues in that trial. To control for the visual angle, the same type of weighted linear fit as used to assess Δ-modulation of non-cue-locked cells was used, with weights chosen so that the φ_*cue*_ distributions are the same in three equally sized quantile bins when conditioned on *N* E {1 −3, 4 −6, 7-9, > 10}. Null hypotheses were constructed by shuffling the amplitudes *A* across trials for a fixed choice *C*, and the slope 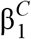 for a cell is considered to be significant if less than 5% of shuffled-data fits have slopes greater than that value. For cells with either significant 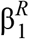 or 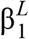, these slopes are highly significantly correlated across choice categories *[r =* 0.44, *p* = 2.1 × 10^−5^), and we therefore used the choice-averaged slope 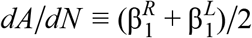 to identify significantly modulated cells. The significance of this choice-averaged slope was analogously defined by comparing its value to the choice-averaged slopes in shuffled-data fits.

Because sensory responses are expected to have a causal influence on the behavioral choice, it is possible that the above method to control for choice effects may artificially introduce a negative correlation between the amplitudes of sensory responses *A* and cue counts *N*. For example, trials with low right-cue counts tend to be more ambiguous (smaller differences between right- and left-cue-counts), and therefore positive fluctuations in right-cue sensory responses may bias the decision towards a right-turn choice. This is in contrast to trials with high right-cue counts, which tend to already have unambiguously high rightwards evidence, and therefore both positive and negative fluctuations have little effect on the decision to turn right. In other words, when selecting only right-choice trials we may inadvertently also select trials where the sensory response amplitudes are biased upwards for low *N* and less biased for high *N*, resulting in a negative *dA/dN*. We checked for this by performing linear regression of *A* vs. *N* without controlling for choice (i.e. using all of the data), and comparing this to when choice was controlled for. The slopes *dA/dN* estimated using both methods are highly correlated (Fig. S5F; *r* = 0.63, *p* = 2.5 × 10^−42^ for all cells; *r* = 0.8, *p* = 7.1 × 10^−15^ for cells identified as having a nonzero slope), with no indication that the choice-control method biases the computed *dA/dN* towards lower values due to the aforementioned negative correlation (Fig. S5G, all points lie close to the diagonal line). We therefore conclude that any choice-selection biases are negligible in our analysis.

For the non-parametric visualization of the dependence of *A* on *N* (Fig. 6D), we computed the weighted mean amplitude in the same bins of *N* as above and with the same weights as used to equalize the φ_*cue*_ distributions. To account for the width of the bins vs. non-uniform distributions of the generated cue counts, the abscissa of this plot is shown at the bin-average *N*. Similarly, the Fano factor *F(A)* ≡ *var(A)/mecm(A)* was computed per *N*-bin as the weighted variance over weighted mean. Linear regression was then performed for *F(A)* vs. the bin-average *N*.

##### Predicted Fano factor of cue-locked responses

Here we consider the theoretical consequences of *N_R_* in Eq. 3 not being perfectly deterministic, but instead Gaussian-distributed random variate with mean *N_R_* and variance 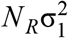 (as per the central limit theorem as explained in the text). This means that 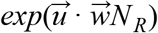 is lognormally distributed with non-logarithmized mean 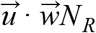 and variance 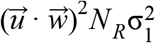. Letting 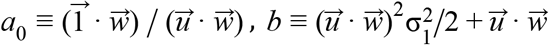 and 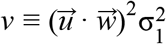 to simplify notation, the accumulator state has the following mean and variance (Johnson, Kotz, and Balakrishnan 1994):

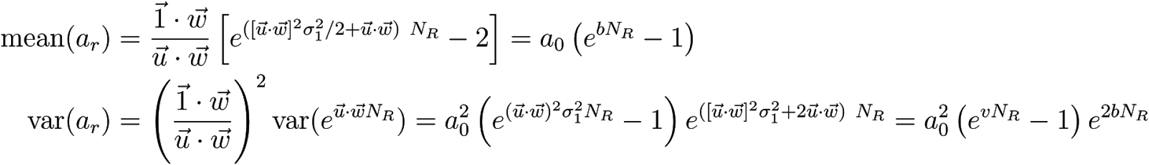

We want to understand the trend of the Fano factor vs. *N_R_* for the activity of a single sensory neuron under pulsatile input. From Eq. 1 this is *r_t_* = *R* (1 + *a_r_u_i_*) where *a_r_* is the stochastic accumulator state above and *R* corresponding to a single input pulse is a Gaussian random variate with mean 1 and variance 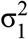. For simplicity we will assume that *R* and *a_r_* are independent random variables, which would be the case in the discrete-time version of Eq. 3 since for causality reasons the scaling by the accumulator state should depend on the inputs up to but not including *R.* We use the following rule for the variance of a product of two independent random variables:

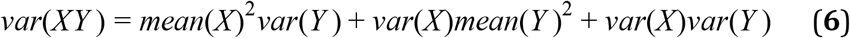

The Fano factor of the distribution of *r_i_* is thus:

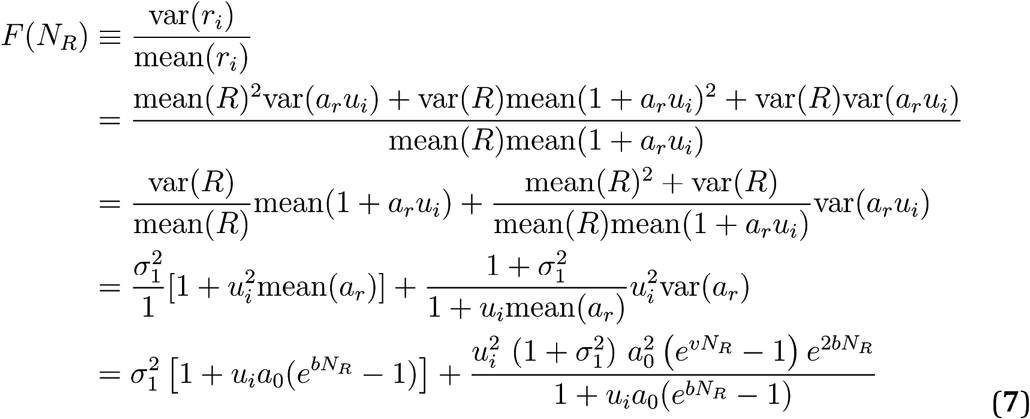

In the regime of small *N_R_* we expand Eq. 10 in a Taylor’s series, 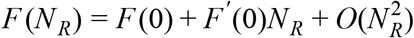. Direct substitution of *N_R_* = 0 in the above gives us 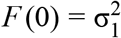. For the first-order term, we have:

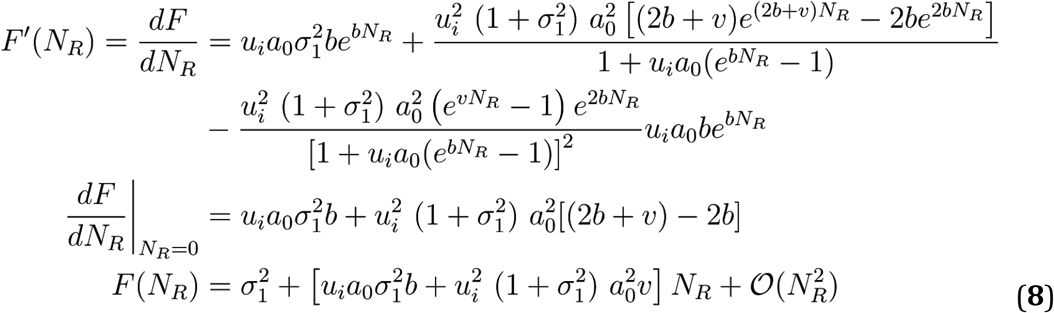

All the variables *a*_0_, *b*, *v*, and are positive, whereas *u_i_* can take on either sign. This means that for cells with feedback strength *u_i_* > 0, all the terms in Eq. 8 are non-negative, in particular the first-order term a.k.a. the slope, and *F* is thus a monotonically increasing function of *N_R_*. Viewed as a function of the feedback strength, *f(u_i_)* ≡ *F* (0) has roots at 0 and 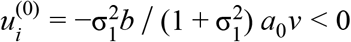, so for *u_i_* < 0 cells there is a range of 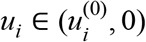 where *F*′(0) < 0 i.e. *F* is a monotonically decreasing function of *N_R_*, for small enough *N_R_* such that the Eq. 8 expansion holds. However as *N_R_* grows large but still for *u_i_* < 0, this trend reverses. We can see this by considering the behavior of *F*(*N_R_*) close to the singular point where *mean*(1 + *a_r_u_i_*) → 0. Going back to the form of Eq. 7 as 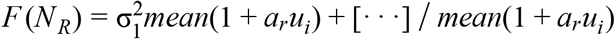, the first term proportional to the mean becomes negligible compared to the second which is inversely proportional. The latter diverges to + ∞, because the denominator decreases monotonically from 1 down to 0, while the numerator is always non-negative. The latter follows from the squares of real numbers always being non-negative, which is also the case for those of the form *e^x^* ≥ 1 when all *x* ≥ 0. Thus to recap, for *u_i_* < 0 cells the Fano factor first decreases with *N_R_*, then at some point turns to increasing since it diverges to + ∞ as *mean*(1 + *a_r_u_i_*) → 0. For extremely strong negative feedback such that the mean sensory response would decrease below zero, we must extend the model to include rectification effects to prevent this from happening. This is beyond the scope of this work, although depending on the type of rectification, this can also modify the behavior of the Fano factor (Charles et al. 2018).

We illustrate the above analytical results using simulations where for a given true input count of *N*, we model the single-pulse stochasticity *R* as a Gamma-distributed random variable with mean 1 and variance, the accumulator state as Eq. 3 but with *N_R_* replaced with a Gamma-distributed random variable with mean *N*-1 and variance 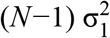 (discrete-time version). The parameters used are 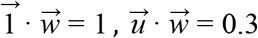, and *u_i_* =± 0.01. The resulting distribution of sensory responses as in Eq. 1 is shown in Fig. S5A, but bounded to be no less than zero. For *N* = 1, the accumulator state is zero and thus there is no distinction between positively and negatively modulated sensory responses. For *N* = 5 and *N* = 10 the feedback effect grows progressively larger, inducing a separation between *u_i_* > 0 and *u_i_* < 0 cases. Fig. S5B shows the mean, variance, and Fano factor of the sensory response distributions for a range of true counts. The Fano factor monotonically increases for *u_i_* > 0 and has the expected non-monotonic trend of first decreasing then increasing for the *u_i_* < 0 case given our choice of parameters. These results are not sensitive to the type of distribution (e.g. Gamma vs. Gaussian) modeled, so long as the variance remains finite and there is not an excessive amount of probability mass that is truncated at zero.

##### Sources of noise in various accumulator architectures

To account for imperfect psychophysical performance, we consider three sources of noise in modeling the distribution of *a_r_*.

First is sensory noise, any source of stochasticity that occurs either in the stimulus *R(t)* itself or in the activity 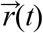 of the sensory neurons. Because the latter are slaved to their input drive, the detail of where along the feedforward path this noise arises from does not matter in its effect on the accumulator state. Also since only the projection of the sensory state onto the readout direction 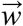 matters, only the net 1-dimensional variability along 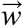 matters. Sensory noise is thus equivalent to replacing *R(t)* → *R(t)* + *ε(t)* where *ε(t)* is a (scalar) noise process. Assuming the central limit theorem, we model this as replacing the true integral 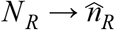 where 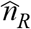 is a Gaussian distributed random variable with mean *N_R_* and spread 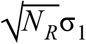.

Secondly, we consider modulatory noise that may be present in gain modulations of the sensory population. For the *fdbk* model, as the sensory gain modulation is proportional to the feedback weight, this is equivalent to there being stochasticity in the feedback weights 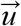, which for tractability we consider to be slow on the timescale of the cue period. Again, only the 1-dimensional variability along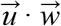 matters, so in sum this source of stochasticity leaves Eq. 3 unchanged but instead can be considered as 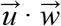 being drawn randomly per trial, 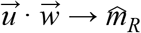. We model 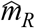 with a lognormal distribution, which is the limiting case of a product of many positive random variables. For *the ffwd* model, we instead hypothesized a non-specific source of slow gain fluctuations, equivalent to scaling the accumulator value with a stochastic variable per trial, 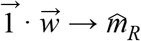 in Eq. 4.

Third, for both models there can be noise associated with comparing the two accumulators to form a decision, which we model as the decision variable *c* being Gaussian-distributed around *a_r_*–*a_i_*.

Sources of variability that we for simplicity do not model include initial and drift noise in the accumulator, as these were seen to be negligible according to behavioral models of pulsatile evidence accumulation (Brunton, Botvinick, and Brody 2013; Scott et al. 2015; Lucas Pinto et al. 2018). Accumulator drift noise also predicts a time-dependence to the behavior that we do not observe.

Formally, the modified differential equation for the evolution of the *fdbk* model accumulator state given a sensory noise process *ε(t)* and a modulatory noise process *(t)* is:

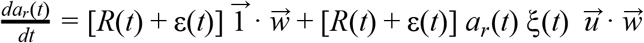

The solution as discussed above (assuming that *ξ(t)* is slower than the trial duration) is:

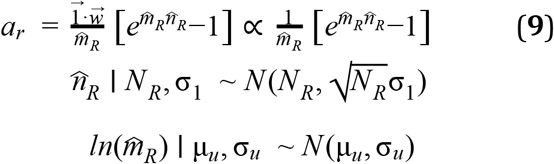

The model predicts that the animal should make a choice to the right with probability given by the decision variable *c* ~ *N*(*a_r_−a_l_*,σ_*c*_) being positive. The likelihood of getting exactly (*k_R_,k_L_*) right- and left-choice trials in the behavioral data for a fixed (*N_R_,N_L_*) is binomially distributed around this probability. The constant 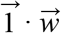 that *a_r_* is proportional to can be neglected since this just sets the arbitrary units of the accumulator, which we presume that the brain adjusts so that the right and left accumulators can be compared in a way that is not biased by having potentially different units. We additionally include a lapse rate that acts as the animal instead making a fair coin toss to go right *p_lapse_* of the time. Lastly, we maximized the model likelihood over the five free parameters {σ_1_, μ_*u*_, σ_*u*_, σ_*c*_,*p_lapse_*} best fit the behavioral data as described in the next section.

As discussed in the text, we assessed how well the the *fdbk* model predicts behavior in comparison to three other architectures. The *ffwd* model has accumulator states distributed as 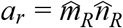, and the full set of five free parameters {σ_1_, μ_*u*_, σ_*c*_,*p_lapse_*} same as for the *fdbk* model. The *intg* variant of this model has no modulatory nor accumulator-comparison noise 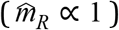, and so has two free parameters {σ_1_,*p_lapse_*}. The *webr* variant instead has no sensory nor accumulator-comparison noise 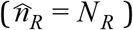, and so has three free parameters {μ_*u*_, σ_*u*_, *p_lapse_*} The modulatory noise 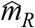 is the cause of Weber-Fechner scaling in all three of the *webr, fdbk* and *ffwd* models.

##### Fitting accumulator models to behavioral data

The behavioral data that we used here are:

- mice with ≥ 500 trials from (Lucas Pinto et al. 2018)
- data from head-fixed rats performing a flash accumulation task (Scott et al. 2015)

Let *k_R_*(*N_R_,N_L_*) and *k_L_*(*N_R_,N_L_*) be the number of right- and left-choice trials respectively out of all trials in the behavioral data with a given number of true right (*N_R_*) and left (*N_L_*) cue counts. A particular accumulator model *M* ∈ {*intg, webr, ffwd, fdbk*} is characterized by its predicted probability of right-choice trials given the true cue counts, 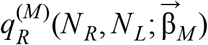 where 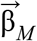 are the free parameters of the model. Including a lapse rate *p_lapse_*, we optimized the models by minimizing their negative log-likelihood, which have the same form as given by the binomial probability distribution:

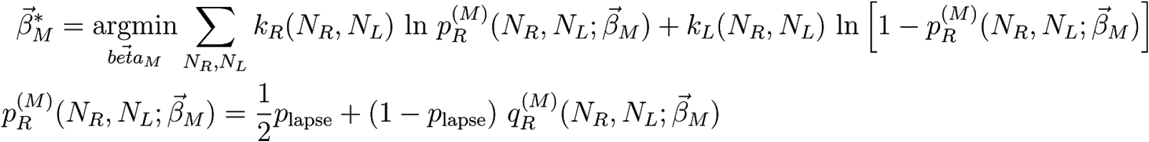

For the *intg* model, the probability distribution of right- and left-accumulator states are Gaussian according to the central limit theorem. Assuming that the accumulators are independent and have the same parameters, the joint probability distribution is 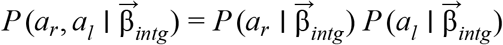, and the right-choice probability is given by the region 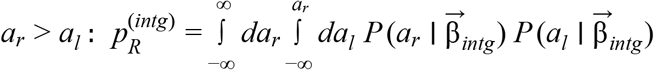. This has a well-known simplification in that the difference *c* ≡ *a_r_*−*a_l_* of two (independent) Gaussian-distributed variables is also Gaussian-distributed, so 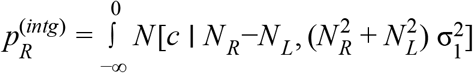. For the non-negative variant *intg*^+^ (Fig. S5I) we assume Gamma-distributed sensory noise, and performed the following integral numerically (Matlab functions *integral, gampdf* and *gamcdf*):

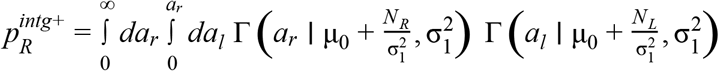

μ_0_ is an additional free parameter of the *intg*^+^ model, which was required because the Gamma distribution is otherwise not defined at zero counts. Otherwise the Gamma distributions were selected to have the same mean and variance as a function of the true counts as the Gaussian distributions in the intg model, following the central limit theorem hypothesis.

For the remaining three models, we evaluated the necessary integrals using a Monte Carlo method. For a given combination of (*N_R_,N_L_*), we drew 200,000 samples each for the five stochastic quantities 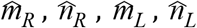(see Eq. 9), and 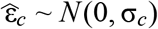 which acts as the accumulator comparison noise. For the non-negative variants *webr*^+^, *ffwd*^+^, and *fdbk*^+^, the integrated distributions are instead 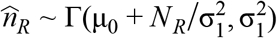 and 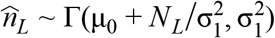. The decision variable was computed as 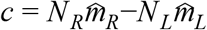 for the *webr* models, 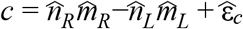 for the *ffwd* models, and 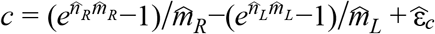 for the *fdbk* models. The right-choice probabilities for the model were then estimated as the fraction out of the 200,000 samples for which *c* > 0.

The method used for minimizing the negative log-likelihood for the *intg* models was the interior-point algorithm implemented by the Matlab function *fmincon*. For the other models, this could not be used because the stochastic nature of the Monte Carlo integration resulted in slightly non-smooth log-likelihood landscapes. Instead, the gradient-free direct search method *patternsearch* (Global Optimization Toolbox) was used for these models. Parameter ranges were bounded during the minimization in order to obey non-negativity constraints and in some cases to avoid numerical under/overflow issues at extreme values (particularly for the *fdbk* model, which involves exponentiation). For all models, σ_1_ ∈ [0,100], μ_*u*_ ∈ [-5,2], σ_*u*_ ∈ [0,20], and *p_lapse_* ∈ [0,0.5]. The bounds for the comparison noise was σ_*c*_ ∈ [0,20] for the *ffwd* models and σ_*c*_ ∈ [0,50] for the *fdbk* model. The motivation for this is because the accumulator states in the *fdbk* model are on an exponential scale compared to the *ffwd* models, thus larger values of comparison noise for the *fdbk* models can have comparable effects on *p_R_* compared to smaller values in the ffwd models. Lastly, for the non-negative model variants we constrained μ_0_ ∈ [0.01,10], and we optimized 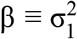 directly instead of σ_1_, with bounds β ∈ [0.2,20] since the Gamma distributions become undefined (and run into numerical integration issues) as β → 0.

Interestingly, for the mouse *fdbk* model fits, accumulator-comparison noise had a more dominant effect than sensory noise (Fig. S5H-left, Fig. S6A), whereas the reverse holds for rats (Fig. S5H-right, Fig. S6B). This could be related to the stimuli in the rat task being very brief (10ms) LED flashes, compared to the high visual salience of cues in the mouse task. The navigational component of the mouse task instead adds unmodeled sources of behavioral variability that may have been absorbed into the accumulator-comparison noise term.

##### Theoretical effects of various sources of noise

In the *ffwd* model, the relative uncertainty in 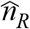 diminishes with increasing *N_R_* as for the intg model, so asymptotically 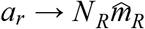 as in Weber-Fechner scaling. More formally, we can perform a variance analysis to assess the contributions of sensory, modulatory, and accumulator-comparison noise. The variance of this is given by application of Eq. 6:

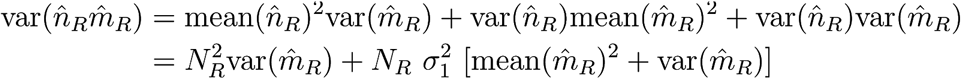

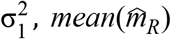, and 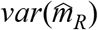 remain at fixed magnitudes as we consider growing *N_R_*, so eventually the 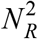 term dominates, leading to Weber-Fechner scaling. The decision variable is 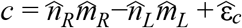 as previously discussed, from which we obtain 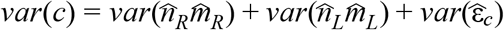 which follows from the two accumulators and the comparison noise all being uncorrelated random variates. The comparison noise variability 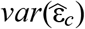 is the only non-count-dependent term in this expression, and given the non-negative nature of all other terms, therefore has the most quickly diminishing effect with increasing counts.

For the *fdbk* model, even though Eq. 3 predicts that the accumulator state is exponentially related to the true counts, this does not on its own predict any change in perceptual performance if there is only sensory noise. This is because if 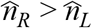 for the perfect integrator, then 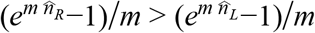, being a monotonic transformation of both left- and right-hand-sides (assuming *m* > 0). What distinguishes the *ffwd* and *fdbk* models is therefore the interaction of sensory and other (downstream) sources of noise. At small counts we can expand Eq. 3 as 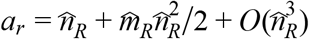, which means that at small counts i.e. to first order the predicted performance follows the *intg* model more closely than the *ffwd* model, seen in Fig. 7B as a steeper increase in performance (for everything else fixed). At high counts 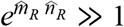, so 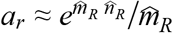. Making use of the fact that *ln a_r_* > *ln a_l_* if and only if *a_r_* > *a_l_*, we can equivalently consider the performance of comparing the log-transform of the accumulators, 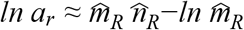. At large counts the 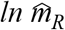 term becomes negligible (and already 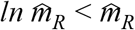), so also the *fdbk* model converges to the *ffwd* model, and in the same way, Weber-Fechner scaling.

## Supplemental Information

**Table SI.** Number of imaging sessions and mice for various areas and layers, for the main experiment.

|  | Layers 2/3 |  |  |  |  |  | Layer 5 |  |  |  |  |  |
| --- | --- | --- | --- | --- | --- | --- | --- | --- | --- | --- | --- | --- |
|  | V1 | AM | PM | MMA | MMP | RSC | V1 | AM | PM | MMA | MMP | RSC |
| # sessions | 9 | 18 | 11 | 15 | 8 | 30 | 8 | 12 | 6 | 9 | 7 | 12 |
| # mice imaged | 4 | 8 | 6 | 9 | 4 | 8 | 4 | 7 | 3 | 8 | 4 | 6 |

**Table S2.** Overall performance and number of imaging sessions for the main experiment, per mouse (rows), in various areas and layers (columns). Mice of the Thy1 GP5.3 strain have names starting with “gp”, and those from the Ai93-Emx1 strain have names starting with “ai” (see Methods).

|  |  | Layers 2/3 |  |  |  |  |  | Layer 5 |  |  |  |  |  |
| --- | --- | --- | --- | --- | --- | --- | --- | --- | --- | --- | --- | --- | --- |
|  | % correct | V1 | AM | PM | MMA | MMP | RSC | V1 | AM | PM | MMA | MMP | RSC |
| gp31 | 73.0 | 1 | 1 | 1 | 1 | 0 | 2 | 1 | 1 | 1 | 1 | 1 | 1 |
| gp36 | 75.3 | 2 | 1 | 1 | 1 | 1 | 2 | 2 | 1 | 1 | 1 | 1 | 1 |
| gp37 | 70.1 | 0 | 1 | 0 | 1 | 1 | 1 | 0 | 0 | 0 | 1 | 0 | 0 |
| gp40 | 67.7 | 0 | 2 | 0 | 0 | 0 | 9 | 0 | 1 | 0 | 0 | 0 | 5 |
| gp42 | 67.1 | 0 | 1 | 0 | 2 | 3 | 4 | 0 | 1 | 0 | 1 | 2 | 1 |
| gp46 | 67.8 | 0 | 0 | 1 | 3 | 0 | 4 | 0 | 0 | 0 | 1 | 0 | 2 |
| ai50 | 68.7 | 0 | 2 | 1 | 3 | 0 | 7 | 0 | 1 | 0 | 2 | 0 | 2 |
| ai53 | 67.8 | 2 | 0 | 0 | 1 | 0 | 0 | 2 | 0 | 0 | 0 | 0 | 0 |
| ai55 | 71.6 | 0 | 3 | 0 | 2 | 0 | 1 | 0 | 3 | 0 | 1 | 0 | 0 |
| ai56 | 71.9 | 4 | 7 | 6 | 1 | 3 | 0 | 3 | 4 | 4 | 1 | 3 | 0 |
| ai57 | 67.2 | 0 | 0 | 1 | 0 | 0 | 0 | 0 | 0 | 0 | 0 | 0 | 0 |

**Table S3.** Factors (columns) present in the six amplitude-modulation models (rows). φ_CMe_ and speed are effects that depend on the momentary visual stimulus and are included in most models. Sensory adaptation, location, and cue counts (*#R, #L*) are factors that can explain systematic trends vs. time, and are exclusively present in any one model.

|  |  | Factors |  |  |  |  |  |
| --- | --- | --- | --- | --- | --- | --- | --- |
| | | $\varphi_{cue}$ | Speed | Adaptation | Location | Choice | #R,#L |
| Models | $\varphi$ | ✓ | | | | | |
|  | v | ✓ | ✓ |  |  |  |  |
|  | SSA | ✓ | ✓ | ✓ |  |  |  |
|  | y | ✓ | ✓ |  | ✓ |  |  |
|  | C | ✓ | ✓ |  | ✓ | ✓ |  |
| | $\Delta$ | ✓ | ✓ | | | | ✓ |

**Figure S1.**
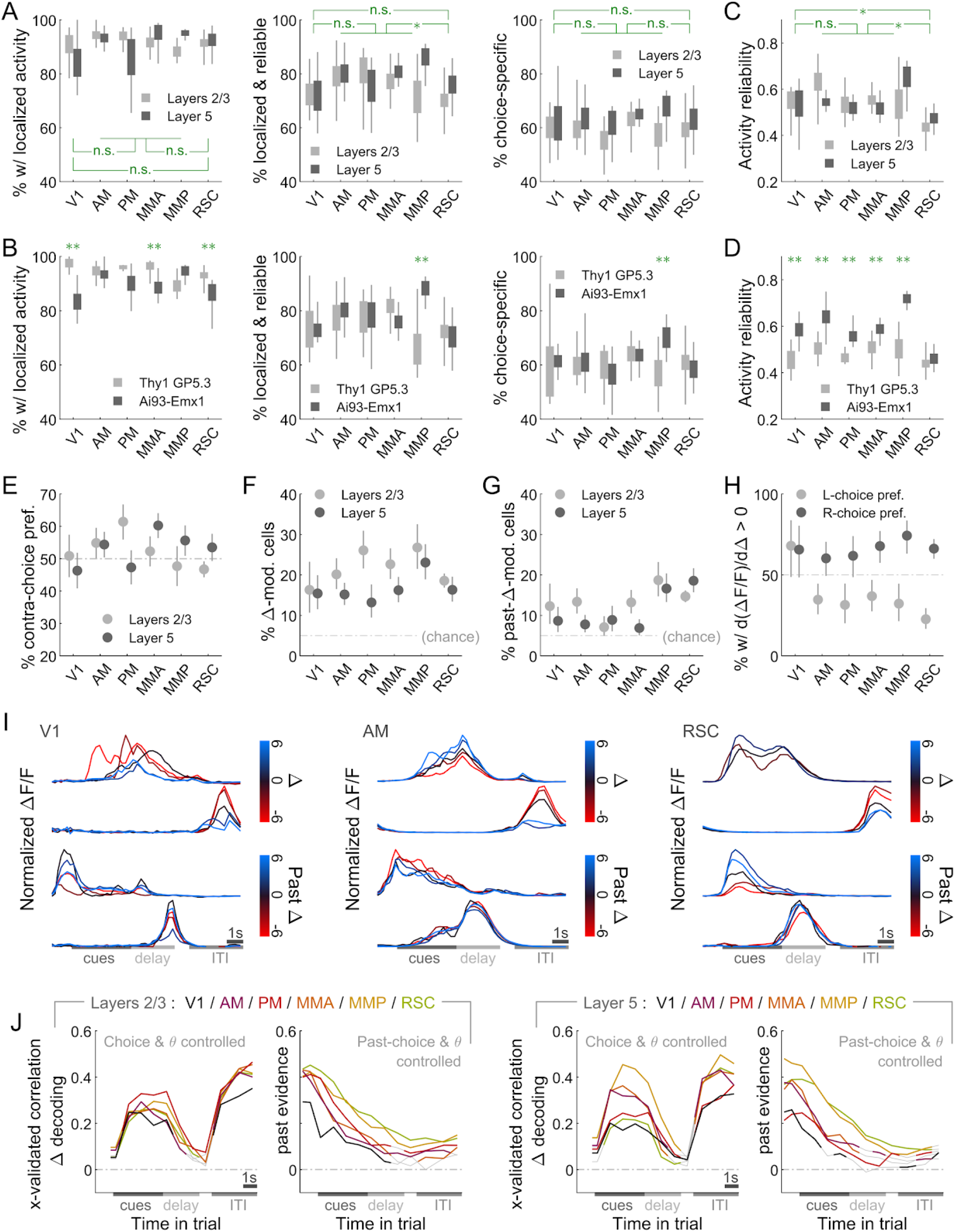
Additional statistics for choice-specific sequences, and evidence modulation. **(A)** Percents of cells that have significant ridge-to-background excess vs. a permutation test (left plot), and additionally are active within their firing fields in ≥ 20% of their (preferred-choice, if any) trials (activity reliability index, middle plot), and additionally have different activity levels in right-vs. left-choice trials (right plot). Error bars: std. dev. across imaging sessions. Rectangles: Median and S.E.M. Stars: significant differences in means (Wilcoxon rank-sum test). **(B)** Like **(A),** but comparing two strains of mice. Data were pooled across layers. Double-stars indicate areas for which there was a significant difference in means (Wilcoxon rank-sum test). **(C)** Average reliability of choice-specific cells in a given area/layer, defined as the fraction of trials in which the cell is significantly active within its putative firing field. Error bars as in **(A). (D)** Like **(C),** but comparing two strains of mice. Data were pooled across layers. Double-stars indicate areas for which there was a significant difference in means (Wilcoxon rank-sum test). **(E)** Percents of choice-specific cells with higher activity in contralateral-than ipsilateral-choice trials, for various areas/layers. Error bars: 95% C.I. **(F)** Percent of cells that have activity significantly modulated by Δ = ***#R-#L,*** after controlling for view angle, choice, and reward outcome (see Methods). Data were pooled over recordings for a given area/layer. Error bars: 95% C.I. **(G)** Same as **(F)** but for cells that are significantly modulated by the evidence in the past trial. **(H)** Percents out of all significantly Δ-modulated cells that have positive Δ-modulation slopes, shown separately for left-vs. right-choice preferring cells. Data were pooled across layers for a given area. Error bars: 95% C.I. **(I)** Activity vs. time in the trial for 12 non-cue-locked cells with significant Δ-modulation. The bottom two rows are for cells with significant dependence on Δ in the previous trial. Lines: average activity across trials with similar Δ values (color). **(J)** Correlation between the value of Δ and that decoded from neural states in various areas/layers, analogous to Fig. 3A-B. To control for choice, decoding is performed separately in right-vs. left-choice trials, and the results averaged.

**Figure S2.**
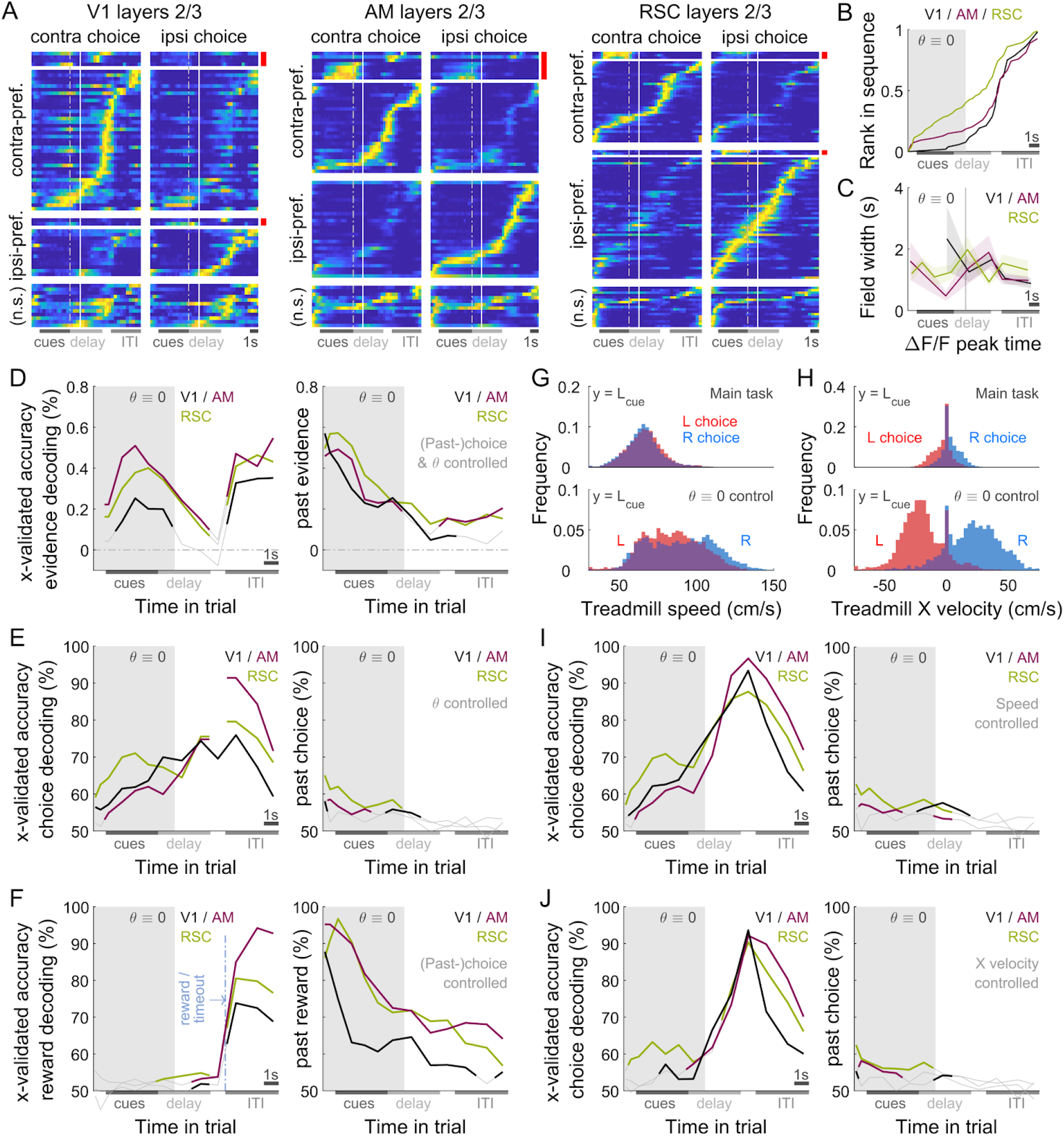
Qualitatively similar neural phenomena in view-angle-locked control experiments. **(A-C)** Choice-specific sequences and tiling statistics, as in Fig. 2A-C. **(D-F)** Evidence, choice and reward decoding accuracies, as in Fig. 3. **(G)** Distribution of treadmill rotation speeds at the end of the cue period, for data collected in the main task (top plot) vs. θ-controlled scenario (bottom plot). **(H)** As in **(G),** but for the treadmill X (roll) velocity. (H) Choice decoding accuracy as in **(E),** but controlling for the treadmill speed only, in lieu of view angle. (I) Choice decoding accuracy as in **(E),** but controlling for the treadmill X velocity in lieu of view angle.

**Figure S3.**
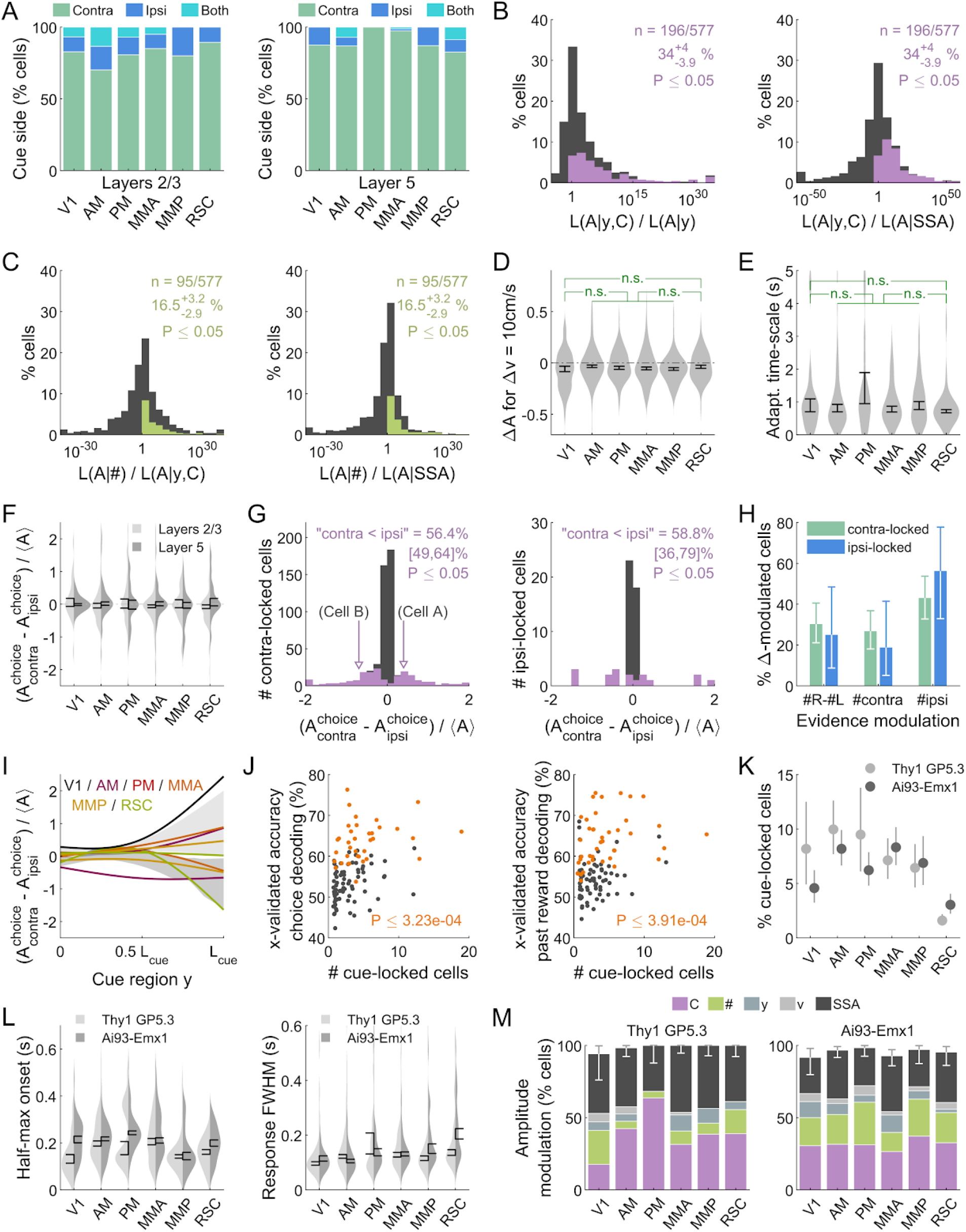
Additional statistics for cue-locked cells and amplitude modulations thereof. **(A)** Percentages of cells with various types of contextual modulations, as determined by the selected best amplitude model per cell. Error bars: 95% C.I. for the sum over modulation types; the remaining fraction are cells that favor the angular-receptive-field-only model. **(B)** Distribution of AIC_c_ likelihood ratios for the location-and-choice model vs. location-only model (left plot), and vs. the SSA model (right plot). The colored area corresponds to cells for which the location-and-choice model is the best for that cell. Data were pooled across all recordings. **(C)** Same as **(B)** but for the cue-counts model vs. the location-and-choice model (left plot), and vs. the SSA model (right plot). **(D)** Distribution (kernel density estimate) of predicted speed-induced changes in amplitudes for a change in speed of 10cm/s. Data were pooled across layers and include cells that favor the SSA, y, C, and # models. Error bars: S.E.M. Stars: significant differences in means (Wilcoxon rank-sum test). **(E)** Distribution of adaptation/enhancement timescales for cells that favor the SSA model, defined as the amount of time taken for the amplitude to recover by a factor of 1/*e*. Error bars: S.E.M. Stars: significant differences in means (Wilcoxon rank-sum test). **(F)** Distribution of choice modulation effect sizes for cue-locked cells in various areas/layers. Cells with numerically near-zero modulations (*|δA^choice^*| ≤ 10 ^4^) were excluded. Error bars: S.E.M. **(G)** Distribution of choice modulation effect sizes for contralateral-cue-locked (left plot) and ipsilateral-cue-locked (right plot) subsets of cells. The colored area corresponds to cells that favor the location-and-choice model. Arrows indicate the effect size values for cells in Fig. 5A-B. Data were pooled over all areas. The left- (right-)most bins include under- (over-)flow. **(H)** Proportions out of all significantly count-modulated cells for which the best evidence-based model is that which depends on the difference (left columns) in or single-side numbers (middle and right columns) of cues, separately for contralateral- and ipsilateral-cue-locked cells. Error bars: 95% C.I. **(I)** Choice modulation strength vs. location in the cue period for ipsilateral-cue-locked cells. **(J)** Choice (left plot) and reward (right plot) decoding accuracy vs. number of cue-locked cells decoded from. The highlighted (orange) points are for sessions where the decoding accuracy is significantly above chance. **(K)** Percent of cue-locked cells vs. area, like Fig. 4H except comparing two strains of mice. Data were pooled across layers. **(L)** Onset of half-max response (left plot) and full-width-at-half-max (right plot) of cue-locked cell responses, like Fig. 4I-J except comparing two strains of mice. Data were pooled across layers. **(M)** Percentages of cue-locked cells that favor various amplitude modulation models, like Fig. 5C except comparing two strains of mice. Data were pooled across layers.

**Figure S4.**
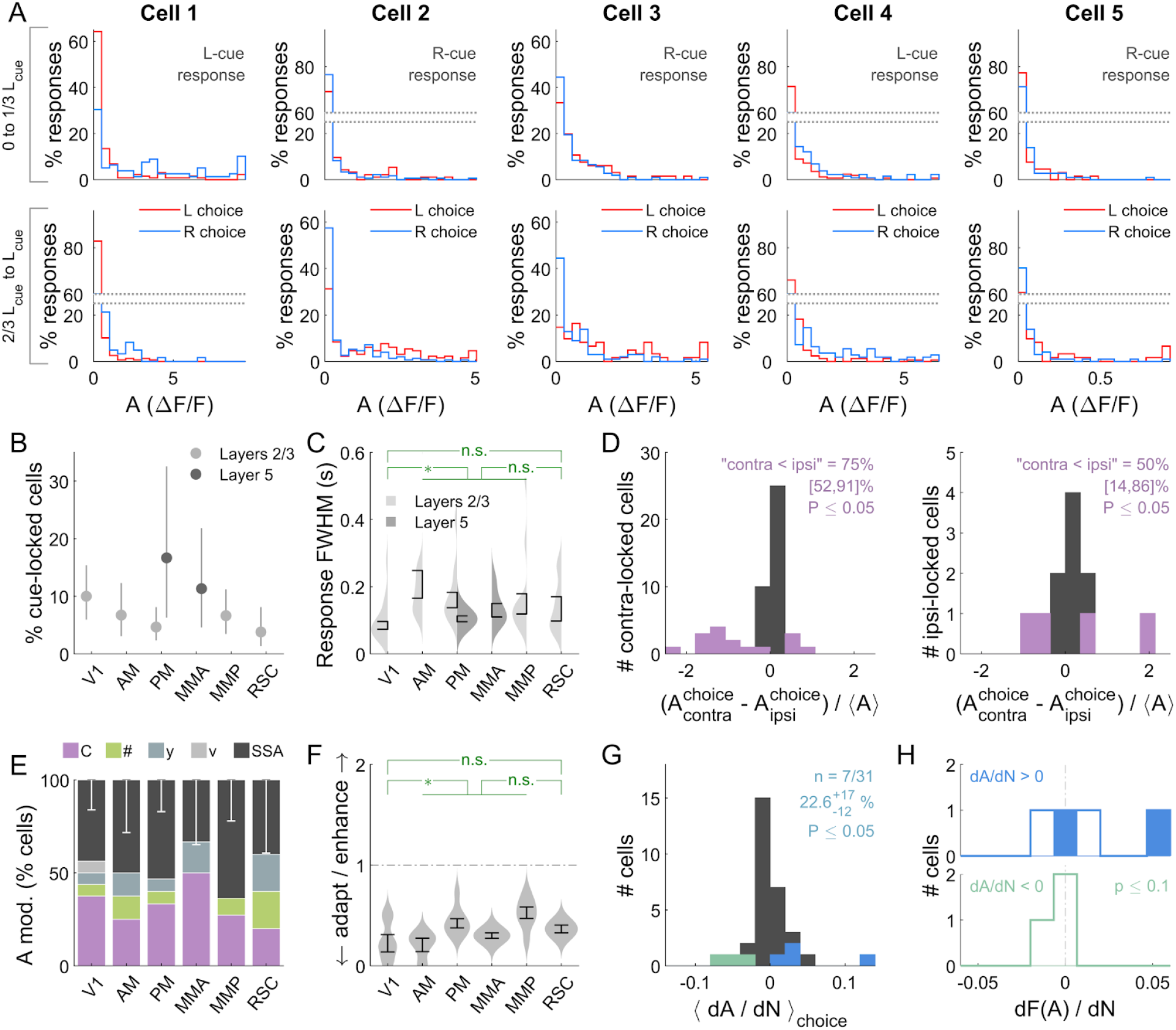
Qualitatively similar cue-locked amplitude modulations in view-angle-locked control experiments. **(A)** Distribution of amplitudes in right vs. left choice trials in the view-angle-locked control experiments, for ten cue-locked cells with the highest choice modulation indices. **(B)** Percents of cue-locked cells for various areas/layers, as in Fig. 4H. **(C)** Distribution (kernel density estimate) of full-width-at-half-max of the cue-locked responses, as in Fig. 4J. **(D)** Distribution of choice modulation strengths for contralateral- and ipsilateral-cue-locked cells, as in Fig. S3G. **(E)** Percentages of cue-locked cells that favor various contextual-modulation models, as in Fig. 5C. **(F)** Adaptation/enhancement factors for cells that favor the SSA model, as in Fig. 5D. **(G)** Distribution of choice-averaged amplitude-vs.-counts slope, with significant cells highlighted in color, as in Fig. 6C. **(H)** Distribution of Fano-factor-vs.-counts slopes for significantly positively and negatively modulated cells in **(G)**. c.f. Fig. 6E.

**Figure S5.**
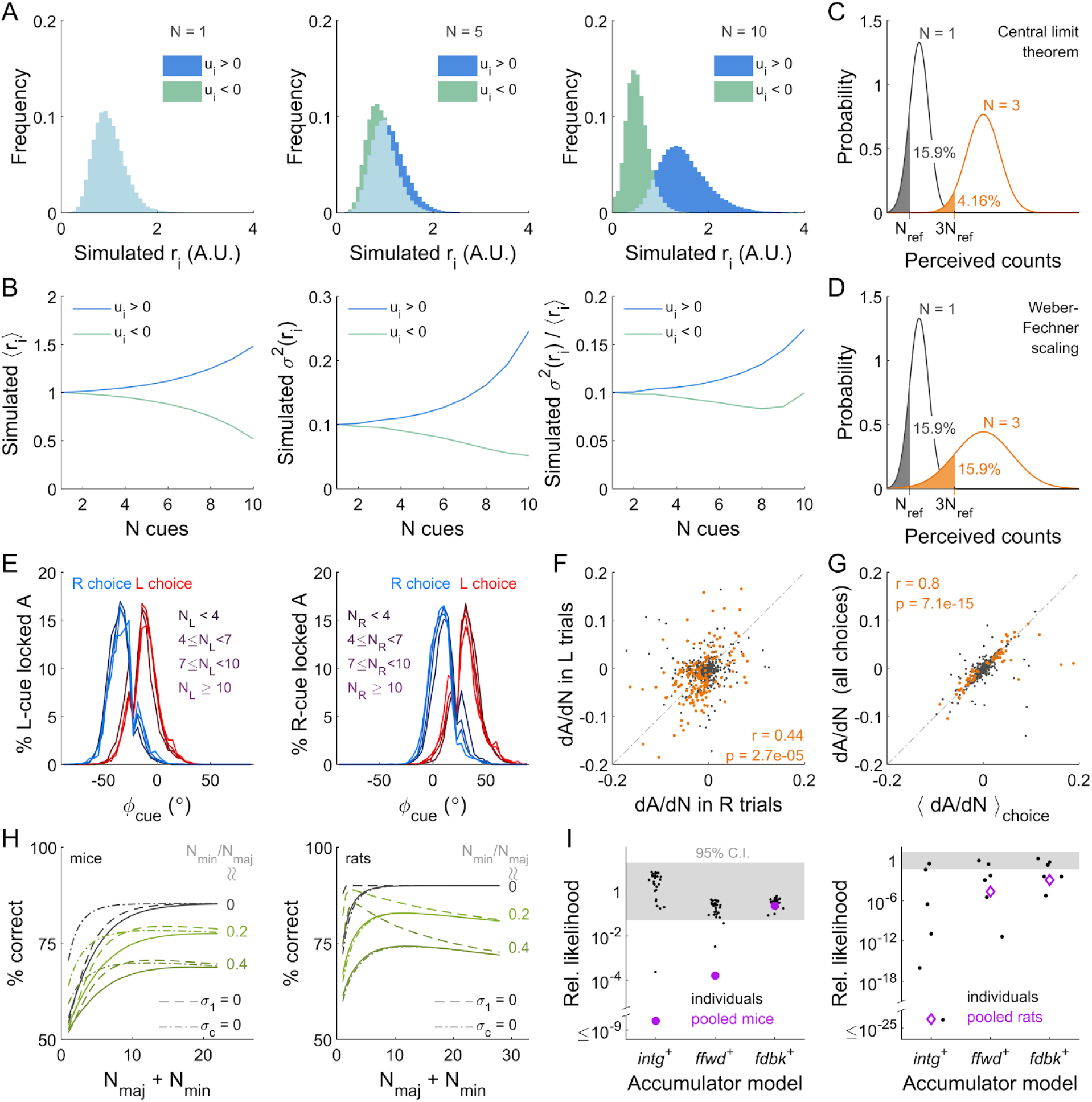
Neural phenomena and behavioral performance predictions compatible with the fdbk circuit model. **(A)** Distribution of simulated sensory responses according to the feedback-loop model Eq. 3, for a range of true counts (columns) that result in a stochastic distribution of accumulator states. Positively (negatively) modulated neurons are shown in blue (green). **(B)** Mean (left), variance (middle), and Fano factor (right plot) for the sensory responses in **(A). (C)** Illustration of how the distributions of perceived counts should change given a true count of *N* = 1 (gray) vs. a true count of *N* = 3 (orange), as predicted by the Weber-Fechner Law of perceptual discrimination. This law prescribes that the *N* = 3 distribution is equivalent to taking the *N* = 1 distribution and scaling the perceived-counts axis by a factor of 3, including the reference level *N_ref_* → 3*N_ref_*. This preserves the area under the reference level (15.9%) and for the same reason perceptual discriminability. **(D)** As in **(C)**, but as predicted by the central limit theorem instead. Given 3 times more integrated counts, the width of the *N* = 3 distribution increases by 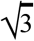 relative to that of the *N* = 1 distribution. This leads to less area falling under the reference level (15.9% for *N_ref_* as opposed to 4.16% for 3*N_ref_*), and corresponds to improved perceptual discriminability at higher counts. (E) Dependence of visual angle distributions on the number of left (left plot) and right (right plot) cues, for either choice category (color). (F) Slope of linear regression of cue-locked amplitudes vs. cue counts, computed either right-choice (x-axis) or left-choice (y-axis) trials only. Highlighted points are cells for which either of these slopes are significantly different from zero (permutation test; see Methods). (G) Slope of linear regression of cue-locked amplitudes vs. cue counts, computed either with controlling for choice (x-axis) or using all the data regardless of choice (y-axis). Highlighted points are for cells with a significant *dA/dN* slope post controlling for choice. More cells could be identified as having a significant *dA/dN* slope without 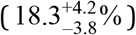 compared to with 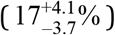 controlling for choice, because the latter method necessarily decreases statistical power. **(H)** Performance vs. counts as predicted by the *fdbk* model (solid lines), compared to the same when the sensory noise (dashed lines) or accumulator-comparison noise (dash-dotted lines) parameters are set to zero. The left plot is for fits to pooled mouse data, the right plot for fits to pooled rat data. **(I)** Relative AIC_c_ likelihood for accumulator models with Gamma-distributed sensory noise (disallows mis-classification of right-for left-side cues), vs. the best model which is *fdbk* (Gaussian-distributed sensory noise). The left plot is for mouse data and the right plot for rat data.

**Figure S6.**
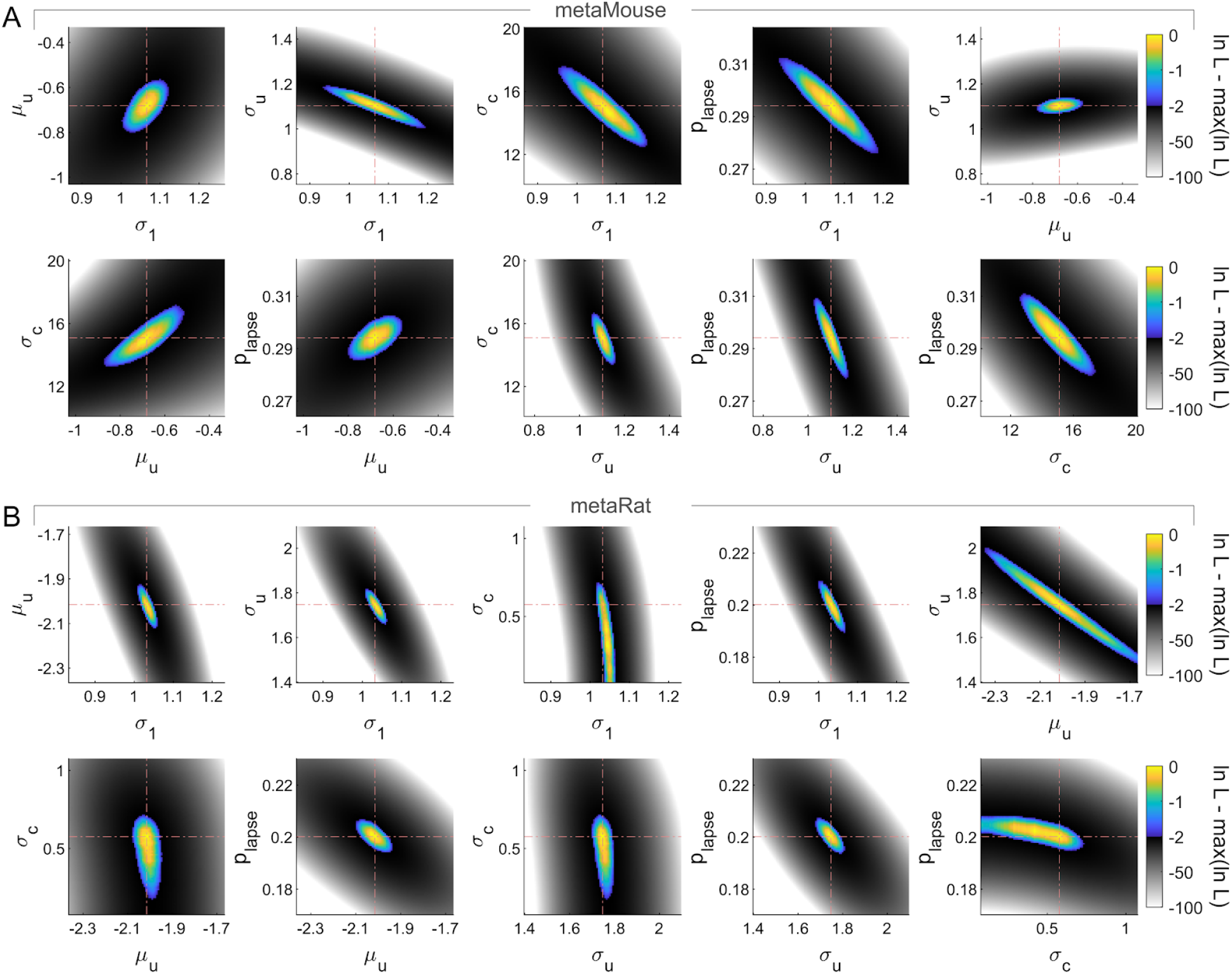
Log-likelihood landscape for the fdbk model vs. all pairs of model parameters. **(A)** *fdbk* model log-likelihood for the pooled mouse data, relative to the maximum (heat map), and for various combinations of parameter values (x-vs. y-axes, different parameter pairs in each plot). All points within the 95% C.I. region are shown in color while points outside are rendered in grayscale. The optimal parameters are indicated by crosshairs. **(B)** Same as **(A)** but for pooled rat data.

**Figure S7.**
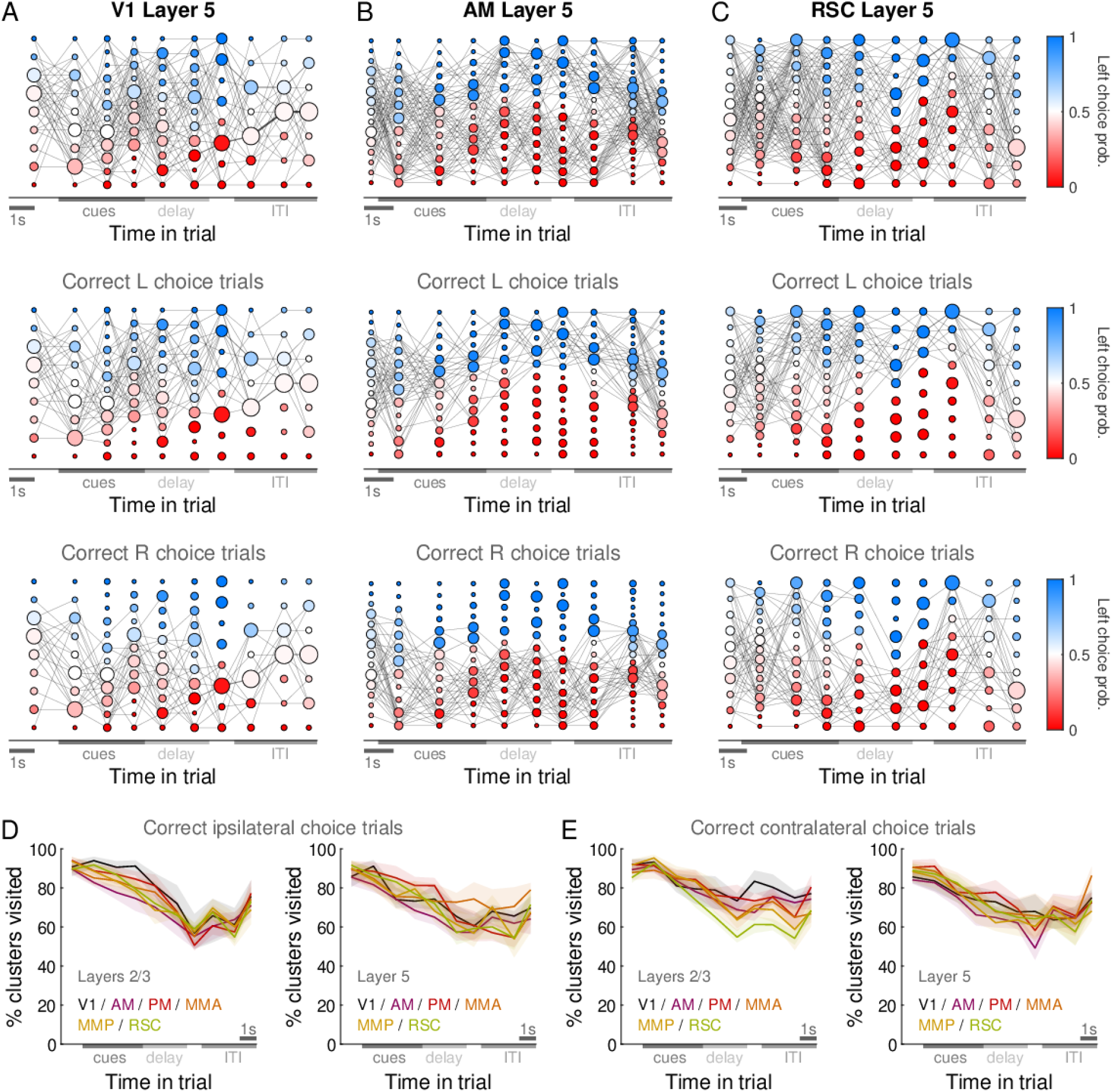
Trial-to-trial variability in trajectories through neural state space. **(A)** Transition probability matrix for neural states through time in the trial (see methods in (Morcos and Harvey 2016)). The neural state space is clustered separately per time-bin in the trial (x-axis), and the resulting clusters depicted as disks with area proportional to the number of trials in which the neural state visited that cluster. Disks are colored by the fraction of trials in which the mouse made a left choice. Lines are drawn between clusters at a given time-bin to clusters at the subsequent time-bin, with intensity proportional to the fraction of trials in which the neural state transitioned from one to the other cluster. The data here is for a single-session recording in layer 5 of V1, same as for Fig. 2. The second and third rows show the transition matrix restricted to correct left-choice and correct right-choice trials respectively. **(B)** Same as **(A)**, but for a single-session recording in layer 5 of AM, same as for Fig. 2. **(C)** Same as **(A)**, but for a single-session recording in layer 5 of RSC, same as for Fig. 2. **(D)** Percent of neural-state clusters at a given time-bin, for which there was at least one trial that was a correct ipsilateral-choice trial. For a single session, this would correspond to e.g. the fraction of clusters per column in the middle row of **(A)** that had one or more edges. Lines: Mean across datasets. Bands: S.E.M. **(E)** Same as **(D)**, but for contralateral-choice trials.

